# Chromosome-level genome assemblies of two hemichordates provide new insights into deuterostome origin and chromosome evolution

**DOI:** 10.1101/2023.11.01.564642

**Authors:** Che-Yi Lin, Ferdinand Marlétaz, Alberto Pérez-Posada, Pedro Manuel Martínez García, Siegfried Schloissnig, Paul Peluso, Greg T. Conception, Paul Bump, Yi-Chih Chen, Cindy Chou, Ching-Yi Lin, Tzu-Pei Fan, Chang-Tai Tsai, José Luis Gómez Skarmeta, Juan J. Tena, Christopher J. Lowe, David R. Rank, Daniel S. Rokhsar, Jr-Kai Yu, Yi-Hsien Su

## Abstract

Deuterostomes are an animal superphylum that includes Hemichordata and Echinodermata (together Ambulacraria) and Chordata. The diversity of deuterostome body plans has made it challenging to reconstruct their ancestral condition and to decipher the genetic changes that drove the diversification of deuterostome lineages. Here, we generate chromosome-level genome assemblies of two hemichordate species, *Ptychodera flava* and *Schizocardium californicum*, and use comparative genomic approaches to infer the chromosomal architecture of the deuterostome common ancestor and delineate lineage-specific chromosomal modifications. We show that hemichordate chromosomes (1*N*=23) exhibit remarkable chromosome-scale macrosynteny when compared to other deuterostomes, and can be derived from 24 deuterostome ancestral linkage groups (ALGs). These deuterostome ALGs in turn match previously inferred bilaterian ALGs, consistent with a relatively short transition from the last common bilaterian ancestor to the origin of deuterostomes. Based on this deuterostome ALG complement, we deduced chromosomal rearrangement events that occurred in different lineages. For example, a fusion-with-mixing event produced an Ambulacraria-specific ALG that subsequently split into two chromosomes in extant hemichordates, while this homologous ALG further fused with another chromosome in sea urchins. Orthologous genes distributed in these rearranged chromosomes are enriched for functions in various developmental processes. We found that the deeply conserved Hox clusters are located in highly rearranged chromosomes but have lower densities of transposable elements within the clusters. We also provide evidence that the deuterostome-specific pharyngeal gene cluster was established via the combination of three pre-assembled microsyntenic blocks. We suggest that since chromosomal rearrangement events and formation of new gene clusters may change the regulatory controls of developmental genes, these events may have contributed to the evolution of diverse body plans among deuterostomes.

## Introduction

The evolutionary events that gave rise to the diverse body plans of deuterostomes remains one of the major mysteries in biology. It is widely accepted that the Deuterostomia superphylum includes Echinodermata, Hemichordata and Chordata, as these animals are characterized by several unique developmental and morphological features, including radial cleavage, deuterostomy, enterocoely formation of the mesoderm, mesoderm-derived skeletal tissues, and pharyngeal openings/slits^1, 2^. Despite these common characters, the different deuterostome lineages have evolved distinct body plans. Chordates are defined by their dorsal tubular central nervous system, notochord and segmented somites^3^, while echinoderms evolved a pentaradially symmetrical adult body, calcitic endoskeleton and a water vascular system^4^, and hemichordates are characterized by a tripartite body organization, which includes a proboscis, collar and trunk^5^. Molecular phylogenetic analyses have supported a sister group relationship between Echinodermata and Hemichordata, forming a clade called Ambulacraria^6, 7^ (Fig. 1a). While subsequent phylogenomic studies have reinforced support for the ambulacrarian clade, some have suggested a sister group relationship between Ambulacraria and Xenacoelomorpha (a group of marine worms lacking definitive coeloms) and even questioned the monophyletic grouping of the Deuterostomia^8–11^. Due to the long evolutionary history of deuterostome lineages and the difficulties in assigning definitive stem fossils during the early diversification of the group, it remains challenging to postulate the ancestral condition of their common ancestor, let alone to decipher the genomic basis underlying the origins of diverse body plans and phylogenetic affiliations. To address these issues, it is helpful to reconstruct the ancestral genome architectures at major nodes of the animal tree using species that occupy key phylogenetic positions, and trace the subsequent evolutionary trajectories along each lineage.

**Fig. 1.**
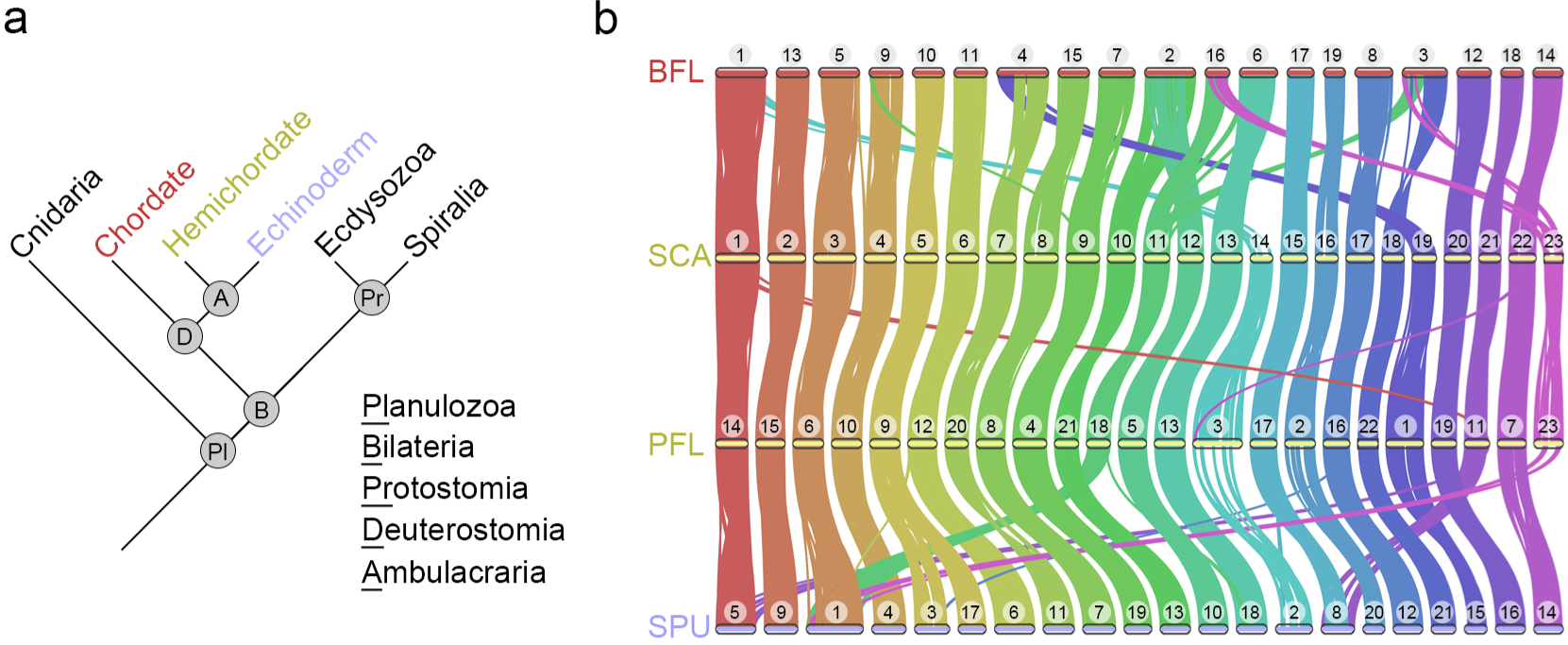
Highly conserved macrosyntenic structure among deuterostomes was detected based on chromosome-level genome assemblies, including two new hemichordate genomes. **a**, A simplified phylogenetic tree of major branches in Planulozoa (Cnidaria+Bilateria). **b**, Macrosynteny conservation among deuterostome species including *Branchiostoma floridae* (BFL), *Ptychodera flava* (PFL), *Schizocardium californicum* (SCA) and *Strongylocentrotus purpuratus* (SPU). Horizontal bars with numbers above represent chromosomes of each species. The conserved synteny blocks between two species are connected by curve lines (minimum of 4 gene pairs within a maximum distance of 75 genes between two matches).

Comparison of diverse metazoan genomes has revealed extensive conservation of chromosome-scale linkage (i.e., “macrosynteny”) across animals^12–15^ and enabled the reconstruction of ancestral chromosome-scale units (chromosomes or chromosome arms)^16–20^. These reconstructions have been used to identify shared and derived synteny patterns that can help to resolve long-standing evolutionary questions, infer lineage-specific chromosomal rearrangements, and clarify animal phylogenetic relationships that have been difficult to resolve using conventional phylogenetic approaches^17–22^. For example, identifications of synapomorphic traits of chromosomal fusion-with-mixing events among sponge, cnidarian and bilaterian genomes provide strong evidence to support the hypothesis that ctenophores are the sister group to all other animals^17^.

Among deuterostomes, vertebrates show extensive genomic duplications^19^, but comparisons of sea urchin with other bilaterians^18^, and analysis of sub-chromosomal assemblies of hemichordates^14^ (1) implied that the chromosomes of the deuterostome ancestor retained the 24 bilaterian ancestral linkage groups, and (2) identified subsequent rearrangement in the sea urchin and chordate lineages^18^. Assembling a complete picture of deuterostome genome evolution, however, requires comparisons including chromosome-scale assemblies of hemichordates. Analyses of karyotype evolution including all deuterostome phylum-level lineages could yield important insights into deuterostome ancestry and the evolution of their diverse body plans.

In this study, we generated chromosome-level genome assemblies for two distantly related hemichordates, the ptychoderid *Ptychodera flava* and spengelid *Schizocardium californicum*. Our comparative genomic analysis showed remarkable macro-syntenic conservation among deuterostome species. Based on the principle of parsimony and comparative analyses with outgroups, we deduced that the last common ancestor of deuterostomes possessed 24 ALGs that match the bilaterian ALGs as previously proposed^18^. We also discovered lineage-specific rearrangements that reflect the temporal progression towards the chromosomal architectures of extant deuterostomes. While our phylogenetic analysis using synteny-based characters supports a monophyletic deuterostome grouping, we did not identify shared derived macrochromosomal rearrangements that distinguish deuterostomes from other bilaterians. Our results confirm that the genomic architectures of deuterostomes retain more ancestral traits than those of protostomes, consistent with a very short evolutionary distance from the last common ancestor of bilaterians to the origin of deuterostomes. Our study thus provides a roadmap for understanding chromosomal evolution and contributes to deciphering the possible developmental genetic changes underlying the emergence of diverse body plans in deuterostomes.

## Results and discussion

### Chromosome-level genome assemblies of two hemichordates

Deuterostomes are composed of three major phyla, including hemichordates, echinoderms and chordates, with the former two constituting a group called Ambulacraria (Fig. 1a). Previous short read-based genome sequencing of two hemichordate species, *Saccoglossus kowalevskii* and *Ptychodera flava*, provided a cornerstone for studies on deuterostome evolution^14^. The fragmented nature of these genome assemblies, however, limits our understanding of chromosome evolution among deuterostome lineages. To address this issue, we employed PacBio long-read and Hi-C technologies to sequence genomes of two enteropneust hemichordates *P. flava* (PFL) and *Schizocardium californicum* (SCA) (Supplementary Fig. 1). The long read-based genome assemblies of PFL and SCA consist of 1.16 Gbp and 0.93 Gbp, respectively (Supplementary Fig. 1). After consideration of HiC contacts (Supplementary Fig. 2), 23 chromosome-scale scaffolds were obtained for both genomes, which matches the 2*N*=46 karyotype of PFL^14^. Protein-coding genes were annotated in the two genome assemblies using transcriptome data and *ab initio* prediction approaches, resulting in 35,856 (PFL) and 27,463 (SCA) annotated genes with high BUSCO scores (Supplementary Fig. 1). Therefore, these two hemichordate genome assemblies reached chromosome level with high completeness in gene annotation.

The 23 chromosomes of the two hemichordate species generally exhibit a one-to-one correspondence based on pairwise comparisons of the positions of orthologous genes (Fig. 1b and Supplementary Fig. 3a). This correspondence further supports the chromosomal-scale accuracy of the independently conducted genome assemblies, since conserved syntnies are unlikely to be generated spuriously by assembly errors. Extending this analysis to sea urchin (*Strongylocentrotus purpuratus*, SPU) and amphioxus (*Branchiostoma floridae*, BFL), which are representative echinoderm and chordate species, we confirmed chromosome-scale syntenic conservation (macrosynteny) among deuterostomes (Fig. 1b and Supplementary Fig. 3b). Given that macro-syntenic conservation has been used to reconstruct ancestral genome architectures and identify lineage-specific chromosomal rearrangement events^18, 19^, we broadened the synteny analysis by including additional species within and outside the deuterostome superphylum. This approach allowed us to confirmthe genomic architecture of the last common ancestor (LCA) of deuterostomes and explore how it evolved among deuterostome lineages.

### Reduction of chromosome numbers during deuterostome evolution

To reconstruct the ancestral chromosomal architectures at key phylogenetic nodes in deuterostomes and investigate the evolutionary history of chromosomal changes, we carried out pairwise genome comparisons of multiple deuterostomes (Supplementary Fig. 4). To identify orthologous chromosomes between species in an unbiased fashion, we employed Fisher’s exact test with Bonferroni correction and risk difference to designate chromosome pairs containing orthologous genes (see Methods). Following refs. 18 and 19, we reasoned that the syntenic units that are conserved between genomes are most likely descended from a common ancestral linkage group (ALG) in the LCA of the two species under investigation. We used the scallop (*Patinopecten yessoensis*, PYE) genome as an outgroup (Supplementary Fig. 5) due to its slow evolution compared with other protostomes^23^ and previously demonstrated conserved syntenies with other animals^18^. Using this comparative approach we inferred ancestral chromosomal architectures at major nodes of the deuterostome phylogeny.

In order to reconstruct the ambulacrarian ancestral chromosomes, we compared the hemichordate PFL genome with the genomes of two echinoderm species (sea urchin SPU and sea star *Pisaster ochraceus*, POC), with the amphioxus or scallop genome serving as an outgroup (Supplementary Fig. 6-9). The dot plot between hemichordate (PFL) and sea urchin (SPU) showed 17 one-to-one corresponding chromosomes (Supplementary Fig. 4a), suggesting that (1) these chromosome pairs are homologous and (2) the LCA of PFL and SPU (i.e., the ambulacrarian LCA) already possessed these 17 ALGs. We also identified several one-to-two and one-to-three corresponding chromosomes between PFL and SPU, implying that large-scale chromosomal rearrangement events occurred after the lineages diverged from the ambulacrarian LCA. We polarized the direction of chromosomal change, and identified the likely ancestral state by comparing to the outgroup species. For example, *P. flava* PFL11 and PFL17 together correspond to *S. purpuratus* SPU8 (Supplementary Fig. 8d), implying that either PFL11 and PFL17 arose by a split of an ancestral ambulacrarian chromosome or SPU8 arose by the fusion of two ancestral chromosomes. Comparison with amphioxus chromosomes, however, showed that PFL11 and PFL17 respectively correspond to amphioxus BFL18 and BFL17 (Supplementary Fig. 8g), indicating that these two chromosome pairs evolved from two distinct ALGs in the deuterostome LCA. Based on the parsimony principle, we reasoned that hemichordates inherited the two ALGs directly as PFL11 and PFL17, while sea urchin SPU8 was fused from the two distinct ancestral chromosomes, as also noted in ref. 14 using a different sea urchin species *Lytechinus variegatus* (Supplementary Fig. 8a). By reiterating such comparisons (Supplementary Fig. 6-9), we find that the LCA of deuterostomes possessed 24 ALGs (DALGs). Importantly, these 24 DALGs correspond to the 24 bilaterian ALGs (BALGs) deduced by Simakov et al.^18^, confirming that the deuterostome LCA and the bilaterian LCA possessed very similar chromosomal architectures. Our notation for the deuterostome ALGs therefore follow those of the bilaterian ALGs^18^. Among the 24 DALGs, nine remain intact in all five deuterostome species we investigated, while fifteen have undergone lineage-specific changes (Fig. 2).

**Fig. 2.**
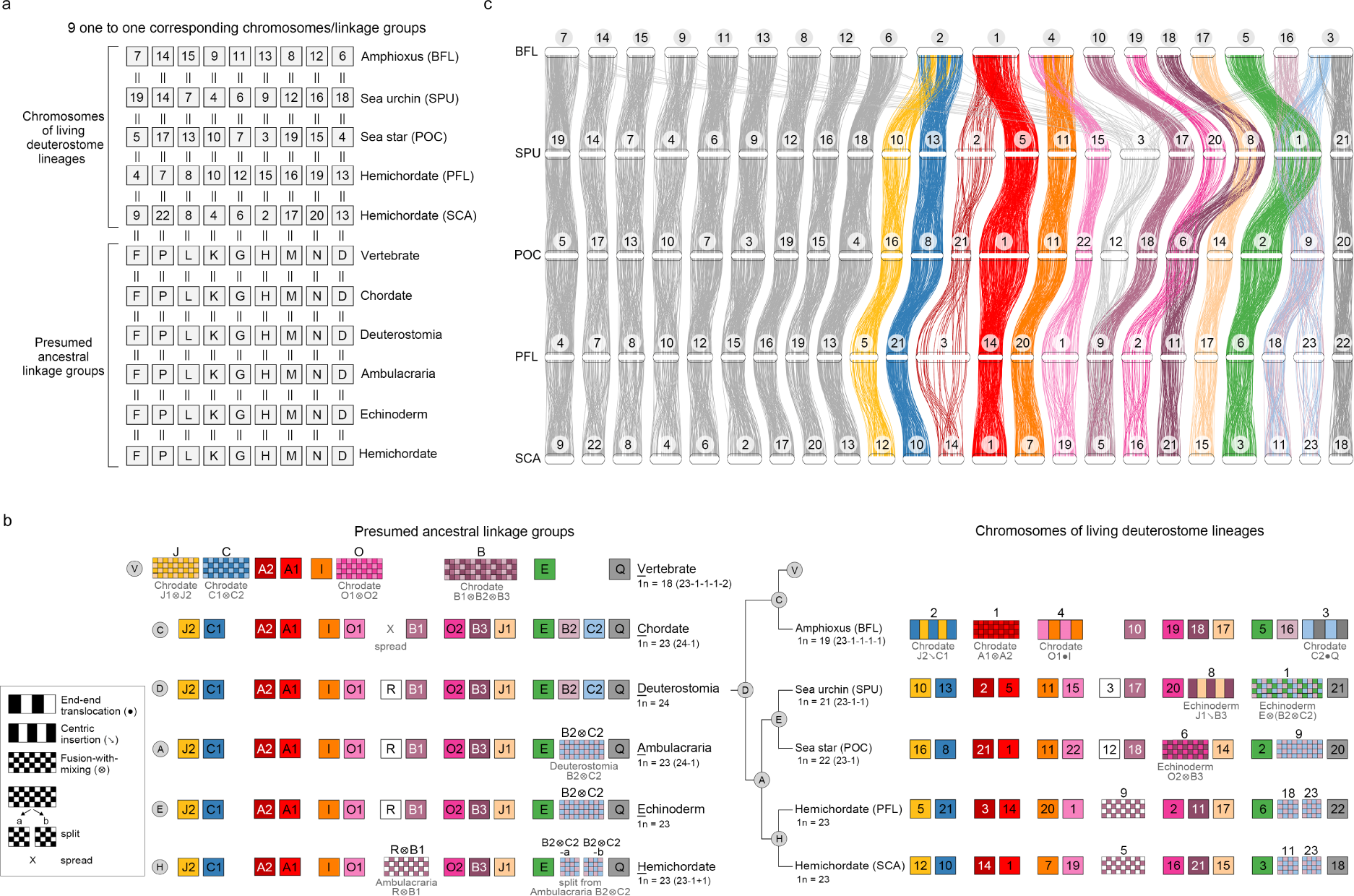
Evolutionary history of deuterostome chromosome architectures. **a-b**, A schematic representation of chromosome evolution in deuterostome lineages. The chromosomal architectures of presumed LCAs (bottom in **a** and left in **b**) and the chromosomal architectures of living deuterostome species (top in **a** and right in **b**). Each box denotes an individual chromosome. Haploid number (1*N*) and increase (+) or decrease (-) in quantity of chromosomes are indicated. The color code of boxes is taken from the previous study on vertebrate ancestral chromosomes, except for the nine one-to-one corresponding chromosomes (**a**, light gray boxes). Chromosomal architecture of the LCA of vertebrates was based on the previous study^19^. In cases where chromosomal fusion events were deduced, types of changes are indicated below color boxes with symbols defined previously^18^; end-end translocation (●), centric insertion (↘) and fusion-with-mixing (⊗). Box sizes do not reflect the actual sizes of chromosomes. **c**, Chromosomal positions of the orthologous gene pairs among five deuterostome species. Horizontal bars with numbers on top represent chromosomes of each species. In total, 3668 orthologous gene pairs are illustrated. For ease of comparison, the chromosome sizes are scaled proportionally such that the five genome assemblies reach equal sizes. Except for the genes that spread into multiple chromosomes in amphioxus (BFL), gene pairs that are not located on the corresponding chromosomal pairs or cannot be found in all five species are not shown.

Figure 2 illustrates chromosomal rearrangement events with color boxes: interspersed boxes represent chromosomal fusions followed by translocations, while checkerboards depict chromosomal fusions followed by extensive mixing, which is a common feature of deep chromosome evolution^18^ (Fig. 2); rearrangements were determined based on pairwise conserved syntenies between target species (Supplementary Fig. 6-9). These illustrations correspond to the chromosomal rearrangement events defined by Simakov et al.^18^, with algebraic symbols indicating end-end fusion (●), centric insertion (↘) and fusion-with-mixing (⊗)^18^. Notably, four interspersed boxes correspond to end-end fusions and five correspond to centric insertions followed by chromosomal translocations (e.g., BFL4 and BFL2 in Fig. 2b). From the 24 DALGs, we inferred that the numbers of chromosomes were reduced in a lineage-specific manner. In the lineage leading to ambulacrarians (node A in Fig. 2b), DALGs B2 and C2 fused and mixed extensively to become the ambulacrarian ALG B2⊗C2, while other DALGs remained relatively intact, resulting in 23 ambulacrarian ALGs (AALGs). In the hemichordate lineage (node H in Fig. 2b), AALG B2⊗C2 split into two chromosomes (B2⊗C2-a and B2⊗C2-b), while AALGs R and B1 fused and mixed (AALG R⊗B1) to become a single chromosome (PFL9 and SCA5, respectively), resulting in 1*N*=23 chromosomes in both hemichordate species. The split of the AALG B2⊗C2 can be understood as a possible Robertsonian (i.e., centric) fission in which a presumably metacentric chromosome is transformed into two acrocentrics. Whether the shared chromosomal linkages of PFL and SCA represents the ancestral hemichordate state can only be determined by analysis of pterobranch hemichordate genomes, but it is clear from the pairwise comparison (Supplementary Fig. 3a) that no macrosyntenic changes have occurred since the last common ancestor of PFL and SCA, which lived more than 370 mya^14^.

Similarly, the echinoderm LCA (node E in Fig. 2b) likely possessed 23 ALGs (EALGs), with the same chromosomal architecture as the ambulacrarian LCA; subsequently, different fusion events occurred in the sea star and sea urchin lineages. In the sea star, EALGs O2 and B3 fused (O2⊗B3) and evolved into POC6, resulting in a 1*N*=22 karyotype. In the sea urchin *S. purpuratus*, SPU8 chromosome arose through the fusion of EALGs J1 and B3, via central insertion (J1↘B3), while SPU1 arose by fusion with extensively mixing from EALGs E and B2⊗C2, denoted as E⊗(B2⊗C2), resulting in 1*N*=21 chromosomes. The three-way fusion E⊗(B2⊗C2), is also shared by *Lytechinus variegatus*^18^ and *Paracentrotus lividus*^24^, and is therefore likely a shared derived character of the superorder Echinacea, a hypothesis that can be tested by sequencing other members of this group.

In the chordate lineage (node C in Fig. 2b), orthologous genes located on DALG R were dispersed into many chromosomes^18^, leading to 23 chordate ALGs (CALGs). This dispersion was inferred from the observation that no particular amphioxus chromosomes show significant enrichment of syntenic blocks corresponding to DALG R-derived chromosomes in echinoderms (SPU3 or POC12, Fig. 2c and Supplementary Fig. 4e-f). Similarly, no concentration of R was found in vertebrates or ascidians^18^. In the amphioxus *B. floridae*, four chromosomal fusion events occurred (J2↘C1, A1⊗A2, O1●I and C2●Q), reducing the number of chromosomes to 1*N*=19^19^. The inferred chordate-specific chromosomal dispersion and the four chromosomal fusion events in amphioxus BFL are consistent with previous findings^18^. One of these fusion events (A1⊗A2) was also observed in the sea urchin *Paracentrotus lividus*^24^, suggesting that A1 and A2 were arms of a metacentric chromosome that fused independently in urchin and amphioxus. From the 23 chordate ALGs, previous studies^18, 19^ deduced that the lineage leading to vertebrates had undergone four chromosomal fusion events (J1⊗J2, C1⊗C2, O1⊗O2 and B1⊗B2⊗B3), reducing the 23 CALGs to 18 vertebrate ALGs. These chromosomal rearrangement events and the evolutionary history of genomic architectures among deuterostomes are summarized in Figure 2.

### Stepwise changes in chromosomal architectures within the sea urchin lineage

We expect that chromosomal fusion-with-mixing events would occur in a stepwise process as evolution proceeds. As such, two distinct chromosomes (at t_0_) would fuse (at t_1_), either by end-end fusion or centric insertion, and this event would be followed by rounds of intrachromosomal inversions and translocations (at t_2_) until the fused chromosome became scrambled (at t_s_) (as illustrated in Supplementary Fig. 10). We therefore postulate that comparing chromosome architectures between species with a relatively short divergence time should allow us to identify the evolutionary state of individual chromosomes during this stepwise process. We thus analyzed two additional sea urchin species, *L. variegatus* (LVA) and *L. pictus* (LPI), for which chromosomal-level genome assemblies are available for syntenic comparison^15, 25^. LVA and LPI are within the genus *Lytechinus*, which share a common ancestor with *S. purpuratus* 50 million years ago (mya)^26^. By analyzing syntenic conservation of these three sea urchin species (Supplementary Fig. 11), we inferred that their LCA (tentatively assumed to be sea urchin LCA) possessed 21 ALGs (SALGs) due to two shared chromosomal fusion events, J1↘B3 and E⊗(B2⊗C2) (node S in Supplementary Fig. 10). These two fusions were also observed in the recently decoded sea urchin *P. lividus* genome^24^, indicating a common genomic trait of currently available sea urchin genomes. We also deduced 20 ALGs (LALGs) in the *Lytechinus* LCA, owing to a *Lytechinus*-specific chromosomal fusion event (G●D) (node L in Supplementary Fig. 10). Descending from the *Lytechinus* LCA, *L. variegatus* and *L. pictus* each underwent a distinct chromosomal fusion event, F●(J1⊗B3) into *L. variegatus* LVA1 and F●C1 into *L. pictus* LPI5), independently resulting in 1*N*=19 chromosomes for both species.

Based on the phylogenetic relationships and deduced chromosomal architectures (Supplementary Fig. 10), we construct a putative history of several chromosomal fusion events. For example, two echinoderm ALGs (J1 and B3 at t_0_) fused via centric insertion after which a translocation event resulted in the sea urchin ALG J1↘B3 (at t_2_). This chromosome then underwent extensive recombinations to become the *Lytechinus* ALG J1⊗B3 (at t_s_). In the lineage leading to *L. variegatus*, but not *L. pictus*, end-end fusion of *Lytechinus* ALGs F and J1⊗B3 resulted in the extant LVA1 chromosome (at t_1_). Within the LVA1 chromosome, we observed no obvious translocation between regions descended from LALGs J1⊗B3 and F, suggesting that the end-end fusion likely occurred recently in the lineage leading to *L. variegatus*. In *L. pictus*, chromosome LPI5 was derived from end-end fusion of LALGs F and C1 followed by a translocation event. Intriguingly, the independent, species-specific fusion event of the two *Lytechinus* species involved the same chromosome (LALG F). Such recent chromosomal fusions may alter recombination rate and cause reproductive isolation, as observed during nematode speciation^27^. Together, the fusion events in sea urchins clearly illustrate how stepwise changes may occur in chromosomal architectures.

In several fusion-with-mixing cases, we did not observe transitional states (e.g., SALG E⊗(B2⊗C2) resulted from EALGs E and B2⊗C2, Supplementary Fig. 10), implying that these fusion events occurred at a relatively ancient time. Assuming that intrachromosomal rearrangements occurred at a constant rate, we postulate the order of fusion events based on synteny patterns. For example, in comparison with the centric insertion pattern of SALG J1↘B3, SALG E⊗(B2⊗C2) exhibits fusion-with-mixing, suggesting that the fusion of EALGs E and B2⊗C2 occurred earlier than that of EALGs J1 and B3. Therefore, from the echinoderm LCA that possessed 23 ALGs to the sea urchin LCA (or more specifically, the LCA of the three sea urchin species under investigation) that contained 21 ALGs, there may have been a transitional state with 1*N*=22 chromosomes, when EALGs E and B2⊗C2 were already fused but J1 and B3 remained separated. Intriguingly, it has been reported that the haploid genomes of *Cidaris cidaris* and *Arbacia punctulata*, which respectively belong to an early branching sea urchin group and an euechinoid outgroup of *Lytechinus* and *S. purpuratus*, each contain 22 chromosomes^28, 29^, suggesting that only one fusion event occurred in early branching sea urchins. Thus, we hypothesize that EALGs E and B2⊗C2 fused before the divergence of the sister subclasses of sea urchins, cidaroids and euechinoids, at least 268 mya^30^. The second fusion event, involving EALGs J1 and B3, possibly occurred later, after the emergence of *Arbacia* and before the divergence of *Lytechinus* and *S. purpuratus* (i.e., between ∼185 and 50 mya)^31^. If that is the case, the LCA of all living sea urchins would have possessed 1*N*=22 chromosomes, instead of the presumed 21 ancestral chromosomes illustrated in Supplementary Figure 10. Future synteny analyses and chromosomal architecture reconstructions using genomes of early branching sea urchins will help to resolve this question.

### Lineage-specific chromosomal fusion events in major animal groups

To understand whether the deuterostome chromosomal architectures differ from those of protostomes, we extended our analysis to include several recently published chromosome-level genome assemblies of protostomes. Consistent with previous observations^16, 23^, we found that the chromosomes of most protostome species are highly rearranged. Nevertheless, we were able to identify genomes of five spiralian species^23, 32–35^, including three bivalves (two clam species, *Ruditapes philippinarum* and *Sinonovacula constricta*, and the aforementioned scallop *P. yessoensis*) and two polychaete annelids (*Paraescarpia echinospica* and *Streblospio benedicti*), which are more conserved and comparable to the presumed bilaterian ALGs and extant deuterostome genomes. Our syntenic analysis shows that all the five spiralian species share four specific fusion-with-mixing events (Supplementary Fig. 12-13), as predicted previously based on four syntenic synapopmorphies of spiralians identified using different datasets^18, 36^. Comparisons of six chromosome-scale ecdysozoan genomes, however, showed that that they are highly reorganized relative to the bilaterian ancestor^18^, making it difficult to reconstruct the chromosomal architecture of their LCA. The four spiralian fusions, however, are clearly absent in ecdysozoan, consistent with their status as spiralian syntenic synapomorphies^18^. For example, these four fusion events are clearly absent in two butterfly genomes (Supplementary Fig. 14). Based on these pairwise syntenic comparisons, we inferred that the LCA of protostomes most likely also possessed 24 ALGs that correspond to the 24 bilaterian ALGs (Supplementary Fig. 12). This correspondence suggests that the genomic architecture of the deuterostome LCA and protostome LCA did not undergo large-scale inter-chromosomal fusions when they initially diverged from the bilaterian LCA. However, during subsequent evolution, protostome lineages appear to have accumulated much more extensive changes in their chromosomal architectures than deuterostome lineages.

After chromosomal fusion with extensive mixing, it is unlikely that genes in a fused chromosome would be sorted to reassemble back into individual chromosomes with the original makeup^17, 18^, and, such irreversible chromosomal fusion-with-mixing events can be used as polarized traits for probing deep phylogenetic relationships of animals^17, 18^. Recent phylogenomic studies have provided evidence to support the sister-group relationship between Ambulacraria and Xenacoelomorpha, and some even questioned the monophyletic grouping of Deuterostomia^8–11^ (Fig. 3a). We thus sought to test whether the identified chromosomal fusion-with-mixing traits could help to resolve this issue. We first recorded all chromosomal rearrangement events as category data, which was then converted into a binary matrix (Fig. 3b and c). Bayesian phylogenetic and clustering analyses based on these synteny-based characters united the deuterostomes (Fig. 3d-e). Regarding derived chromosomal changes, we identified an ambulacrarian-specific chromosomal fusion (B2⊗C2), a chordate-specific chromosomal dispersion^18^ (originated from ALG R) and four spiralian-specific chromosomal fusion events^18^ (L⊗J2, O2⊗K, Q⊗H and O1⊗R) (Supplementary Fig. 16). Notably, among the five ambulacrarian and chordate species we analyzed, no common chromosomal fusion (i.e., synapomorphy) was identified, despite the fact that all the species retain the same ancestral traits (nine one-to-one corresponding chromosomes). We also noted that the bilaterian chromosomal rearrangement events were not observed in the jellyfish (*Rhopilema esculentum*, RES) genome^18^ (Supplementary Fig. 15-16). Therefore, the five major extant animal groups (i.e., ambulacrarians, chordates, spiralians, ecdysozoans and cnidarians) do not share common derived traits in terms of inter-chromosomal rearrangement events, and the observed chromosomal fusion events appear to have occurred after the divergence of these major animal groups. Such chromosomal fusion-with-mixing events may help to resolve the phylogenetic positions of animal groups with weak phylogenetic signals based on sequence divergence. For example, Xenacoelomorpha, a group comprising xenoturbellids and acoelomorphs, have been placed as early branching bilaterians (Nephrozoa hypothesis) or as a sister group of ambulacrarians (Xenambulacraria hypothesis)^10, 11, 37, 38^. To find support for either hypothesis, we examined the recently available chromosome-level genome assembly of *Xenoturbella bocki*^39^, which belongs to xenoturbellids. We found no evidence of the ambulacraria-specific chromosomal fusion (B2⊗C2) in the *X. bocki* genome. Therefore, this fusion event is truly specific to ambulacrarians and does not provide evidence supporting either hypothesis. Overall, our results reinforce the idea that the branch length between bilaterian LCA and deuterostome LCA is likely very short^8^, and our analyses also show that deuterostome lineages experienced fewer chromosomal fusion events than protostomes during evolution.

**Fig. 3.**
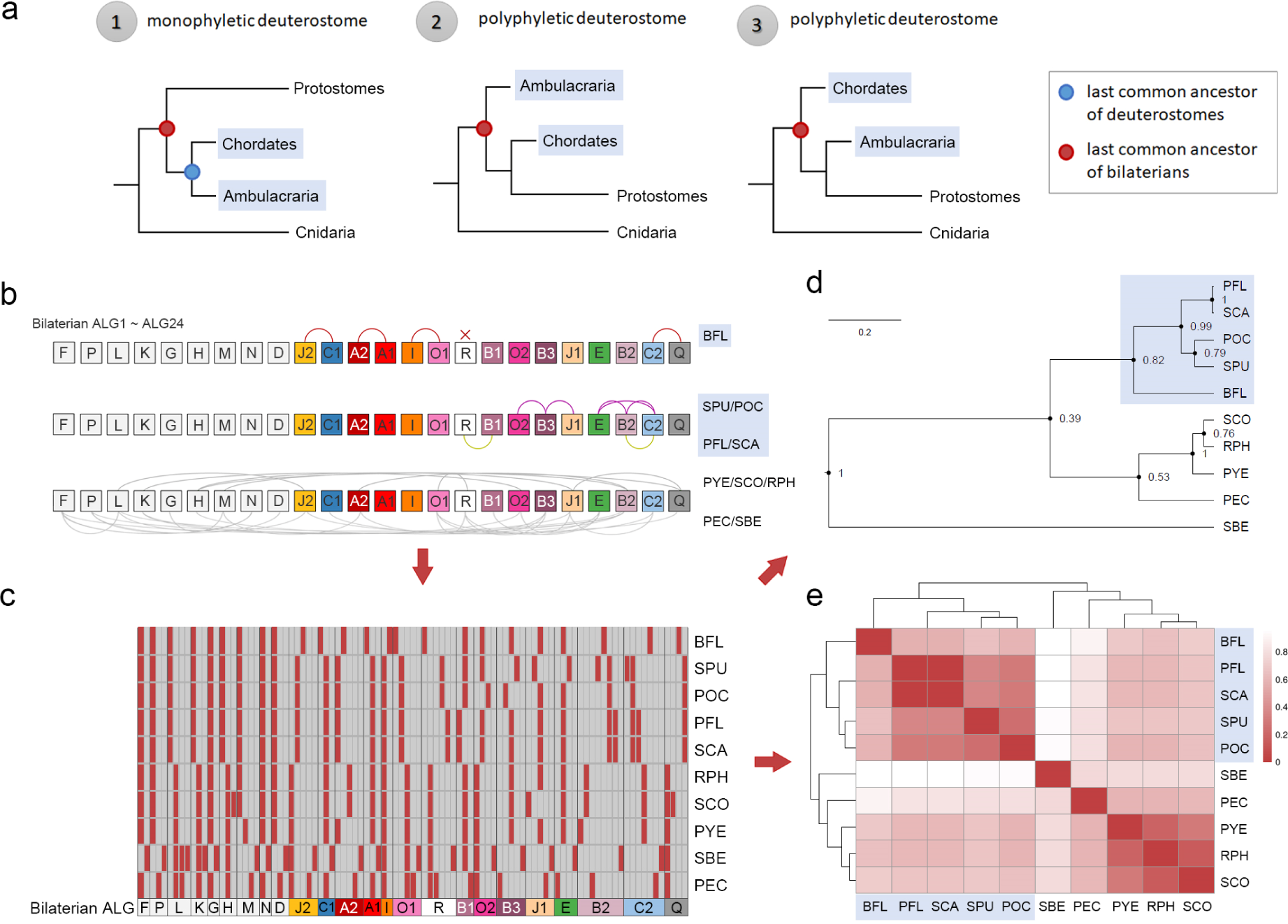
Category clustering analysis based on rearrangement events of the 24 bilaterian ancestral chromosomes. **a**. Three scenarios of phylogenetic relationships among bilaterians can be postulated. In the first scenario (monophyletic deuterostome), two deuterostome branches, chordates and ambulacrarians, are grouped together, and their LCA (the LCA of deuterostomes) is denoted with a blue dot. The LCA of bilaterians is indicated with a red dot. In the other two scenarios, one of the deuterostome branches is grouped with protostomes, resulting in polyphyletic deuterostomes. In these latter two scenarios, the LCA of deuterostomes and bilaterians is the same. **b**. Distinct chromosomal rearrangement events of each species, including fusion, split and spread events, are recorded into the category data based on changes deviated from the 1*N*=24 bilaterian ancestral linkage groups (BALGs). For example, there are three categories for the bilaterian ALG K, including (1) no rearrangement event (BFL, SPU, POC, SCA, PFL), (2) O2⊗K (RPH, SCO, PYE, PEC) and (3) J1⊗(O2⊗K) (SBE). **c**. Conversion of the category data into a binary data matrix. Dark vertical lines distinguish different chromosomes. Red box denotes the chromosomal status of each species as compared to the BALGs. For the ten species that we examined, the number of the chromosomal rearrangement categories ranges from 2 (BALGs F, G, N and I) to 8 (BALG B2). A detailed binary code table is provided in Supplementary Data 1. **d-e**. Bayesian phylogenetic analysis (d) and clustering analysis (e) based on the binary data shows that the five deuterostome species (shaded in blue) are grouped together.

### GO enrichment analyses of lineage-specific chromosomal rearrangement events

We presume that chromosomal fusion-with-mixing would likely disrupt long-range enhancers and/or topological association domains (TADs) to cause changes in gene regulation. Therefore, the particular genes located in chromosomes that underwent lineage-specific fusions may provide hints as to the origins of lineage-specific novelties. To understand potential biological consequences of specific chromosomal changes in deuterostome species, we performed gene ontology (GO) enrichment analyses on genes located on the corresponding chromosomes of extant deuterostomes. The ambulacraria-specific chromosomal fusion-with-mixing resulted in the inferred AALG B2⊗C2, which has remained as a single chromosome POC9 in the sea star (Fig. 2b and Fig. 4). We found that genes located in POC9 are enriched in several GO terms related to development, including germ layer formation, neural development, axial patterning, gastrulation and regulation of BMP and Wnt signaling pathways (Fig. 4a and Supplementary Fig. 17e). This observation suggests that in the lineage leading to ambulacrarians, many developmental regulatory genes would have experienced considerable shuffling in their relative positions via chromosomal fusion-with-mixing (B2⊗C2), which may have caused significant changes in their expression patterns. The fused AALG B2⊗C2 further underwent distinct chromosomal fusion and splitting events in sea urchins and hemichordates, respectively (Fig. 2b and Fig. 4). In all the three sea urchin genomes we analyzed, a single chromosome (e.g., SPU1) was derived from the fusion of EALGs E and B2⊗C2 (Supplementary Fig. 10). GO analysis revealed that genes related to development are also enriched in SPU1 (Fig. 4b and Supplementary Fig. 18e). Intriguingly, genes involved in bone and otolith development are also enriched in this sea urchin-specific fusion chromosome. Further analysis on genomes of other sea urchin species and functional experiments will be required to determine whether the rearrangement of these genes is related to the emergence of the unique skeletogenic lineage of sea urchins. In both hemichordate species, we inferred that two chromosomes (PFL18 and PFL23 of *P. flava* and SCA11 and SCA23 of *S. californicum*) were split from the fused AALG B2⊗C2, resulting in HALGs B2⊗C2-a and B2⊗C2-b in the LCA of hemichordates (Fig. 2b). GO enrichment analysis revealed that genes located on PFL18 (descendant of either HALG B2⊗C2-a or B2⊗C2-b) are enriched in biological processes associated with immune response and chemotaxis, suggesting that distinct interactions with environmental factors could have emerged during hemichordate evolution (Fig. 4c and Supplementary Fig. 19c). Additional lineage-specific fusion events observed in deuterostomes include the echinoderm O2⊗B3 and J1↘B3 (resulting in the sea star POC6 and the sea urchin SPU8, respectively) and the hemichordate-specific fusion R⊗B1 (corresponding to PFL9 and SCA5) (Fig. 2b). The top GO terms enriched in POC6, SPU8 and PFL9 include neuronal regulation, thyroid hormone transport and germ cell migration, respectively (Fig. 4d and Supplementary Fig. 17-19).

**Fig. 4.**
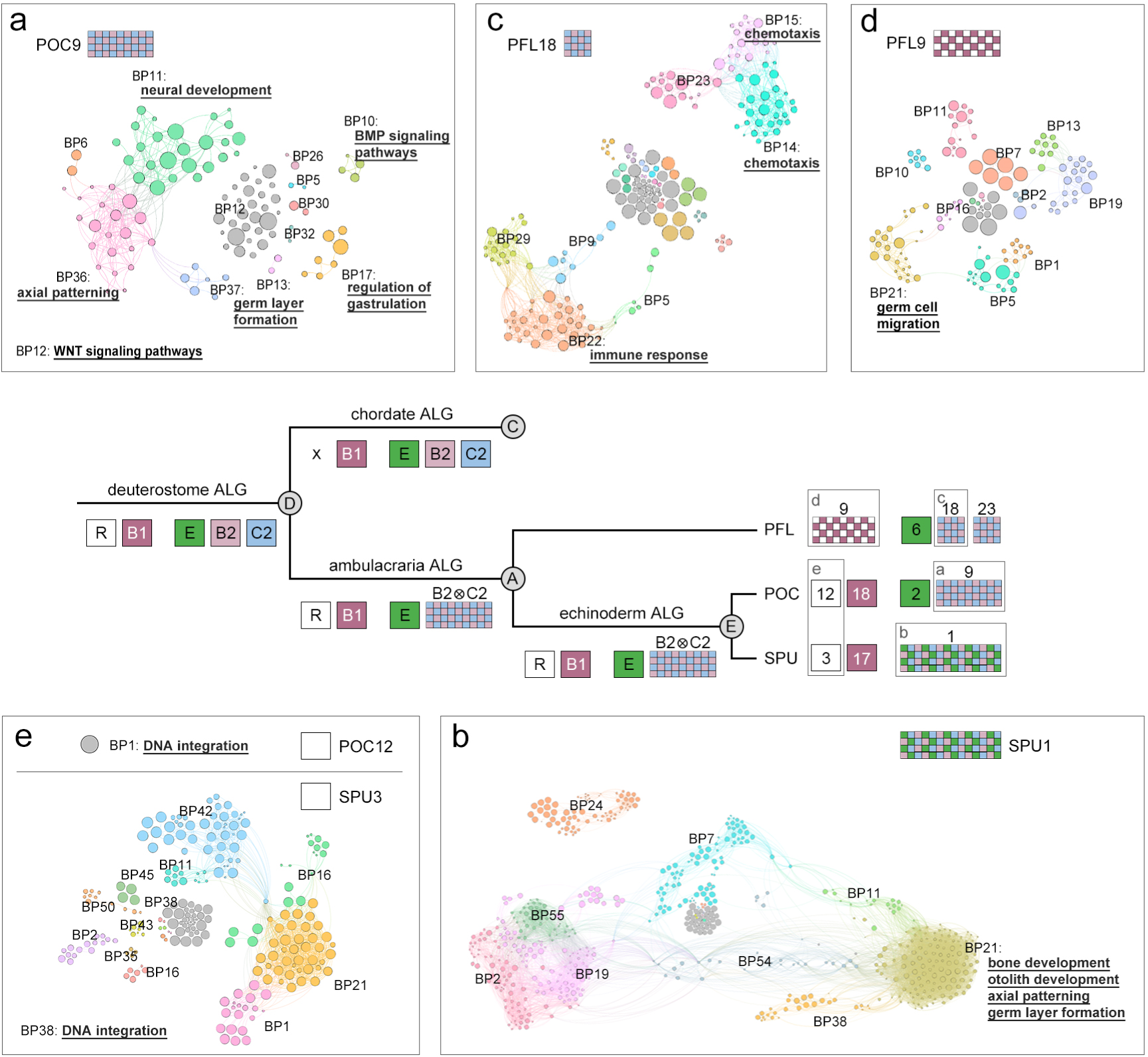
Gene ontology (GO) enrichment analyses of chromosomes that underwent lineage-specific changes in deuterostomes. GO enrichment analysis of genes located in the sea star POC9 (**a**), sea urchin SPU1 (**b**), hemichordate PFL18 (**c**) and PFL9 (**d**). The echinoderm chromosomes (POC12 and SPU3) corresponding to deuterostome DALG R were also analyzed to understand the chordate-specific chromosomal dispersion (**e**). The enriched GO terms (adjusted *p*-value < 0.1) are clustered and divided into different modules, and selected terms are underlined. The top three enriched biological process (BP) GO terms of each module are shown in Supplementary Figs. 17-19.

One common feature of chordate chromosomal rearrangements is the dispersion of deuterostome ALG R (Fig. 2b-c). To assess the potential biological consequences of this chordate-specific event, we performed GO enrichment analysis on the corresponding ambulacrarian chromosomes (POC12 and SPU3), which have remained as individual chromosomes. Intriguingly, we found that genes involved in DNA integration, including several transposase genes, are enriched in both POC12 and SPU3 (Fig. 4e and Supplementary Fig. 17a and 18a). This result suggests that the dispersion of DALG R in the chordate lineage could have been due to the misregulation of transposase genes. Taken together, our GO enrichment analyses provide a global view of possible regulatory and functional changes related to the lineage-specific chromosomal rearrangements. At least some of these potential changes are likely associated with the evolution of distinct lineage-specific features and diverse body plans in deuterostomes.

### Hox clusters in rearranged chromosomes and devoid of transposable elements

Hox genes are often arranged in clusters and specify bilaterian body regions along the anteroposterior axis^40^. Contrary to their functional conservation, we noted that Hox clusters are located in chromosomes that underwent fusion with extensive mixing among the ten bilaterian species we examined, with the sole exception of amphioxus BFL (Supplementary Fig. 16). In the LCA of bilaterians, the Hox cluster was inferred to be positioned in BALG B2. The descendant of this ALG (DALG B2) contributed to the ambulacraria-specific fusion with DALG C2 to form AALG B2⊗C2. Subsequently, its descendant in echinoderms further underwent an additional fusion-with-mixing with ALG E to give rise to SPU1 in sea urchins. Meanwhile, in hemichordates, AALG B2⊗C2 split into HALGs B2⊗C2-a and B2⊗C2-b (represented by the extant PFL18 and PFL23, Supplementary Fig. 16). Intriguingly, this splitting event in hemichordates separated the Hox cluster and the *distalless* gene, which are commonly linked in vertebrate genomes^41^. This genetic feature appears to be unique to hemichordates, as the Hox cluster and *distalless* gene are linked and located in the same chromosome in all other deuterostome species we examined (i.e., amphioxus BFL16, sea star POC9 and sea urchin SPU1). Nevertheless, it remains unclear whether the separation of the Hox cluster and *distalless* gene during the hemichordate-specific chromosomal split would have resulted in functional consequences related to the origin of the hemichordate body plan. BALG B2 is also involved in different fusion-with-mixing events in the five spiralian species, with the spiralian Hox clusters located on the highly rearranged RPH14, SCO9, PYE1, PEC4 and SBE9 (Supplementary Fig. 16). It is tempting to speculate that these chromosome rearrangement events may have changed the regulatory landscape of Hox genes and contributed to the evolution of lineage-specific body plans. Further studies would certainly be required to test this hypothesis.

While intrachromosomal rearrangement events are highly associated with the accumulation of transposable elements (TEs)^42, 43^, Hox clusters are known to be largely devoid of TEs in chordates^44, 45^. The exclusion of TEs from Hox clusters is thought to be chordate-specific, as this trend was not detected in five protostome species that have been analyzed (including four insects and the nematode *Caenorhabditis elegans*)^45^. The observation that most Hox clusters are situated in chromosomes that underwent fusion-with-mixing prompted us to analyze TE densities in the Hox-bearing chromosomes. By analyzing the distributions of different TEs, including DNA transposons (DNA), long terminal repeats (LTR), long interspersed nuclear elements (LINE) and short interspersed nuclear elements (SINE), we observed a clear drop-off of TE densities within Hox clusters as compared with the non-Hox regions of the same chromosomes; this trend was observed in all ten bilaterian species we examined (Fig. 5a and Supplementary Fig. 20-22). Of note, the overall TE densities in Hox-bearing chromosomes were similar to the densities in the entire genomes (Supplementary Fig. 23). The exclusion of TEs in Hox clusters is particularly apparent in amphioxus BFL and hemichordate PFL (∼77% less than the density of non-Hox regions) in which the Hox clusters are relatively intact (Fig. 5). Therefore, the trend of lower TE densities in Hox clusters can be widely observed in bilaterians and is not limited to chordates. Thus, the mechanism that prevents TE invasion appears to remain functional in Hox clusters even though these regions are situated in highly rearranged chromosomes. Moreover, we noticed that many genes neighboring Hox clusters, except for the *evx* genes, are highly rearranged and their orthologous genes are commonly found in different chromosomes (Supplementary Fig. 24-25). This result is consistent with the observation that TEs exist at higher densities outside of Hox clusters, where they can promote intrachromosomal rearrangements. Further characterizations of TE distributions within Hox clusters revealed a slightly higher density of TEs around the posterior Hox genes (between Hox9 and Hox15) within the amphioxus BFL Hox cluster. This higher density is consistent with a previous observation of repeat islands between the amphioxus posterior Hox genes that may contribute to the highly derived posterior region of the amphioxus Hox cluster^44, 46, 47^. Despite the generally low TE density across the Hox cluster of hemichordate PFL, we noticed that the inversion of *Hox13b* and *Hox13c* coincides with the presence of more TEs near the posterior end of the Hox cluster (Fig. 5b, PFL). Similarly, the numbers, positions and orientations of Hox genes between *Hox5* and *Hox11/13* in the three sea urchin species (SPU, LVA, and LPI) have undergone notable changes, which is in line with the higher densities of TEs detected in these regions (Fig. 5b). Taken together, these results indicate that exclusion of TEs from Hox clusters appears to be a conserved feature in bilaterians. Nevertheless, TE invasions sometimes occur in the posterior regions of deuterostome Hox clusters, and these invasions have likely contributed to local rearrangements of Hox genes. Our observations are reminiscent of the proposed “deuterostome posterior flexibility” model, which explains how the posterior Hox genes evolved faster in deuterostomes than in protostomes^47, 48^. In conclusion, the distributions of TEs both outside and within certain regions of Hox clusters coincide with intrachromosomal gene rearrangements, which likely lead to modifications in TAD structures of Hox clusters and thus the transcriptional regulation of Hox genes.

**Fig. 5.**
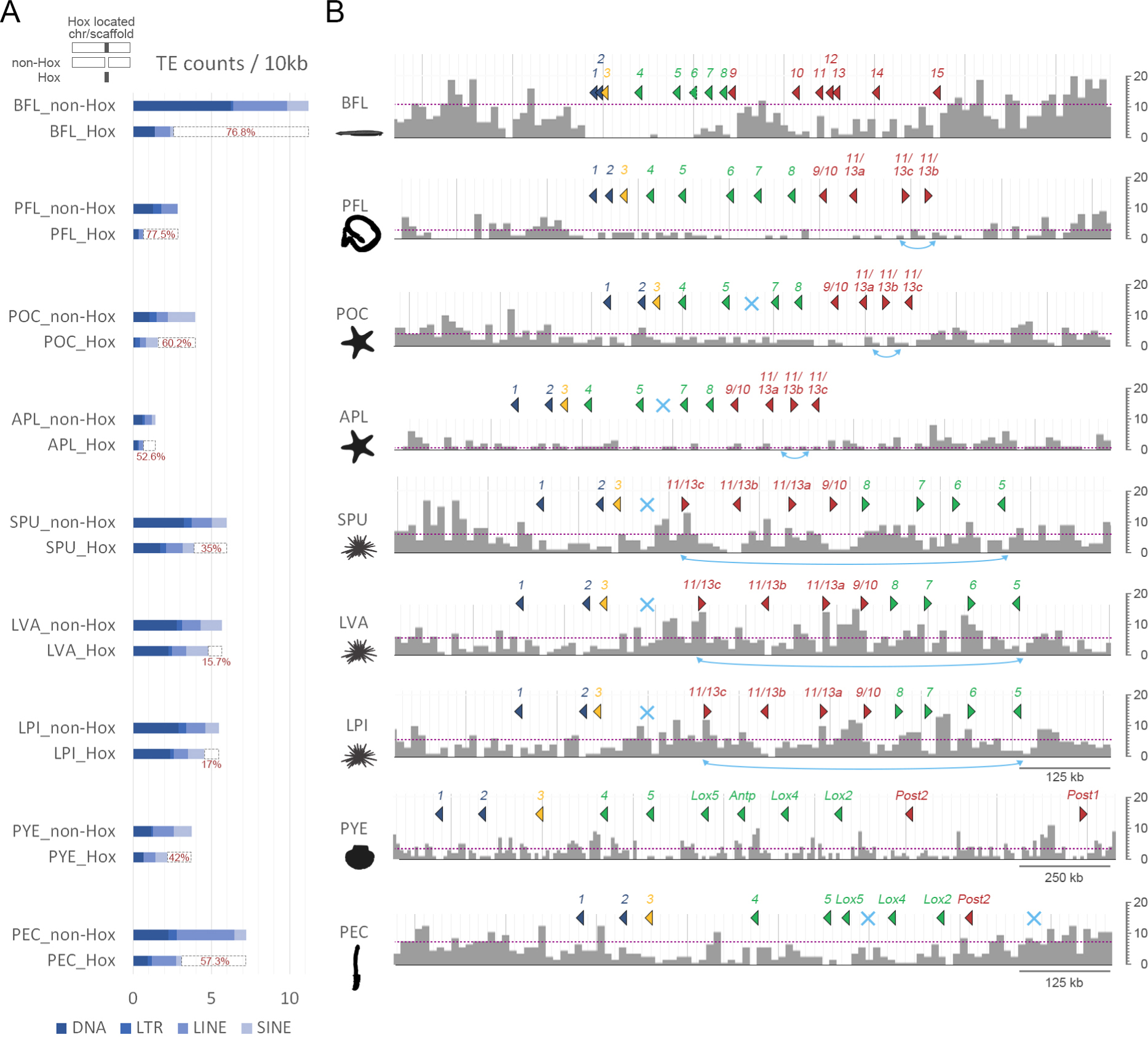
Counts of transposable elements (TEs) around the Hox-bearing genomic regions. **a**. Densities of all TEs (DNA + LTR + LINE + SINE) within the Hox cluster region and non-Hox region of the Hox-bearing chromosome/scaffold of each species. The percent differences in normalized TEs counts between the non-Hox region and the Hox cluster region are illustrated (dashed bars and red values). **b**. Distributions of all TEs around the Hox gene cluster of each species. The bin size for each histogram is 10,000 bp. Dotted lines indicate the averaged TE densities of the Hox-bearing chromosomes/scaffolds. Color coding denotes division of Hox genes in “anterior” (dark blue), “group 3” (yellow), “middle” (green) and “posterior” (red) groups. The light blue crosses represent missing Hox genes. Double arrows in light blue indicate inversion events of Hox genes.

### Evolutionary history of the pharyngeal gene cluster

The pharyngeal gene cluster contains four transcription factor genes (in the order of *nkx2.1*, *nkx2.2*, *pax1/9* and *foxa*) and two non-transcription factor genes (*slc25a21a* and *mipol1*), and their expression in the pharyngeal slits and surrounding endoderm is considered to be a deuterostome-specific feature^14^. Three additional genes, *msx*, *cnga* and *egln3*, which respectively encode a homeobox transcription factor, a subunit of cyclic nucleotide-gated channels and Egl-9 family hypoxia inducible factor 3, are also linked to the cluster in some deuterostome species^8, 14, 49^. The complete pharyngeal gene cluster has so far only been found in deuterostomes, but some of the genes are also linked in protostomes^8^. It has thus been proposed that rather than being a deuterostome-specific trait, an intact cluster may have already been present in the LCA of bilaterians and was later dispersed in protostome lineages^8^. To gain insights into the evolutionary history of the pharyngeal cluster, we analyzed gene complements of the cluster in several bilaterian and non-bilaterian genomes (Fig. 6a and b). In all the deuterostome genomes we analyzed, we found that *xrn2*, which encodes a 5’ to 3’ exoribonuclease, is associated with the aforementioned pharyngeal genes and usually located upstream and adjacent to *nkx2.1*. Based on the gene repertoire and linkage relationships in the deuterostomes genomes, we deduced that the complete complement of the pharyngeal cluster in the LCA of deuterostomes included ten genes. The complement began with *xrn2*, followed by three transcription factor genes (*nkx2.1*, *nkx2.2* and *msx*), then *cnga*, *pax1/9*, *slc25a21*, *mipol1* and *foxa*, and it ended with *egln3*. Several lineage-specific changes then took place within the pharyngeal clusters of deuterostomes (Fig. 6b and Supplementary Fig. 26). In the hemichordate PFL, *cnga* was duplicated, and *ghrA* genes invaded the pharyngeal cluster between the *cnga* and *pax1/9* genes. In the sea urchin SPU, the pharyngeal cluster is broken into three parts, although the three parts are still located on the same chromosome (SPU1), and the second part (including *msx*, *cnga*, *pax1/9* and *slc25a21*) is inverted.

**Fig. 6.**
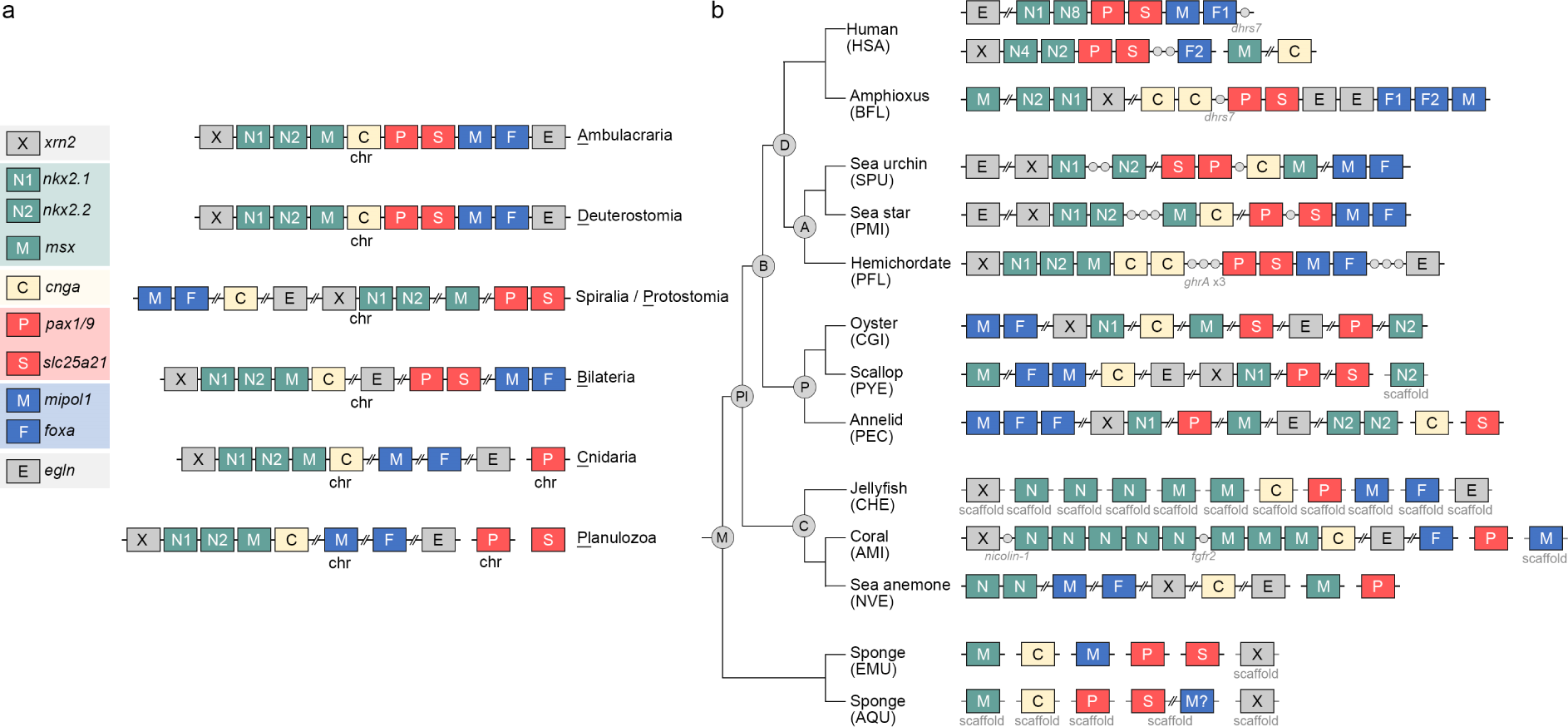
A possible evolutionary history of the pharyngeal gene cluster. The pharyngeal gene architectures are shown for presumed last common ancestors at key phylogenetic nodes (**a**) and selected living metazoan species (**b**) (see Supplementary Fig. 21 for the complete dataset). Genes that are commonly linked together are shown in the same color; homeobox-containing genes, including *nkx2.1*, *nkx2.2* and *msx*, are in green, *mipol1* and *foxa* genes are in blue, and *pax1/9* and *slc25a21* are in red. The gray circles indicate genes that are located within the pharyngeal gene cluster. Double slashes are introduced when more than three genes are located in between two pharyngeal genes. Because *pax* genes of cnidarians and sponges do not show one-to-one correspondence with those of bilaterians, we surveyed the locations of all potential *pax* genes and found that none is linked with the other pharyngeal-related genes in cnidarians and sponges.

In all six spirarian genomes we analyzed, orthologs of *xrn2* were found to be adjacent to *nkx2.1*, and *mipol1* and *foxa* genes were also linked (Fig. 6b and Supplementary Fig. 26). In a previous study^8^, paired gene linkages of *nkx2.1* and *nkx2.2*, *pax1/9* and *slc25a21*, and *mipol1* and *foxa* were also identified in various protostomes. These results support the existence of three microsyntenic blocks, including (1) *xrn2* and *nkx2* genes, (2) *pax1/9* and *slc25a21*, and (3) *mipol1* and *foxa*, as conserved features of protostomes. Intriguingly, most of the orthologous genes of the pharyngeal cluster are located on the same chromosome, regardless of whether the microsyntenic relationships are maintained. Based on these observations, we proposed two scenarios for the evolution of the pharyngeal gene cluster: (1) the LCA of bilaterians (similar to the LCA of deuterostomes) possessed a complete pharyngeal gene cluster that later broke up into three microsyntenic blocks in protostomes; (2) the LCA of bilaterians (similar to the LCA of protostomes) had the pharyngeal genes arranged in three microsyntenic blocks in the same chromosome and became closely linked as a compact cluster in deuterostomes. To find evidence supporting or excluding these scenarios, we analyzed the genomic positions of the orthologous genes in nonbilaterians, including several cnidarians and sponges (Supplementary Fig. 26). In the coral AMI, we observed a syntenic block containing *xrn2*, *nkx2*, *msx-*related and *cgna* genes. Other cnidarian species either had preserved parts of this syntenic block (e.g., *xrn2* and *nkx2* are adjacent in the coral XSP; *msx-*related and *cgna* are linked in the sea anemone SCA) or they lacked the syntenic relationships (Supplementary Fig. 26). Additionally, in all six cnidarian genomes we analyzed, *slc25a21* was absent. This gene was likely lost in cnidarians, because an ortholog of *slc25a21* was identified in the sponge genomes. Moreover, except for the *pax* genes, orthologs of the other pharyngeal genes are located on the same chromosome of most cnidarian genomes we analyzed. In the two sponge genomes, orthologs of the pharyngeal genes are mostly located on different chromosomes or scaffolds, and no microsyntenic blocks were identified. Based on the parsimony theory, we can infer that one microsyntenic block (composed of *xrn2*, *nkx2*, *msx-*related and *cgna* genes) was already present in the LCA of bilaterians and cnidarians, and the other pharyngeal genes were located on the same chromosomes but had not yet formed microsyntenic blocks. We therefore suspect that in the LCA of bilaterians, the two additional microsyntenic relationships (*pax1/9-slc25a21* and *mipol1-foxa*) were established. In the lineage leading to the examined spiralian species, the more ancient syntenic block was likely partially disrupted, with only *xrn2* and *nkx2* genes remaining tightly associated. During the evolution of deuterostomes, the three microsyntenic blocks became linked and the *egln* gene was added at the end, forming the complete pharyngeal gene cluster. This hypothetical process agrees with the second scenario in which the compact pharyngeal gene cluster in deuterostomes was gradually established. We consider the first scenario to be less likely, because although the individual pharyngeal genes were mostly located on the same chromosome, the genes would have needed to be assembled into an ordered cluster in the bilaterian ancestor before breaking into three blocks in protostomes. Assembly of the three microsyntenic blocks into the deuterostome pharyngeal gene cluster likely contributes to the co-regulation of the genes. Indeed, analysis of the temporal expression profiles using available transcriptomic datasets support the idea that clustering of the pharyngeal genes likely contributes to their co-regulation^50^.

## Conclusions

In this study, we generated chromosome-level genome assemblies for two hemichordate species. The hemichordate chromosomes (1*N*=23) exhibit remarkable chromosome-scale macrosynteny when compared to other deuterostomes, including several echinoderm and chordate species. This high level of conservation allows us to infer that the LCA of deuterostomes possessed 24 ALGs, the same complement as inferred for the bilaterian ancestor. We further deduced lineage-specific chromosomal rearrangement events that resulted in reduced numbers of chromosomes during deuterostome evolution. Genes distributed in chromosomes that underwent lineage-specific fusions are enriched for functions in developmental processes, immune responses and chemotaxis. Changes to the regulatory control of these genes may be related to the evolution of distinct lineage-specific features in deuterostome lineages. One example of this concept is the deeply conserved Hox cluster, which is commonly situated in a chromosome that is highly rearranged. Nevertheless, Hox genes in deuterostomes generally remain tightly linked with the posterior Hox genes showing higher flexibility, consistent with the distribution pattern of TEs within the Hox clusters.

Another conserved gene cluster, the deuterostome pharyngeal gene cluster, appears to have been established gradually by combining three pre-assembled microsyntenic blocks present in the LCA of bilaterians. Complete clustering likely contributes to the co-regulation of the pharyngeal genes. In summary, these results showcase how the global view provided by comparative genomics can contribute to our understanding of genome evolution. Moreover, the lineage-specific genomic changes identified herein may help to delineate molecular mechanisms driving the evolution of the diverse body plans of deuterostomes.

## Methods

### Sample preparation and sequencing

High molecular weight (HMW) genomic DNA of *Ptychodera flava* (PFL) was extracted using DNAzol (ThermoFisher Scientific) from the sperm of a single male individual collected from Penghu Islands, Taiwan. The size of the purified HMW genomic DNA was examined using a pulsed-field gel electrophoresis system (BIO-RAD). The genomic DNA was then sequenced by the Dresden Genome Center using the PacBio platform with 60x coverage. For *Schizocardium californicum* (SCA), HMW DNA was extracted from a ripe male Schizocardium. To keep secretion of mucus to a minimum, animals were washed several times and kept in ice cold seawater during the sperm extraction process. Male spermaducts were opened with forceps and sperm was pipetted with a glass pasteur pipette and transferred to an Eppendorf tube. Tubes were spun down and excess seawater was removed before being placed on ice. The DNA extraction protocol was adapted from Stefanik et al. 2013^51^ with a combination of pouring between Eppendorf tubes instead of pipetting and avoiding any vortexing. The genomic DNA was then sequenced using the PacBio platform with 63x coverage.

### Chromosome-level genome assembly

For PFL, the initial genome assembly was generated using MARVEL assembler^52^ with PacBio reads. Purge Haplotigs (version 1.1.0)^53^ was used to phase the diploid genome assembly onto the haploid assembly. The phased haploid genome assembly was then scaffolded using HiRISE with a HiC library (Dovetail Genomics). The sequences of the genome assemblies were further curated using Pilon (version 1.23.2)^54^ with the Illumina short reads. For SCA, the raw read error correction, read trimming and assembly were performed with the Canu assembler (version 1.5)^55^. Canu was configured to run with a genomeSize parameter set at approximately 1.8GBp or roughly twice the expected genome size due to high heterozygosity. After assembly, two rounds of polishing were performed with the Arrow consensus calling algorithm^56^. The completeness of the polished genome assemblies was evaluated by using BUSCO (version 5.1.2)^57^ with the dataset metazoa_odb10, which contains 954 BUSCO gene groups.

### Gene prediction and functional annotation

For *Ptychodera flava*, gene models were predicted using a combination of *ab initio* gene prediction, homology support and transcriptome sequencing. First, *ab initio* gene prediction was conducted using the MAKER2 pipeline^58^ (Dovetail Genomics). Second, the protein sequences from other species, including mouse, chick, zebrafish, spotted gar, sea lamprey, amphioxus, ascidian, sea urchin and sea anemone, were aligned to the PFL genome assembly using GeMoMa (version 1.7)^59^. Third, the Illumina RNA-seq short reads from PFL at 16 stages^50^ were mapped using STAR aligner (version 2.7.6a)^60^. The subsequent genome-guided transcript reconstruction was conducted with StringTie (version 2.1.4)^61^ and CLASS2 (version 2.1.7)^62^. The transcripts were also assembled *de novo* using Trinity (version 2.11.0)^63^ and then mapped to the genome assembly by minimap2 aligner (version 2.17-r941)^64^. Fourth, the full-length transcripts were generated with PacBio technology (Iso-seq), and IsoSeq3 (version 3.3.0, https://github.com/PacificBiosciences/IsoSeq) was used to cluster the IsoSeq transcripts. LoRDEC (version 0.9)^65^ was used to curate the Isoseq transcripts with the Illumina RNA-seq short reads. The polished IsoSeq transcripts were then mapped to genome assembly using minimap2. Gene models based on Iso-seq data were then reconstructed with cDNA_Cupcake (version 9.1.1, https://github.com/Magdoll/cDNA_Cupcake). Finally, the reconstructed transcripts from the different shreds of evidence were merged and filtered by EvidenceModeler (version 1.1.1)^66^. The combined gene models were further updated by PASA (version 2.4.1)^67^. The amino acid sequences were predicted from the transcripts using TransDecoder (version 5.5.0, https://github.com/TransDecoder/TransDecoder). Each amino acid sequence was aligned against NCBI metazoa subset of the nr database using Blast2GO/OmicsBox (version 1.3.11)^68^ with blastp-fast for gene description. The GO (gene ontology) term for each gene was annotated using Blast2GO/OmicsBox^68–70^.

For *S. californicum*, gene prediction was performed as in Marlétaz et al. 2023^24^. Briefly, hints for *de novo* prediction using Augustus^71^ were derived from transcriptome and protein alignments. Particularly, proteins from *S. kowalevskii* were aligned using Exonerate (version 2.2.0)^72^. A custom repeat library was constructed and annotated using Repeatmodeler and subsequently used to mask repeated regions in the *S. californicum* genome using Repeatmasker (v.4.0.7, http://www.repeatmasker.org). We filtered out gene models that extensively overlapped with mobile elements. Isoforms and UTR regions were added using PASA^67^ leveraging the alignment of the assembled transcriptome.

### The genomic datasets for other species

The genome assemblies and gene annotation files across metazoans were collected from public domains, including human *Homo sapiens* (HAS), amphioxus *Branchiostoma floridae* (BFL), sea urchins *Strongylocentrotus purpuratus* (SPU), *Lytechinus pictus* (LPI) and *Lytechinus variegatus* (LVA), sea stars *Patiria miniata* (PMI), *Acanthaster planci* (APL) and *Pisaster ochraceus* (POC), scallop *Patinopecten yessoensis* (PYE), clams *Ruditapes philippinarum* (RPH) and *Sinonovacula constricta* (SCO), oyster *Crassostrea gigas* (CGI), annelids *Streblospio benedicti* (SBE) and *Paraescarpia echinospica* (PEC), argus *Erebia aethiops* (EAE) and *Aricia agestis* (AAG), prawn *Penaeus chinensis* (PCH), horseshoe crabs *Tachypleus tridentatus* (TTR) and *Carcinoscorpius rotundicauda* (CRO), nematode *Heterodera glycines* (HGL), corals *Acropora millepora* (AMI) and *Xenia sp.* (XSP), jellyfish *Rhopilema esculentum* (RES), *Sanderia malayensis* (SMA) and *Clytia hemisphaerica* (CHE), sea anemones *Nematostella vectensis* (NVE) and *Scolanthus callimorphus* (SCA) and sponges *Ephydatia muelleri* (EMU) and *Amphimedon queenslandica* (AQU). Supplementary Table 1 lists the sources and other information on the genome data used in this study. The Braker2 pipeline (version 2.1.6)^73–79^, including GeneMark (version 3.62)^80^ and AUGUSTUS (version 3.4.0)^71^, was used for gene prediction for genomes lacking gene model annotations.

### Genome comparison

Pairwise syntenic comparisons between species were conducted using MCscan (Python version) of JCVI (version 1.0.9)^81, 82^. The jcvi.compara.catalog module with the LAST aligner of MCscan was used to identify orthologous gene pairs between two species. The parameter C-score was set to 0.99 for filtering the LAST hit to contain the reciprocal best hit. The minimum number of gene pairs in a cluster was set to 1 without a restricted window size. The synteny dot plots were visualized using jcvi.graphics.dotplot module. Chromosomes used in the syntenic comparison were labeled with an abbreviation of the species names and ordered according to size (BFL, PFL, SCA, SPU, POC, PYE, RPH, SBE and PEC) or the existing names (LPI, LVA, EAE, AAG, PCH, TTR, CRO, HGL, SCO and RES).

To assign corresponding chromosome pairs between species, Fisher’s exact test with Bonferroni correction in the R software environment (version 3.6.3) was used to calculate the quantitative significance of orthologs located on the chromosome pairs. Risk difference was used to judge significantly higher or lower than others. For example, in Supplementary Fig. 3a, the number of ortholog pairs in PFL1 and SPU15 is 202 (a); in PFL1 and non-SPU15 it is 99 (b); in non-PFL1 and SPU15 it is 45 (c) and in nonPFL1 and SPU15 it is 8528 (d). These four numbers were subjected to the Fisher’s exact test. The significance levels of all chromosomal pairs were examined; the Bonferroni correction was used for multiple-comparisons. Subsequently, the risk difference was calculated as a/(a+b) – c/(c+d). The criterion for corresponding chromosome pairs between two species was an adjusted *p*-value smaller than 1E-10 and a risk difference value greater than 0. Adjusted *p*-values between 1E-2 and 1E-10 with positive risk differences were considered to be small-scale chromosomal rearrangement events and are not presented in figures describing the evolutionary history of chromosomal architectures.

Macrosyntenic conservation analysis on the four deuterostome species (BFL, PFL, SCA and SPU) shown in Fig. 1f was visualized using the jcvi.graphics.karyotype module of MCscan. The syntenic block was set to a minimum of 4 gene pairs with a maximum distance of 75 genes between two matches.

### Clustering and Bayesian phylogenetic analyses

Distinct chromosomal rearrangement events of the ten bilaterian species were manually recorded into the category data based on changes deviated from the 1*N*=24 bilaterian ancestral chromosomes (ALGs). The category data was subsequently converted into a binary data matrix (Supplementary Data 1) and visualized by using the heatmap.2 function of the gplots R package (version 3.1.3). Notably, most species have only one category per ALG. However, in some species, an earlier fusion event was also recorded due to the stepwise process during chromosomal evolution. Taking PEC chromosome 2 as an example, the fusion of Protostome ALGs L and J2 occurred, resulting in Spiralian ALG L⊗J2. Subsequently, Spiralian ALGs L⊗J2 and C2 were further fused leading to PEC chromosome 2. As a result, both categories, L⊗J2 and C2⊗(L⊗J2), for PEC were recorded as “1”. The redundant categories were then removed to avoid double counting before clustering analysis. The distance matrix among the ten bilaterian species was then calculated based on the binary data matrix using the dist function with the binary method in R. The clustering result was visualized with the pheatmap R package (version 1.0.12). Bayesian phylogenetic analysis was conducted using BEAST (version 1.10.4)^83^. First, the manually converted NEXUS file of binary code matrix (Supplementary Data 1) was transformed into an XML file using BEAUti with default parameters. After 10000 randomly sampled trees were generated using BEAST, the consensus tree was generated using TreeAnnotator with 25% burnin and visualized using FigTree (https://github.com/rambaut/figtree, version 1.4.4).

### GO enrichment analysis

The gene list for each selected chromosome was subjected to GO enrichment analyses using Blast2GO/OmicsBox (version 1.3.11) with an adjusted *p*-value (FDR) of 0.05. The REVIGO algorithm (http://revigo.irb.hr/)^84^ was then used to remove redundant GO terms based on the semantics. Finally, the enriched GO terms were clustered and visualized by Gephi (version 0.9.5, https://gephi.org/).

### Hox gene cluster

The genome assemblies and gene model files of bilaterians for Hox gene analysis were downloaded from the public domain (Supplementary Table 1). Some misannotated Hox genes were manually curated. Repetitive elements for each species were identified *de novo* using RepeatModeler (version 2.0.1)^85^. RepeatMasker (version 4.1.2-p1, http://www.repeatmasker.org) was then used for searching and quantifying the identified repeats on each genome assembly, including four transposable elements: DNA transposons (DNA), long terminal repeats (LTR), long interspersed nuclear elements (LINE) and short interspersed nuclear elements (SIINE). The numbers of the different transposable elements were calculated with a bin size of 10 kilobases or 50 kilobases using BEDTools (version 2.30.0)^86^ and deepTools (version 3.5.1)^87^. The genome sequences and transposable element tracks were subjected to visualization using a local genome browser, JBrowse (version 1.16.10)^88^. The silhouettes were downloaded from PhyloPic (https://www.phylopic.org/).

### Pharyngeal gene cluster

The genome assemblies across metazoans were collected from the public domain (Supplementary Table 1). For the genome lacking annotations, the Braker2 (version 2.1.6) pipeline, including GeneMark (version 3.62) and AUGUSTUS (version 3.4.0), was used to predict gene models. Protein sequences of known pharyngeal-related genes were used as query sequences to blast the genome assemblies, and the hits were further confirmed by searching the NCBI nr database.

## Acknowledgement

The authors wish to thank the staff at the core facility of the Institute of Cellular and Organismic Biology, and NGS Genomics core facility of the Biodiversity Research Center, Academia Sinica for technical assistance. We appreciate the valuable discussions with Dr. Mey-Yeh Lu. We also thank Marcus Calkins for English editing. This work was supported by grants 112-2326-B-001-004 (Y.H.S.) and 110-2621-B-001-001-MY3 (J.K.Y.) from the National Science and Technology Council, Taiwan, grant AS-GC-111-L01 from Academia Sinica, Taiwan (Y.H.S. and J.K.Y.), and grant PID2019-103921GB-I00 from Ministerio de Economía y Competitividad, Spain (J.J.T.). P.M.M.G. was funded by a postdoctoral fellowship from Junta de Andalucía (DOC_00397).

## Author Contributions

Che Y.L., Y.H.S. and J.K.Y. conceived the study. Genomic DNA of *P. flava* and *S. californicum* was prepared by Y.C.C. and P.B., respectively. S.S. and Che Y.L. generated and annotated the *P. flava* genome assembly. P.P. and D.R.K. performed the library preparation and sequencing of *S. californicum*. G.T.C assembled the *S. californicum* genome. F.M., D.S.R. and C.J.L. annotated and analyzed the *S. californicum* genome assembly. A.P.P., P.M.M.G., J.L.G.S. and J.J.T. analyzed the transcriptome and chromosomal conformation of *P. flava.* Che Y.L. analyzed synteny, clustering, GO enrichment and TEs. Y.C.C., C.C., Ching Y.L., T.P.F. and C.T.T. collected the *P. flava* samples. Y.H.S., Che Y.L., J.K.Y. and D.S.R. wrote the manuscript with input from C.J.L., J.J.T., F.M. and A.P.P.

## Data availability

*P. flava* genome assembly used in this work is publicly available: https://www.ncbi.nlm.nih.gov/bioproject/PRJNA747109. The version described in this paper is version JASXRY010000000.

## Reference

1 Lowe, C. J., Clarke, D. N., Medeiros, D. M., Rokhsar, D. S. & Gerhart, J. The deuterostome context of chordate origins. Nature 520, 456–465 (2015). 10.1038/nature14434

2 Nanglu, K., Cole, S. R., Wright, D. F. & Souto, C. Worms and gills, plates and spines: the evolutionary origins and incredible disparity of deuterostomes revealed by fossils, genes, and development. Biol Rev Camb Philos Soc 98, 316–351 (2023). 10.1111/brv.12908

3. Satoh, N. Chordate Origins and Evolution: The Molecular Evolutionary Road to Vertebrates. Chordate Origins and Evolution: The Molecular Evolutionary Road to Vertebrates, 1–206 (2016).

4 McClay, D. R. Evolutionary crossroads in developmental biology: sea urchins. Development 138, 2639-2648 (2011). 10.1242/dev.048967

5 Rottinger, E. & Lowe, C. J. Evolutionary crossroads in developmental biology: hemichordates. Development 139, 2463–2475 (2012). 10.1242/dev.066712

6 Cannon, J. T. et al. Phylogenomic resolution of the hemichordate and echinoderm clade. Curr Biol 24, 2827–2832 (2014). 10.1016/j.cub.2014.10.016

7 Dunn, C. W., Giribet, G., Edgecombe, G. D. & Hejnol, A. Animal Phylogeny and Its Evolutionary Implications. Annual Review of Ecology, Evolution, and Systematics 45, 371–395 (2014). 10.1146/annurev-ecolsys-120213-091627

8 Kapli, P. et al. Lack of support for Deuterostomia prompts reinterpretation of the first Bilateria. Sci Adv 7 (2021). 10.1126/sciadv.abe2741

9 Marletaz, F. Zoology: Worming into the Origin of Bilaterians. Curr Biol 29, R577–R579 (2019). 10.1016/j.cub.2019.05.006

10 Mulhair, P. O., McCarthy, C. G. P., Siu-Ting, K., Creevey, C. J. & O’Connell, M. J. Filtering artifactual signal increases support for Xenacoelomorpha and Ambulacraria sister relationship in the animal tree of life. Curr Biol 32, 5180-+ (2022). 10.1016/j.cub.2022.10.036

11 Philippe, H. et al. Mitigating Anticipated Effects of Systematic Errors Supports Sister-Group Relationship between Xenacoelomorpha and Ambulacraria. Curr Biol 29, 1818-+ (2019). 10.1016/j.cub.2019.04.009

12 Putnam, N. H. et al. The amphioxus genome and the evolution of the chordate karyotype. Nature 453, 1064–1071 (2008). 10.1038/nature06967

13 Putnam, N. H. et al. Sea anemone genome reveals ancestral eumetazoan gene repertoire and genomic organization. Science 317, 86–94 (2007). 10.1126/science.1139158

14 Simakov, O. et al. Hemichordate genomes and deuterostome origins. Nature 527, 459–465 (2015). 10.1038/nature16150

15 Warner, J. F., Lord, J. W., Schreiter, S. A., Nesbit, K. T., Hamdoun, A. & Lyons, D. C. Chromosomal-Level Genome Assembly of the Painted Sea Urchin Lytechinus pictus: A Genetically Enabled Model System for Cell Biology and Embryonic Development. Genome Biol Evol 13 (2021). 10.1093/gbe/evab061

16 Martin-Duran, J. M. et al. Conservative route to genome compaction in a miniature annelid. Nat Ecol Evol 5, 231–242 (2021). 10.1038/s41559-020-01327-6

17 Schultz, D. T., Haddock, S. H. D., Bredeson, J. V., Green, R. E., Simakov, O. & Rokhsar, D. S. Ancient gene linkages support ctenophores as sister to other animals. Nature 618, 110–117 (2023). 10.1038/s41586-023-05936-6

18 Simakov, O. et al. Deeply conserved synteny and the evolution of metazoan chromosomes. Sci Adv 8, eabi5884 (2022). 10.1126/sciadv.abi5884

19 Simakov, O. et al. Deeply conserved synteny resolves early events in vertebrate evolution. Nat Ecol Evol 4, 820–830 (2020). 10.1038/s41559-020-1156-z

20 Technau, U. et al. Sea anemone genomes reveal ancestral metazoan chromosomal macrosynteny. Research Square 10.21203/rs.3.rs-796229/v1 (2021). 10.21203/rs.3.rs-796229/v1

21 Muffato, M., Louis, A., Nguyen, N. T. T., Lucas, J., Berthelot, C. & Roest Crollius, H. Reconstruction of hundreds of reference ancestral genomes across the eukaryotic kingdom. Nat Ecol Evol 7, 355–366 (2023). 10.1038/s41559-022-01956-z

22 Sacerdot, C., Louis, A., Bon, C., Berthelot, C. & Roest Crollius, H. Chromosome evolution at the origin of the ancestral vertebrate genome. Genome Biol 19, 166 (2018). 10.1186/s13059-018-1559-1

23 Wang, S. et al. Scallop genome provides insights into evolution of bilaterian karyotype and development. Nat Ecol Evol 1, 120 (2017). 10.1038/s41559-017-0120

24 Marletaz, F. et al. Analysis of the P. lividus sea urchin genome highlights contrasting trends of genomic and regulatory evolution in deuterostomes. Cell Genom 3, 100295 (2023). 10.1016/j.xgen.2023.100295

25 Arshinoff, B. I. et al. Echinobase: leveraging an extant model organism database to build a knowledgebase supporting research on the genomics and biology of echinoderms. Nucleic Acids Res 50, D970–D979 (2022). 10.1093/nar/gkab1005

26 Cameron, R. A., Kudtarkar, P., Gordon, S. M., Worley, K. C. & Gibbs, R. A. Do echinoderm genomes measure up? Mar Genomics 22, 1–9 (2015). 10.1016/j.margen.2015.02.004

27 Yoshida, K. et al. Chromosome fusions repatterned recombination rate and facilitated reproductive isolation during Pristionchus nematode speciation. Nat Ecol Evol 7, 424–439 (2023). 10.1038/s41559-022-01980-z

28 Auclair, W. The Chromosomes of Sea Urchins, Especially Arbacia punctulata; A Method for Studying Unsectioned Eggs at First Cleavage. Biological Bulletin 128, 169–176 (1965).

29 Colombera, D., Vitturi, R. & Zanirato, L. Chromosome-Number of Cidaris-Cidaris-(Cidaridae-Echinoidea). Acta Zool-Stockholm 58, 185–186 (1977). DOI 10.1111/j.1463-6395.1977.tb00254.x

30 Thompson, J. R., Petsios, E., Davidson, E. H., Erkenbrack, E. M., Gao, F. & Bottjer, D. J. Reorganization of sea urchin gene regulatory networks at least 268 million years ago as revealed by oldest fossil cidaroid echinoid. Sci Rep-Uk 5 (2015). 10.1038/srep15541

31 Kroh, A. & Smith, A. B. The phylogeny and classification of post-Palaeozoic echinoids. J Syst Palaeontol 8, 147–212 (2010). 10.1080/14772011003603556

32 Ran, Z. et al. Chromosome-level genome assembly of the razor clam Sinonovacula constricta (Lamarck, 1818). Mol Ecol Resour 19, 1647–1658 (2019). 10.1111/1755-0998.13086

33 Sun, Y. et al. Genomic Signatures Supporting the Symbiosis and Formation of Chitinous Tube in the Deep-Sea Tubeworm Paraescarpia echinospica. Mol Biol Evol 38, 4116–4134 (2021). 10.1093/molbev/msab203

34 Yan, X. et al. Clam Genome Sequence Clarifies the Molecular Basis of Its Benthic Adaptation and Extraordinary Shell Color Diversity. iScience 19, 1225–1237 (2019). 10.1016/j.isci.2019.08.049

35 Zakas, C., Harry, N. D., Scholl, E. H. & Rockman, M. V. The Genome of the Poecilogonous Annelid Streblospio benedicti. Genome Biol Evol 14 (2022). 10.1093/gbe/evac008

36 Martin-Zamora, F. M. et al. Annelid functional genomics reveal the origins of bilaterian life cycles. Nature (2023). 10.1038/s41586-022-05636-7

37 Cannon, J. T., Vellutini, B. C., Smith, J., Onquist, F. R., Jondelius, U. & Hejnol, A. Xenacoelomorpha is the sister group to Nephrozoa. Nature 530, 89-+ (2016). 10.1038/nature16520

38 Rouse, G. W., Wilson, N. G., Carvajal, J. I. & Vrijenhoek, R. C. New deep-sea species of Xenoturbella and the position of Xenacoelomorpha. Nature 530, 94-+ (2016). 10.1038/nature16545

39 Schiffer, P. H. et al. The slowly evolving genome of the xenacoelomorph worm Xenoturbella bocki. bioRxiv (2023). 10.1101/2022.06.24.497508

40 Duboule, D. The (unusual) heuristic value of Hox gene clusters; a matter of time? Dev Biol 484, 75–87 (2022). 10.1016/j.ydbio.2022.02.007

41 Stock, D. W., Ellies, D. L., Zhao, Z. Y., Ekker, M., Ruddle, F. H. & Weiss, K. M. The evolution of the vertebrate Dlx gene family. P Natl Acad Sci USA 93, 10858–10863 (1996). DOI 10.1073/pnas.93.20.10858

42 Bourque, G. et al. Ten things you should know about transposable elements. Genome Biology 19 (2018). 10.1186/s13059-018-1577-z

43 Klein, S. J. & O’Neill, R. J. Transposable elements: genome innovation, chromosome diversity, and centromere conflict. Chromosome Res 26, 5–23 (2018). 10.1007/s10577-017-9569-5

44 Amemiya, C. T. et al. The amphioxus Hox cluster: characterization, comparative genomics, and evolution. J Exp Zool B Mol Dev Evol 310, 465–477 (2008). 10.1002/jez.b.21213

45 Fried, C., Prohaska, S. J. & Stadler, P. F. Exclusion of repetitive DNA elements from gnathostome Hox clusters. J Exp Zool B Mol Dev Evol 302, 165–173 (2004). 10.1002/jez.b.20007

46 Holland, L. Z. et al. The amphioxus genome illuminates vertebrate origins and cephalochordate biology. Genome Res 18, 1100–1111 (2008). 10.1101/gr.073676.107

47 Pascual-Anaya, J., Adachi, N., Alvarez, S., Kuratani, S., D’Aniello, S. & Garcia-Fernandez, J. Broken colinearity of the amphioxus Hox cluster. Evodevo 3, 28 (2012). 10.1186/2041-9139-3-28

48 Ferrier, D. E., Minguillon, C., Holland, P. W. & Garcia-Fernandez, J. The amphioxus Hox cluster: deuterostome posterior flexibility and Hox14. Evol Dev 2, 284–293 (2000). 10.1046/j.1525-142x.2000.00070.x

49 Zhang, X. J. et al. The sea cucumber genome provides insights into morphological evolution and visceral regeneration. Plos Biol 15 (2017). 10.1371/journal.pbio.2003790

50 Perez-Posada, A. et al. Insights into deuterostome evolution from the biphasic transcriptional programmes of hemichordates. bioRxiv (2022). 10.1101/2022.06.10.495707

51 Stefanik, D. J., Wolenski, F. S., Friedman, L. E., Gilmore, T. D. & Finnerty, J. R. Isolation of DNA, RNA and protein from the starlet sea anemone Nematostella vectensis. Nat Protoc 8, 892–899 (2013). 10.1038/nprot.2012.151

52 Nowoshilow, S. et al. The axolotl genome and the evolution of key tissue formation regulators. Nature 554, 50–55 (2018). 10.1038/nature25458

53 Roach, M. J., Schmidt, S. A. & Borneman, A. R. Purge Haplotigs: allelic contig reassignment for third-gen diploid genome assemblies. BMC Bioinformatics 19, 460 (2018). 10.1186/s12859-018-2485-7

54 Walker, B. J. et al. Pilon: an integrated tool for comprehensive microbial variant detection and genome assembly improvement. PLoS One 9, e112963 (2014). 10.1371/journal.pone.0112963

55 Koren, S., Walenz, B. P., Berlin, K., Miller, J. R., Bergman, N. H. & Phillippy, A. M. Canu: scalable and accurate long-read assembly via adaptive k-mer weighting and repeat separation. Genome Res 27, 722–736 (2017). 10.1101/gr.215087.116

56 Chin, C. S. et al. Nonhybrid, finished microbial genome assemblies from long-read SMRT sequencing data. Nat Methods 10, 563–569 (2013). 10.1038/nmeth.2474

57 Manni, M., Berkeley, M. R., Seppey, M. & Zdobnov, E. M. BUSCO: Assessing Genomic Data Quality and Beyond. Curr Protoc 1, e323 (2021). 10.1002/cpz1.323

58 Holt, C. & Yandell, M. MAKER2: an annotation pipeline and genome-database management tool for second-generation genome projects. BMC Bioinformatics 12, 491 (2011). 10.1186/1471-2105-12-491

59 Keilwagen, J., Hartung, F., Paulini, M., Twardziok, S. O. & Grau, J. Combining RNA-seq data and homology-based gene prediction for plants, animals and fungi. BMC Bioinformatics 19, 189 (2018). 10.1186/s12859-018-2203-5

60 Dobin, A. et al. STAR: ultrafast universal RNA-seq aligner. Bioinformatics 29, 15–21 (2013). 10.1093/bioinformatics/bts635

61 Pertea, M., Pertea, G. M., Antonescu, C. M., Chang, T. C., Mendell, J. T. & Salzberg, S. L. StringTie enables improved reconstruction of a transcriptome from RNA-seq reads. Nat Biotechnol 33, 290–295 (2015). 10.1038/nbt.3122

62 Song, L., Sabunciyan, S. & Florea, L. CLASS2: accurate and efficient splice variant annotation from RNA-seq reads. Nucleic Acids Res 44, e98 (2016). 10.1093/nar/gkw158

63 Grabherr, M. G. et al. Full-length transcriptome assembly from RNA-Seq data without a reference genome. Nat Biotechnol 29, 644–652 (2011). 10.1038/nbt.1883

64 Li, H. Minimap2: pairwise alignment for nucleotide sequences. Bioinformatics 34, 3094–3100 (2018). 10.1093/bioinformatics/bty191

65 Salmela, L. & Rivals, E. LoRDEC: accurate and efficient long read error correction. Bioinformatics 30, 3506–3514 (2014). 10.1093/bioinformatics/btu538

66 Haas, B. J. et al. Automated eukaryotic gene structure annotation using EVidenceModeler and the Program to Assemble Spliced Alignments. Genome Biol 9, R7 (2008). 10.1186/gb-2008-9-1-r7

67 Haas, B. J. et al. Improving the Arabidopsis genome annotation using maximal transcript alignment assemblies. Nucleic Acids Res 31, 5654–5666 (2003). 10.1093/nar/gkg770

68 Conesa, A. & Gotz, S. Blast2GO: A comprehensive suite for functional analysis in plant genomics. Int J Plant Genomics 2008, 619832 (2008). 10.1155/2008/619832

69 Cantalapiedra, C. P., Hernandez-Plaza, A., Letunic, I., Bork, P. & Huerta-Cepas, J. eggNOG-mapper v2: Functional Annotation, Orthology Assignments, and Domain Prediction at the Metagenomic Scale. Mol Biol Evol 38, 5825–5829 (2021). 10.1093/molbev/msab293

70 Huerta-Cepas, J. et al. eggNOG 5.0: a hierarchical, functionally and phylogenetically annotated orthology resource based on 5090 organisms and 2502 viruses. Nucleic Acids Res 47, D309–D314 (2019). 10.1093/nar/gky1085

71 Stanke, M., Keller, O., Gunduz, I., Hayes, A., Waack, S. & Morgenstern, B. AUGUSTUS: ab initio prediction of alternative transcripts. Nucleic Acids Res 34, W435–439 (2006). 10.1093/nar/gkl200

72 Slater, G. S. & Birney, E. Automated generation of heuristics for biological sequence comparison. BMC Bioinformatics 6, 31 (2005). 10.1186/1471-2105-6-31

73 Bruna, T., Hoff, K. J., Lomsadze, A., Stanke, M. & Borodovsky, M. BRAKER2: automatic eukaryotic genome annotation with GeneMark-EP+ and AUGUSTUS supported by a protein database. NAR Genom Bioinform 3, lqaa108 (2021). 10.1093/nargab/lqaa108

74 Buchfink, B., Xie, C. & Huson, D. H. Fast and sensitive protein alignment using DIAMOND. Nat Methods 12, 59–60 (2015). 10.1038/nmeth.3176

75 Gotoh, O. A space-efficient and accurate method for mapping and aligning cDNA sequences onto genomic sequence. Nucleic Acids Res 36, 2630–2638 (2008). 10.1093/nar/gkn105

76 Hoff, K. J., Lange, S., Lomsadze, A., Borodovsky, M. & Stanke, M. BRAKER1: Unsupervised RNA-Seq-Based Genome Annotation with GeneMark-ET and AUGUSTUS. Bioinformatics 32, 767–769 (2016). 10.1093/bioinformatics/btv661

77 Iwata, H. & Gotoh, O. Benchmarking spliced alignment programs including Spaln2, an extended version of Spaln that incorporates additional species-specific features. Nucleic Acids Res 40, e161 (2012). 10.1093/nar/gks708

78 Lomsadze, A., Ter-Hovhannisyan, V., Chernoff, Y. O. & Borodovsky, M. Gene identification in novel eukaryotic genomes by self-training algorithm. Nucleic Acids Res 33, 6494–6506 (2005). 10.1093/nar/gki937

79 Stanke, M., Diekhans, M., Baertsch, R. & Haussler, D. Using native and syntenically mapped cDNA alignments to improve de novo gene finding. Bioinformatics 24, 637–644 (2008). 10.1093/bioinformatics/btn013

80 Bruna, T., Lomsadze, A. & Borodovsky, M. GeneMark-EP+: eukaryotic gene prediction with self-training in the space of genes and proteins. NAR Genom Bioinform 2, lqaa026 (2020). 10.1093/nargab/lqaa026

81 Tang, H., Bowers, J. E., Wang, X., Ming, R., Alam, M. & Paterson, A. H. Synteny and collinearity in plant genomes. Science 320, 486–488 (2008). 10.1126/science.1153917

82 Wang, Y. et al. MCScanX: a toolkit for detection and evolutionary analysis of gene synteny and collinearity. Nucleic Acids Res 40, e49 (2012). 10.1093/nar/gkr1293

83 Suchard, M. A., Lemey, P., Baele, G., Ayres, D. L., Drummond, A. J. & Rambaut, A. Bayesian phylogenetic and phylodynamic data integration using BEAST 1.10. Virus Evol 4, vey016 (2018). 10.1093/ve/vey016

84 Supek, F., Bosnjak, M., Skunca, N. & Smuc, T. REVIGO summarizes and visualizes long lists of gene ontology terms. PLoS One 6, e21800 (2011). 10.1371/journal.pone.0021800

85 Flynn, J. M. et al. RepeatModeler2 for automated genomic discovery of transposable element families. Proc Natl Acad Sci U S A 117, 9451–9457 (2020). 10.1073/pnas.1921046117

86 Quinlan, A. R. & Hall, I. M. BEDTools: a flexible suite of utilities for comparing genomic features. Bioinformatics 26, 841–842 (2010). 10.1093/bioinformatics/btq033

87 Ramirez, F., Dundar, F., Diehl, S., Gruning, B. A. & Manke, T. deepTools: a flexible platform for exploring deep-sequencing data. Nucleic Acids Res 42, W187–191 (2014). 10.1093/nar/gku365

88 Buels, R. et al. JBrowse: a dynamic web platform for genome visualization and analysis. Genome Biology 17 (2016). 10.1186/s13059-016-0924-1

