## Supplementary Figures for "Chromosome-level genome assemblies of two hemichordates provide new insights into deuterostome origin and chromosome evolution"

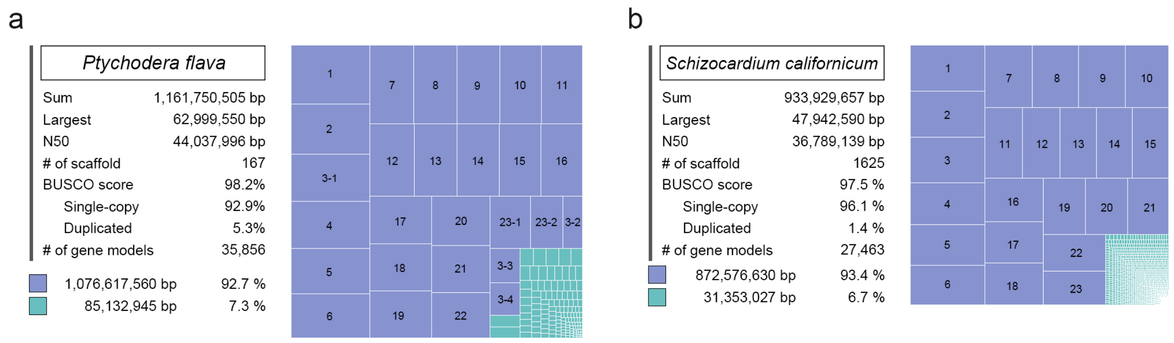


**Supplementary Fig. 1 | Chromosome-level genome assemblies of the two hemichordates.** Statistical data (left) and treemap (right) of *P. flava* (**a**) and *S. californicum* (**b**) genome assemblies based on PacBio long reads; 27 and 23 larger scaffolds of *P. flava* and *S. californicum* were taken into chromosomal sequences and denoted by blue boxes. The green boxes represent the remaining scaffolds.

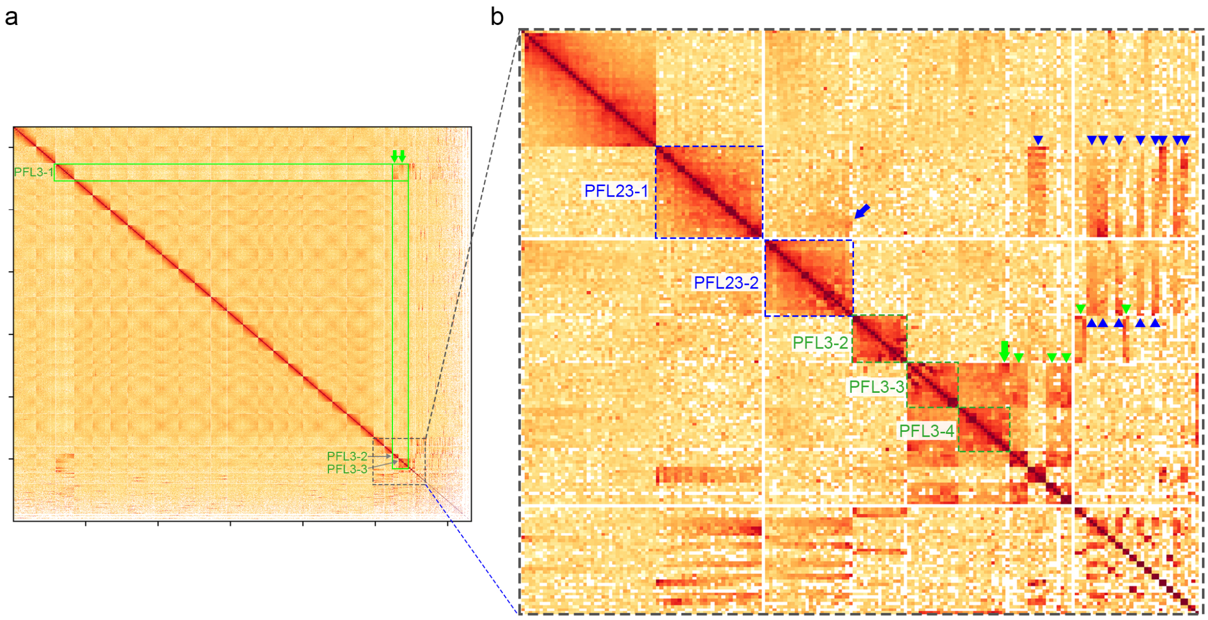


**Supplementary Fig. 2 | Further analysis of the HiC dataset on the *P. flava* genome assembly using the HiC-pro pipeline.**

**a**, A Hi-C contact map of *P. flava* genome assembly based on the HiC-pro pipeline[1]. Note that the 3' end of PFL3-1 interacts with the 5' end of PFL3-2, and the 5' end of PFL3-1 interacts with the 3' end of PFL3-3 (green arrows). The boxed area is magnified to show the chromosomal interactions around the PFL chromosome 23. **b,** The 3' end (right side) of PFL23-1 interacts with the 3' end of PFL23-2 (blue arrow), suggesting that the two scaffolds are closely linked at their 3' ends. These two scaffolds also highly interact with several smaller scaffolds (blue arrowheads). Similarly, the 3' end of PFL3-4 interacts with the 5' end of PFL3-3 (green arrow). Based on the contact information, PFL3-1 to PFL3-4 were assembled in the order of PFL3-4, PFL3-3, PFL3-1 and PFL3-2. PFL3-2 and PFL3-3 also interact with several smaller scaffolds (green arrowheads). *P. flava* chromosome #3 (PFL3) was thus assembled by joining PFL3-1 to PFL3-4; PFL23 was assembled by joining PFL23-1 and PFL23-2.

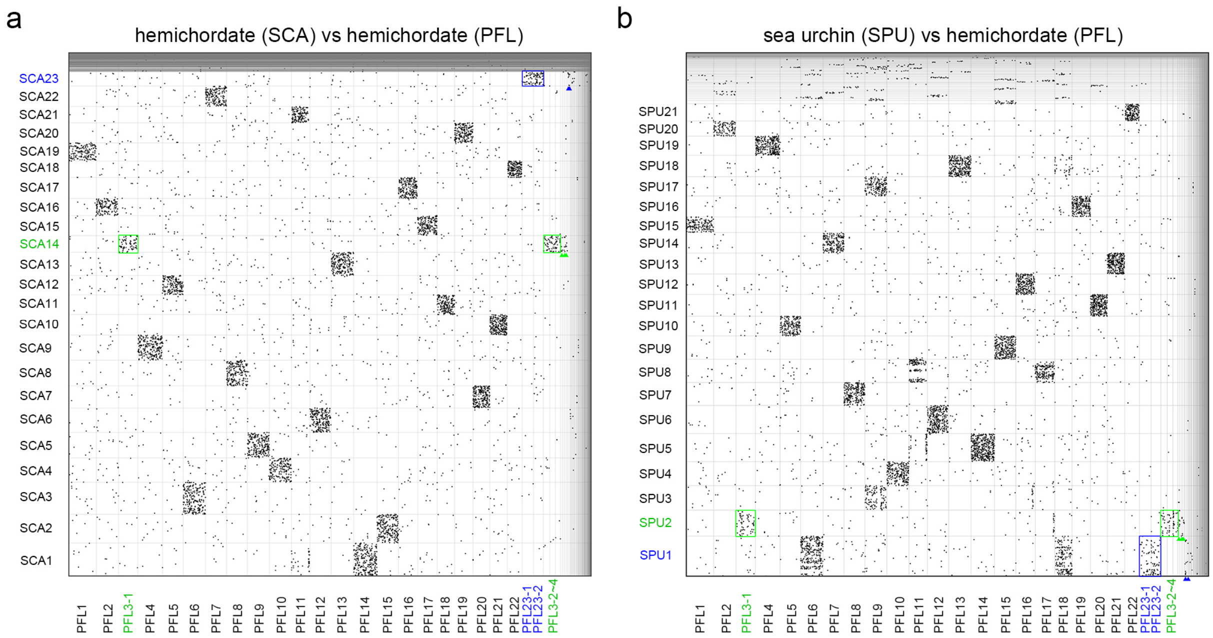


**Supplementary Fig. 3 | Syntenic dot plots between *P. flava* and two deuterostome species.**

Each dot denotes an orthologous gene pair identified between two hemichordates SCA and PFL (a) or between sea urchin SPU and hemichordate PFL (b). Chromosomes/scaffolds are separated by gray lines. *P. flava* PFL3-1, PFL3-2, PFL3-3 and PFL3-4 (green boxes) correspond to *S. californicum* SCA14 (a) and *S. purpuratus* SPU2 (b), further supporting the conclusion that PFL3-1 to PFL3-4 constitute the same chromosome. PFL23-1 and PFL23-2 (blue boxes) correspond to SCA23 (a) and SPU1 (b), supporting the conclusion that PFL23-1 and PFL23-2 are from the same chromosome. Notably, comparison of the two hemichordate genomes did not show apparent microsynteny conservation, suggesting that large-scale intra-chromosomal rearrangements occurred at least in one of the two lineages leading to the two hemichordate species.

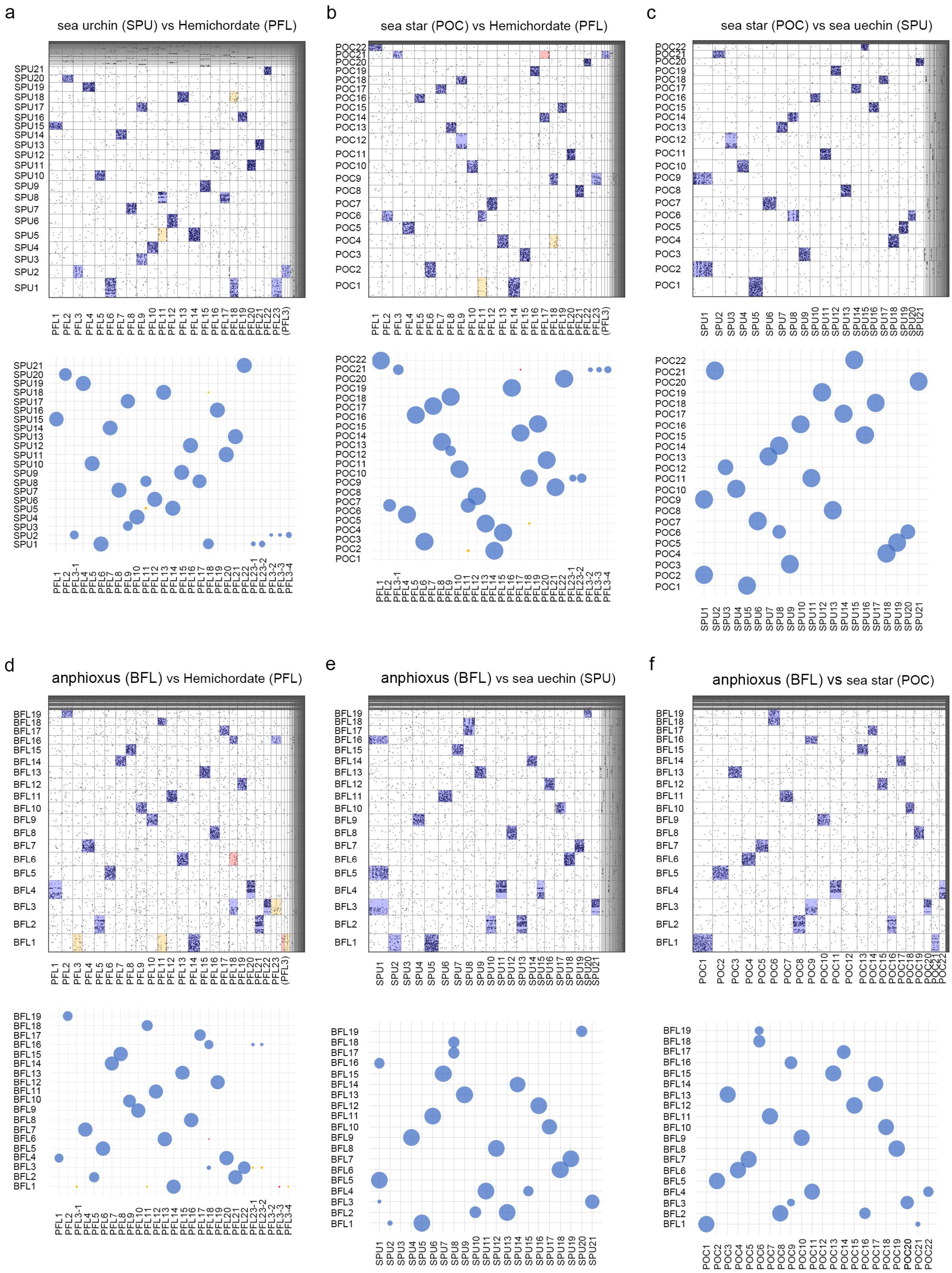


**Supplementary Fig. 4 | Pairwise syntenic dot plots and significant associations between deuterostome species.**

Dot plots (upper panels) showing the chromosomal positions of orthologous gene pairs between two species. Statistically corresponding chromosomes are shaded based on significance level in Fisher's exact test and risk difference. In the scatter plots (lower panels), the circle sizes depict the -log 10 adjusted *p*-value, with a maximum of 300 for each plot. Adjusted *p*-values < 1E-10, between 1E-5~1E-10 and between 1E-2~1E-5 are marked respectively with blue, yellow and red. Adjusted *p*-values > 1E-2 or risk difference < 0 are not shown. For PFL3-1 to PFL3-4 and PFL23-1 to PFL23-2, significance of difference was calculated separately.

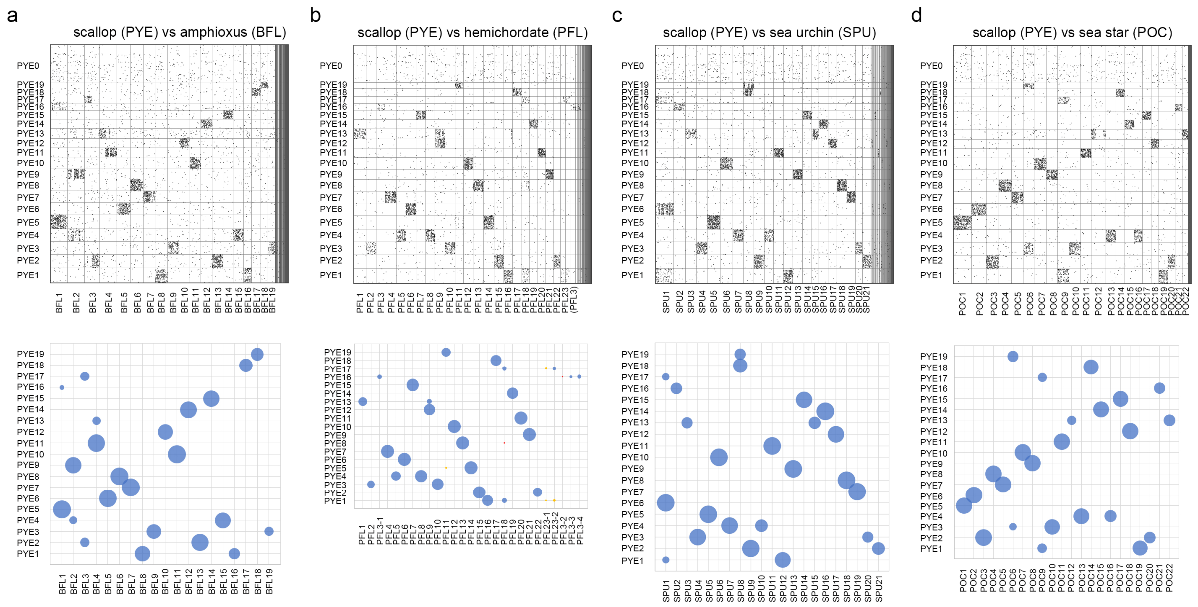


**Supplementary Fig. 5 | Pairwise syntenic dot plots and significant associations between genomes of deuterostome species and the scallop (PYE).**

Dot plots showing the chromosomal positions of orthologous gene pairs identified between scallop PYE and amphioxus BFL (a), hemichordate PFL (b), sea urchin SPU (c) or sea star POC (d). All symbols are the same as those described in Supplementary Fig. 4. PYE0 is an unplaced scaffold[2].

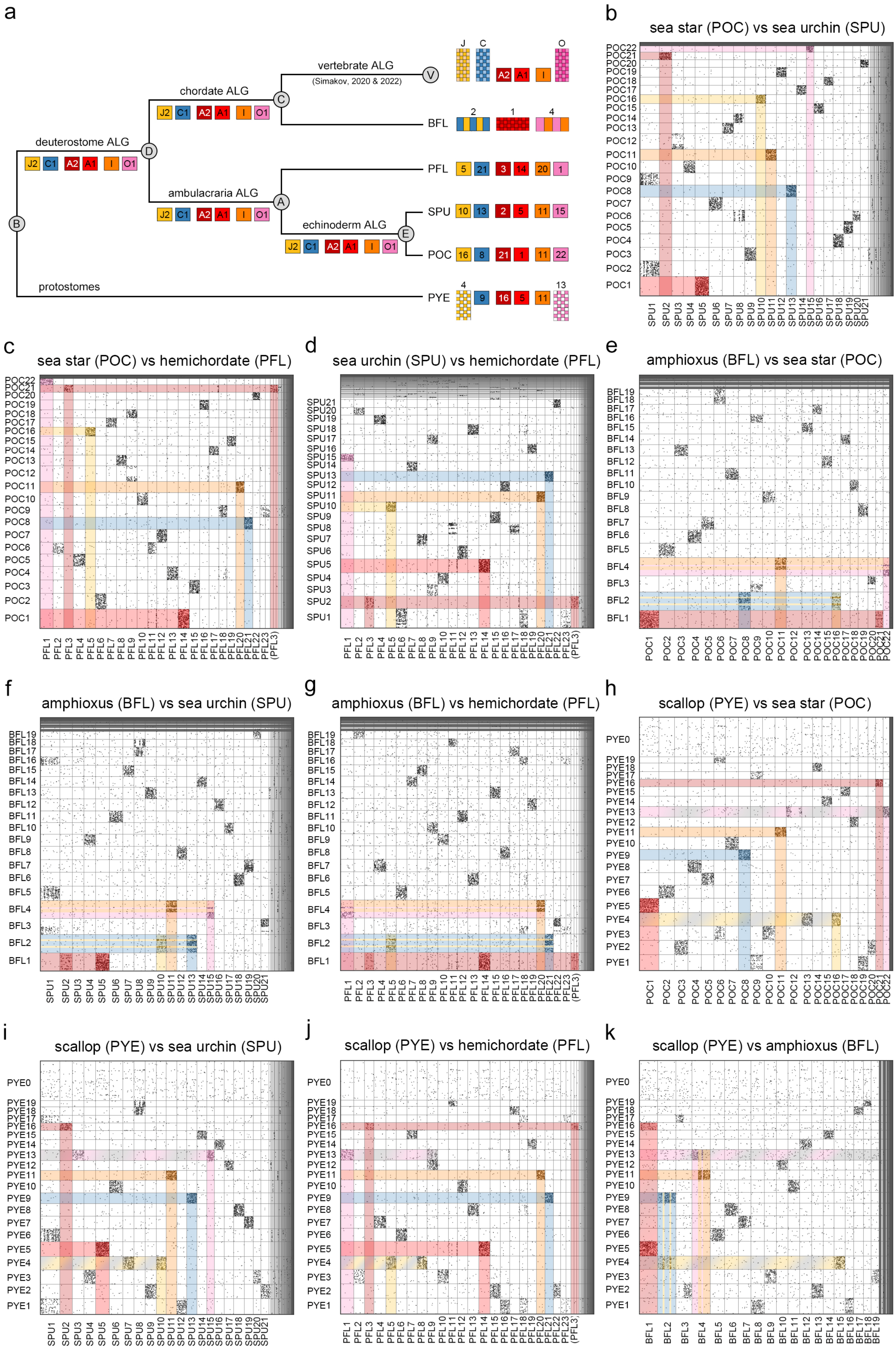
a
vertebrate ALG
(Simakov, 2020 & 2022)
chordate ALG
deuterostome ALG
ambulacraria ALG
echinoderm ALG
protostomes
V
C
D
A
E
B
J2 C1 A2 A1 I O1
J C O
BFL
PFL
SPU
POC
PYE
b
sea star (POC) vs sea urchin (SPU)
c
sea star (POC) vs hemichordate (PFL)
d
sea urchin (SPU) vs hemichordate (PFL)
e
amphioxus (BFL) vs sea star (POC)
f
amphioxus (BFL) vs sea urchin (SPU)
g
amphioxus (BFL) vs hemichordate (PFL)
h
scallop (PYE) vs sea star (POC)
i
scallop (PYE) vs sea urchin (SPU)
j
scallop (PYE) vs hemichordate (PFL)
k
scallop (PYE) vs amphioxus (BFL)

**Supplementary Fig. 6 | Chromosome evolution of deuterostome ALGs J2, C1, A2, A1, I and O1.**

**a**. Reconstruction of deuterostome ALGs J2, C1, A2, A1, I and O1 based on pairwise comparisons among amphioxus BFL, hemichordate PFL, sea urchin SPU, sea star POC and scallop PYE. First, the comparison of POC with SPU showed that POC16, POC8, POC21, POC1, POC11 and POC22 have one-to-one correspondence with SPU10, SPU13, SPU2, SPU5, SPU11 and SPU15, respectively (**b**), suggesting that these six chromosomes were already present in their LCA (echinoderm ALGs J2, C1, A2, A1, I and O1). These six chromosomes also have one-to-one correspondence with hemichordate PFL5, PFL21, PFL3, PFL14, PFL20 and PFL1 (**c** and **d**), indicating that the existence of these six chromosomes could be traced further back to the ambulacrarian LCA (ambulacraria ALGs J2, C1, A2, A1, I and O1). Comparisons with the amphioxus BFL genome showed that both POC16/SPU10/PFL5 and POC8/ SPU13/PFL21 correspond to a single amphioxus chromosome BFL2 (**e**-**g**). Similarly, POC21/SPU2/PFL3 and POC1/ SPU5/PFL14 correspond to amphioxus BFL1; POC11/SPU11/PFL20 and POC22/ SPU15/PFL1 correspond to amphioxus BFL4. To infer the deuterostome ancestral condition, scallop PYE was used as an outgroup. This analysis showed that the six ambulacraria chromosomes correspond to six distinct PYE chromosomes (PYE4, PYE9, PYE16, PYE5, PYE11 and PYE13, see **h**-**j**), supporting the conclusion that these six chromosomes are ancient and were present in the deuterostome LCA (deuterostome ALGs J2, C1, A2, A1, I and O1). Accordingly, the three amphioxus chromosomes (BFL 2, BFL1 and BFL4) correspond to the aforementioned six PYE chromosomes (**k**). Therefore, the amphioxus BFL2, BFL1 and BFL4 were formed from respective fusion events between deuterostome ALGs J2 and C1, ALGs A2 and A1, and ALGs I and O1. These three fusion events are likely amphioxus-specific because the six deuterostome ALGs correspond to six vertebrate ALGs[3,4], which support the notion that these six chromosomes remained intact in the LCA of chordates (chordate ALGs J2, C1, A2, A1, I and O1).

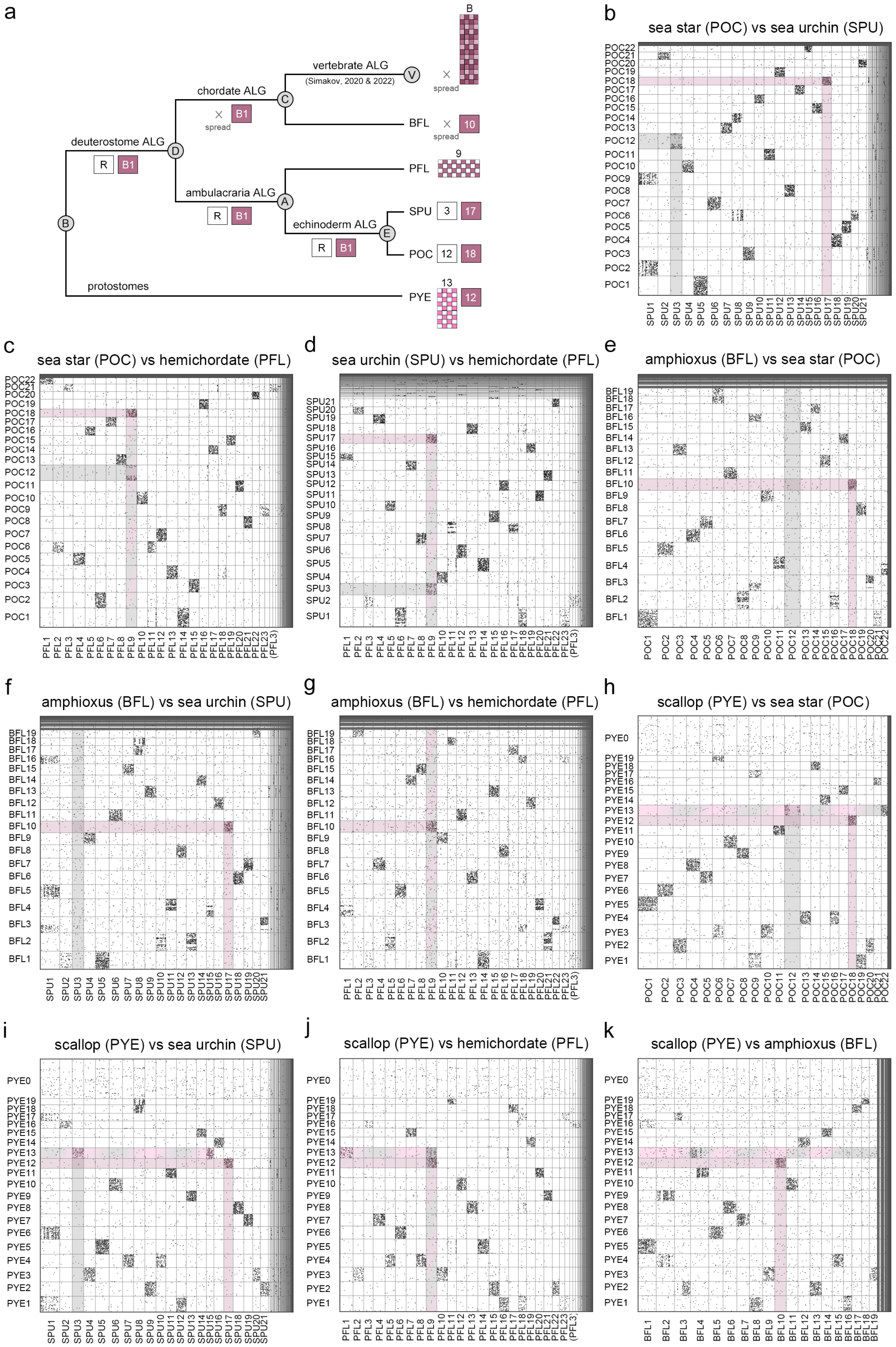
a
B
vertebrate ALG
(Simakov, 2020 & 2022)
V
spread
chordate ALG
C
B1
spread
BFL
10
spread
deuterostome ALG
D
R
B1
9
PFL
ambulacraria ALG
A
R
B1
SPU
3
17
echinoderm ALG
E
R
B1
POC
12
18
B
13
protostomes
PYE
12
b
sea star (POC) vs sea urchin (SPU)
c
sea star (POC) vs hemichordate (PFL)
d
sea urchin (SPU) vs hemichordate (PFL)
e
amphioxus (BFL) vs sea star (POC)
f
amphioxus (BFL) vs sea urchin (SPU)
g
amphioxus (BFL) vs hemichordate (PFL)
h
scallop (PYE) vs sea star (POC)
i
scallop (PYE) vs sea urchin (SPU)
j
scallop (PYE) vs hemichordate (PFL)
k
scallop (PYE) vs amphioxus (BFL)

**Supplementary Fig. 7 | Chromosome evolution of deuterostome ALGs R and B1.**
**a**. Reconstruction of deuterostome ALGs R and B1 based on pairwise comparisons. Using the same logic as described for Supplementary Fig. 6, sea star POC12 and POC18 appear to correspond to sea urchin SPU3 and SPU17, respectively (**b**), supporting the conclusion that their LCA possessed these two chromosomes (echinoderm ALGs R and B1). Comparison between hemichordate PFL and echinoderm species revealed that POC12/SPU3 and POC18/SPU17 correspond to a single hemichordate chromosome PFL9 (**c** and **d**). This observation suggests a fusion event occurred in the ambulacraria ancestor leading to PFL9 or a split event leading to POC12/SPU3 and POC18/SPU17. Using amphioxus BFL as an outgroup, the analysis showed that POC18/SPU17 corresponds to BFL10 (**e** and **f**), while amphioxus orthologs of POC12/SPU3 genes spread in the genome and no single BFL chromosome could be assigned to POC12/SPU3. Another outgroup scallop PYE was then used, revealing that POC12/SPU3 and POC18/SPU17 respectively correspond to PYE13 and PYE12 (**h** and **i**). Based on these comparisons, three major inferences can be made: (1) both deuterostome and ambulacraria ancestors possessed the two distinct chromosomes (deuterostome/ambulacraria ALGs R and B1); (2) at least in the LCA of hemichordates PFL and SCA, ALGs R and B1 were fused, leading to PFL9/SCA5; (3) in amphioxus, orthologous genes of deuterostome ALG R were dispersed to other chromosomes. Notably, in addition to POC12/SPU3, PYE13 also corresponds to POC22/SPU15, explaining the comparability between PYE13 and the hemichordate PFL1 and amphioxus BFL4 (**j** and **k**) and suggesting a fusion event led to PYE13. Consistent with this idea, the hemichordate PFL9 (fused from ALGs R and B1) corresponds to BFL10 (ALG B1) (**g**). It has been proposed that all chromosomes of vertebrates correspond to amphioxus chromosomes (Simakov, 2020 and 2022), suggesting that one ancestral chromosome (ALG R) spread to other chromosomes in the LCA of chordates. The scallop chromosome name was labeled and sorted according to chromosome size. Here, PYE12 is chromosome number 13 and PYE13 is chromosome number 12 in the previous study[2].

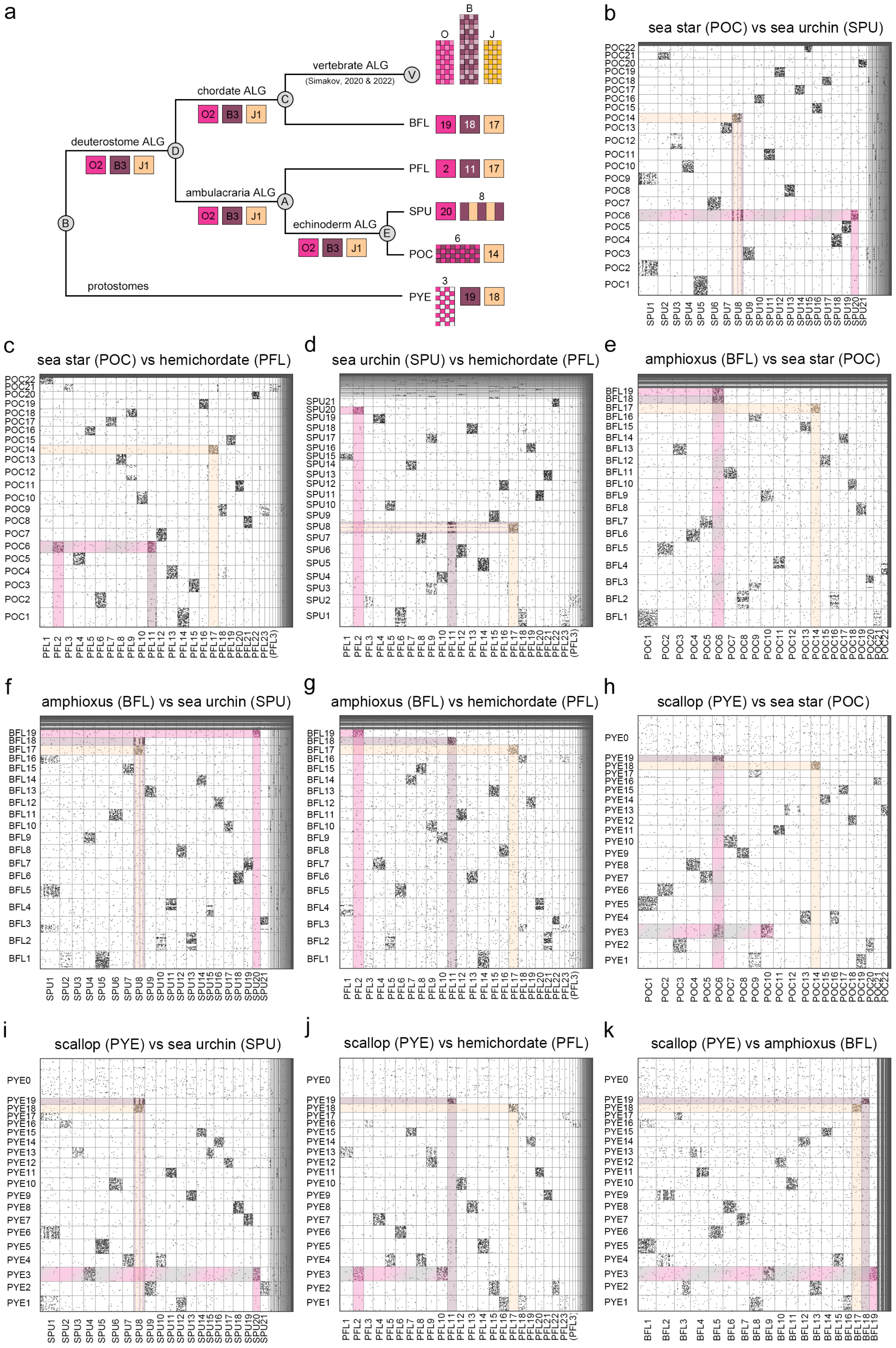
a
vertebrate ALG
(Simakov, 2020 & 2022)
chordate ALG
deuterostome ALG
ambulacraria ALG
echinoderm ALG
protostomes
O2
B3
J1
V
C
D
A
E
B
O
J
BFL
PFL
SPU
POC
PYE
19
18
17
2
11
20
8
6
14
3
b
sea star (POC) vs sea urchin (SPU)
c
sea star (POC) vs hemichordate (PFL)
d
sea urchin (SPU) vs hemichordate (PFL)
e
amphioxus (BFL) vs sea star (POC)
f
amphioxus (BFL) vs sea urchin (SPU)
g
amphioxus (BFL) vs hemichordate (PFL)
h
scallop (PYE) vs sea star (POC)
i
scallop (PYE) vs sea urchin (SPU)
j
scallop (PYE) vs hemichordate (PFL)
k
scallop (PYE) vs amphioxus (BFL)

**Supplementary Fig. 8 | Chromosome evolution of deuterostome ALGs O2, B3 and J1.**
**a**. Reconstruction of deuterostome ALGs O2, B3 and J1 based on pairwise comparisons. The sea star POC6 corresponds to sea urchin SPU20 and SPU8 (**b**) and hemichordate PFL2 and PFL11 (**c**); SPU20 and SPU8 also correspond to these two PFL chromosomes (**d**), indicating that these two chromosomes were present at least in the ambulacrarian and echinoderm LCAs, and POC6 resulted from fusion of the two ancestral chromosomes (ALGs O2 and B3). Intriguingly, in addition to POC6, SPU8 also corresponds to POC14, while POC14 corresponds to a single hemichordate chromosome PFL17. Consistently, SPU8 corresponds to PFL11 and PFL17 (**d**), indicating that a single chromosome corresponding to POC14/PFL17 is an ancestral trait (ALG J1), while SPU8 resulted from chromosomal fusion (ALGs B3 and J1). Therefore, it can be inferred that the LCAs of ambulacrarians and echinoderms possessed these three ALGs (O2, B3 and J1), which remained as individual chromosomes in hemichordates but underwent different fusion events in different echinoderm lineages. Fusion of ALGs O2 and B3 led to sea star POC6, while fusion of ALGs B3 and J1 resulted in sea urchin SPU8. Consistent with this hypothesis, three distinct amphioxus chromosomes BFL19, BFL18 and BFL17 correspond to POC6 and POC14 (**e**), SPU20 and SPU8 (**f**) and PFL2, PFL11 and PFL17 (**g**). This correspondence supports the idea that the presence of the three ALGs can be traced back to the LCA of deuterostomes and remained in the chordate LCA. This conclusion is further reinforced by the observation that the scallop genome contains three distinct chromosomes (PYE3, PYE19 and PYE18) corresponding to POC6 and POC14 (**h**), SPU20 and SPU8 (**i**), PFL2, PFL11 and PFL17 (**j**) and BFL19, BFL18 and BFL17 (**k**). Additionally, the three amphioxus chromosomes BFL19, BFL18 and BFL17 have been shown to correspond to three distinct vertebrate chromosomes[3,4], supporting the conclusion that the chordate LCA possessed these three chromosomes.

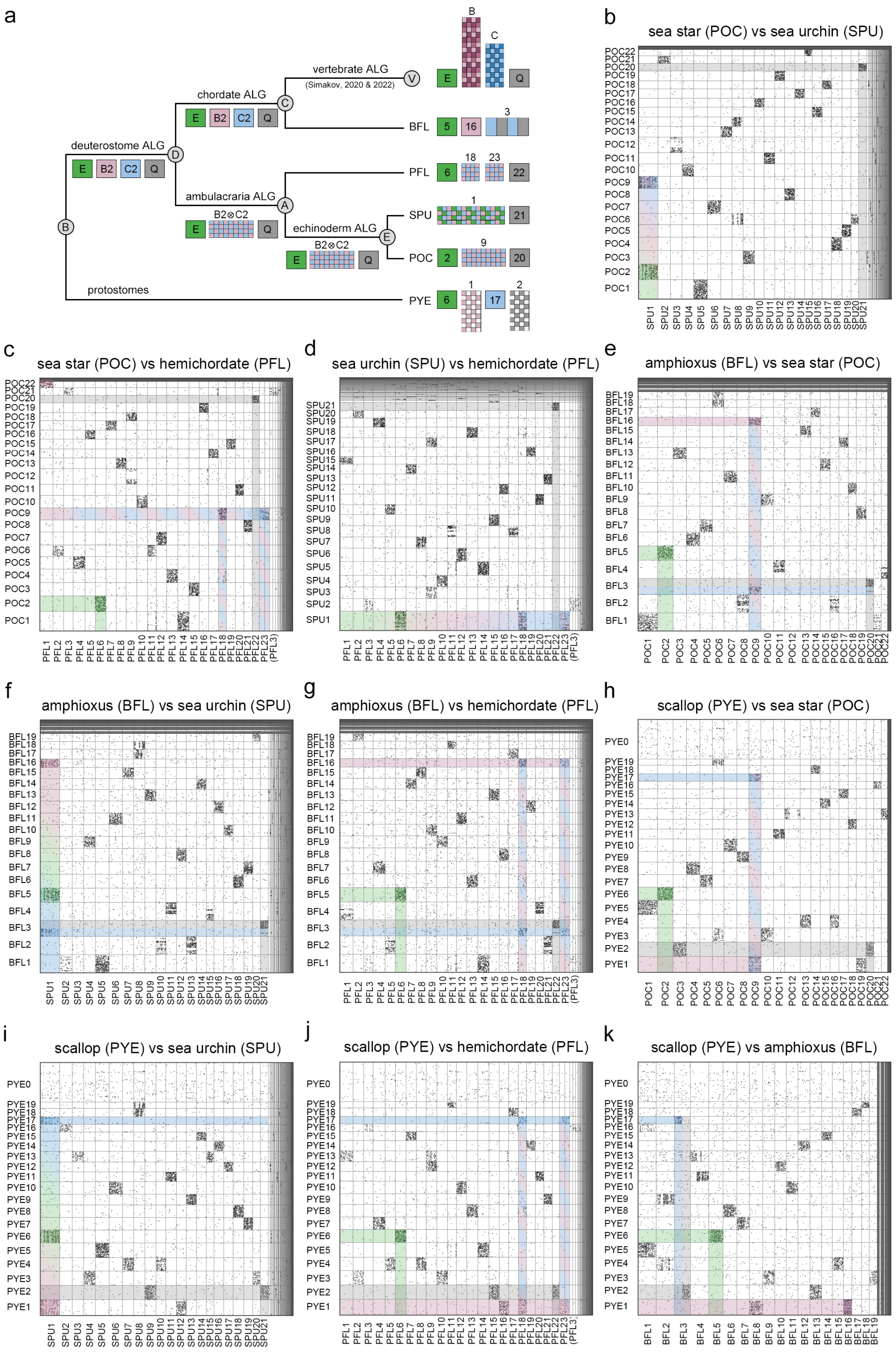
a
vertebrate ALG
(Simakov, 2020 & 2022)
chordate ALG
deuterostome ALG
ambulacraria ALG
echinoderm ALG
protostomes
B2⊗C2
BFL
PFL
SPU
POC
PYE
b
sea star (POC) vs sea urchin (SPU)
c
sea star (POC) vs hemichordate (PFL)
d
sea urchin (SPU) vs hemichordate (PFL)
e
amphioxus (BFL) vs sea star (POC)
f
amphioxus (BFL) vs sea urchin (SPU)
g
amphioxus (BFL) vs hemichordate (PFL)
h
scallop (PYE) vs sea star (POC)
i
scallop (PYE) vs sea urchin (SPU)
j
scallop (PYE) vs hemichordate (PFL)
k
scallop (PYE) vs amphioxus (BFL)

**Supplementary Fig. 9 | Chromosome evolution of deuterostome ALGs E, B2, C2 and Q.** **a**. Reconstruction of deuterostome ALGs E, B2, C2 and Q based on pairwise comparisons. The sea star POC2 and POC9 correspond to sea urchin SPU1 (**b**). POC2 corresponds to a single hemichordate chromosome PFL6, and POC9 corresponds to PFL18 and PFL23 (**c**). These three PFL chromosomes (PFL6, PFL18 and PFL23) also correspond to SPU1 (**d**). This observation suggests that the chromosomes in the sea star (POC2, POC9 and POC20) correspond to those in the LCA of the two echinoderm species, while SPU1 resulted from fusion of the two echinoderm ancestral chromosomes (echinoderm ALGs E and B2⊗C2). To infer the ambulacrarian ancestral condition, the amphioxus BFL genome was compared to the ambulacrarian genomes. POC2 and PFL6 correspond to a single amphioxus chromosome BFL5, supporting the conclusion that echinoderm ALG E has a deeper root in the ambulacrarian LCA and deuterostome LCA (ambulacraria/deuterostome ALG E). On the other hand, POC9 and both PFL18 and PFL23 correspond to two amphioxus chromosomes, BFL16 and BFL3 (**e-g**). Based on this observation, it may be inferred that POC9 could represent the ambulacraria ancestral chromosome (ambulacraria ALG B2⊗C2), and hemichordate PFL18 and PFL23 resulted from a split of ambulacraria ALG B2⊗C2. Notably, in addition to POC9, amphioxus BFL3 also corresponds to POC20 (**e**). POC20 shows one-to-one correspondence with SPU21 and PFL22 (**b-d**), suggesting that an ancestral chromosome was present at least in the LCA of ambulacrarians (ambulacraria ALG Q) and remained intact in the echinoderm lineage (echinoderm ALG Q). To infer the deuterostome ancestral condition and the evolutionary history of BFL3, the scallop PYE genome was compared to those of the deuterostome genomes (**h-k**). The observation that BFL16 corresponds to a single PYE chromosome (PYE1) supports the idea that the deuterostome LCA possessed this chromosome (deuterostome ALG B2). Additionally, BFL3 corresponds to PYE17 and PYE2. PYE2 also corresponds to BFL13 and two one-to-one corresponding chromosomes in ambulacrarian species (POC20/SPU21/PFL22 and POC3/SPU9/PFL15). Therefore, the deuterostome LCA likely possessed ALGs C2 and Q. In the lineage leading to ambulacrarians, deuterostome ALGs B2 and C2 fused and became ambulacraria ALG B2⊗C2. Furthermore, BFL3 also corresponds to two vertebrate chromosomes[3,4], so the chordate LCA likely inherited deuterostome ALGs C2 and Q, and these two chromosomes then fused specifically in amphioxus to become BFL3.

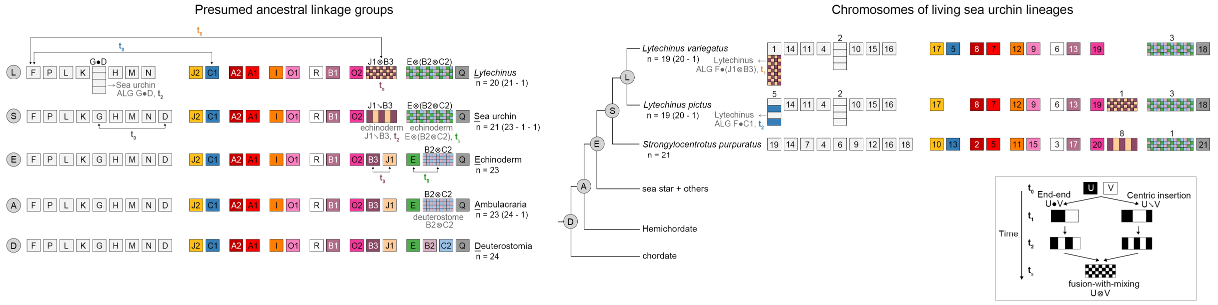


**Supplementary Fig. 10 | Evolutionary history of sea urchin chromosomal architectures.** A stepwise process of sea urchin chromosomal evolution. We divided the process into four time points: $t_0$, $t_1$, $t_2$ and $t_s$ (bottom right panel). At "$t_0$", individual chromosomes have not fused. At "$t_1$", two chromosomes are fused by either end-end translocation or centric insertion. At "$t_2$", intra-chromosomal translocations occur, although long stretches of chromosomal regions are still maintained. At "$t_s$", extensive intra-chromosomal rearrangements have occurred, and the fused chromosome becomes scrambled (fusion-with-mixing). We deduced five major fusion events that occurred during sea urchin chromosomal evolution, as follows. (1) Echinoderm EALGs E and B2$\otimes$C2 fused and mixed to become sea urchin SALG E$\otimes$(B2$\otimes$C2) ($t_0$ to $t_s$ in green). (2) EALGs B3 and J1 fused via centric insertion, followed by translocation to become SALG J1↘B3($t_0$ to $t_2$ in maroon). (3) A *Lytechinus*-specific fusion event resulted from end-end fusion of SALGs G and D without obvious translocation ($t_0$ to $t_1$ in gray). (4) An LVA-specific fusion event involved *Lytechinus* LALGs F and J1$\otimes$B3 without obvious translocation ($t_0$ to $t_1$ in Navajo white). (5) An LPI-specific fusion resulted from end-end fusion of *Lytechinus* LALGs F1 and C1, followed by an intrachromosomal translocation event ($t_0$ to $t_2$ in blue). Box sizes do not reflect the actual sizes of chromosomes.

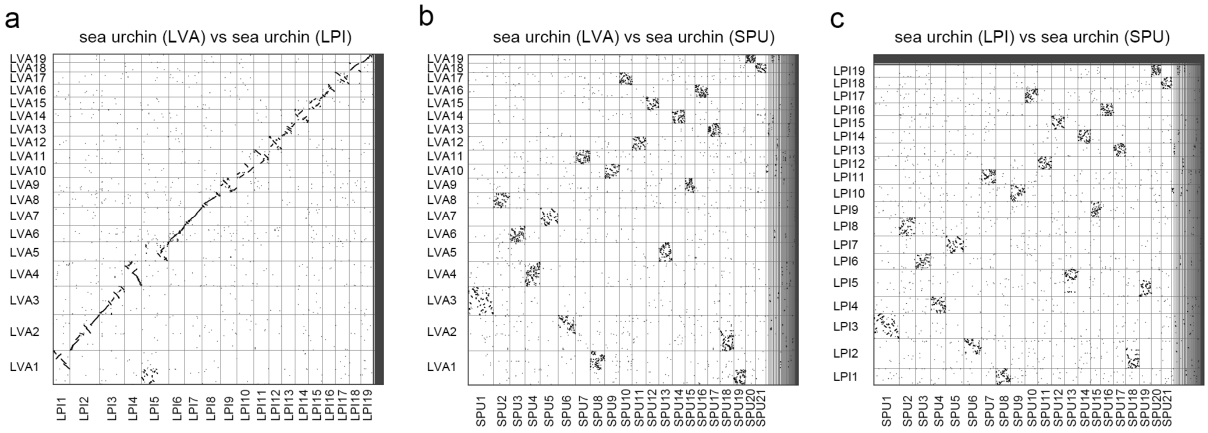


**Supplementary Fig. 11 | Pairwise syntenic dot plots among sea urchin lineages.**
**a.** Syntenic analysis of sea urchin LVA and LPI shows remarkable microsynteny conservation (i.e., linear relationships between chromosome pairs). Sea urchin LVA2 corresponds to SPU6 and SPU18, indicating LVA2 was fused from two ancestral chromosomes (**b**). Similarly, sea urchin LPI2 also corresponds to SPU6 and SPU18 (**c**), suggesting that this fusion event is a common trait in the *Lytechnus* genus. Furthermore, LVA1 corresponds to SPU8 and SPU19 (**b**), and LPI5 corresponds to SPU13 and SPU19 (**c**), indicating additional lineage-specific fusion events in sea urchin LVA and LPI.

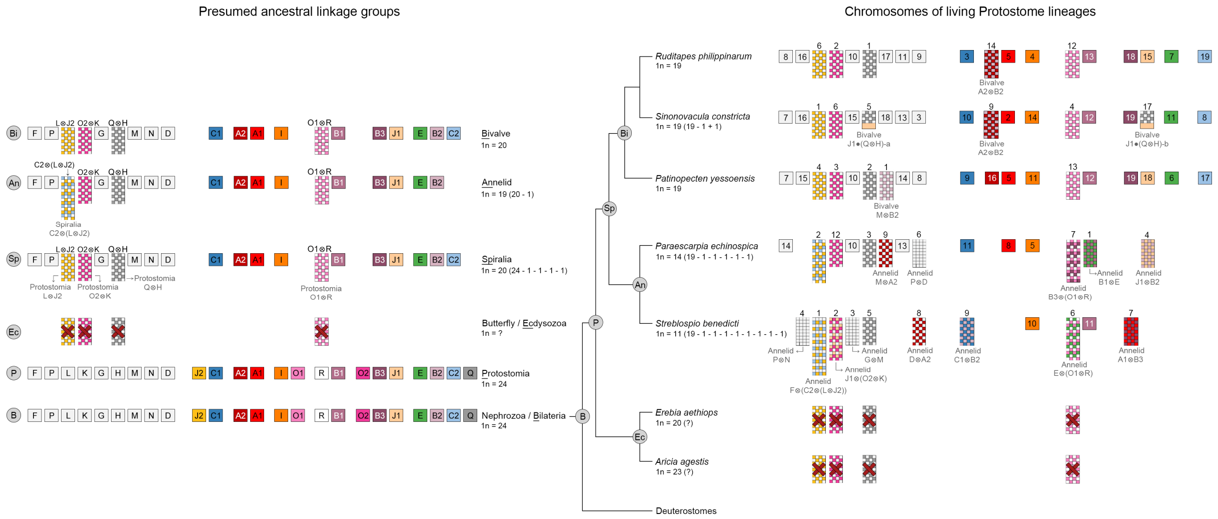


**Supplementary Fig. 12 | Evolutionary history of protostome chromosomal architectures.** The LCA of protostomes likely retained 24 ALGs (PALGs) that show one-to-one correspondence with the 24 bilaterian ALGs. During protostome evolution, different chromosomal rearrangement events occurred in the spiralian and ecdysozoan lineages. All examined spiralian species, including three bivalves and two annelids, share four fusion events (L⊗J2, O2⊗K, Q⊗H and O1⊗R), indicating that their LCA (presumably the LCA of spiralians) already possessed the four fused ALGs, so the overall number of SpALGs is 20. The LCA of the three bivalve species is deduced to have the same complement of ALGs (BiALGs) as the SpALGs, and lineage-specific fusion events are found in the three bivalves (see Supplementary Fig. 13). On the other hand, it can be inferred that the two annelids share an additional fusion event (SpALGs C2 and L⊗J2), which brings the number of the annelid ALGs (AnALGs) to 19. Notably, the four common fusion events in spiralians were not detected in the ecdysozoan species we examined (red crosses over the fused chromosomes) (see Supplementary Fig. 15). Weak syntenic conservation between chromosomes of ecdysozoans and other bilaterians suggests that ecdysozoans underwent more complex chromosomal rearrangements. Box sizes do not reflect the actual sizes of chromosomes.

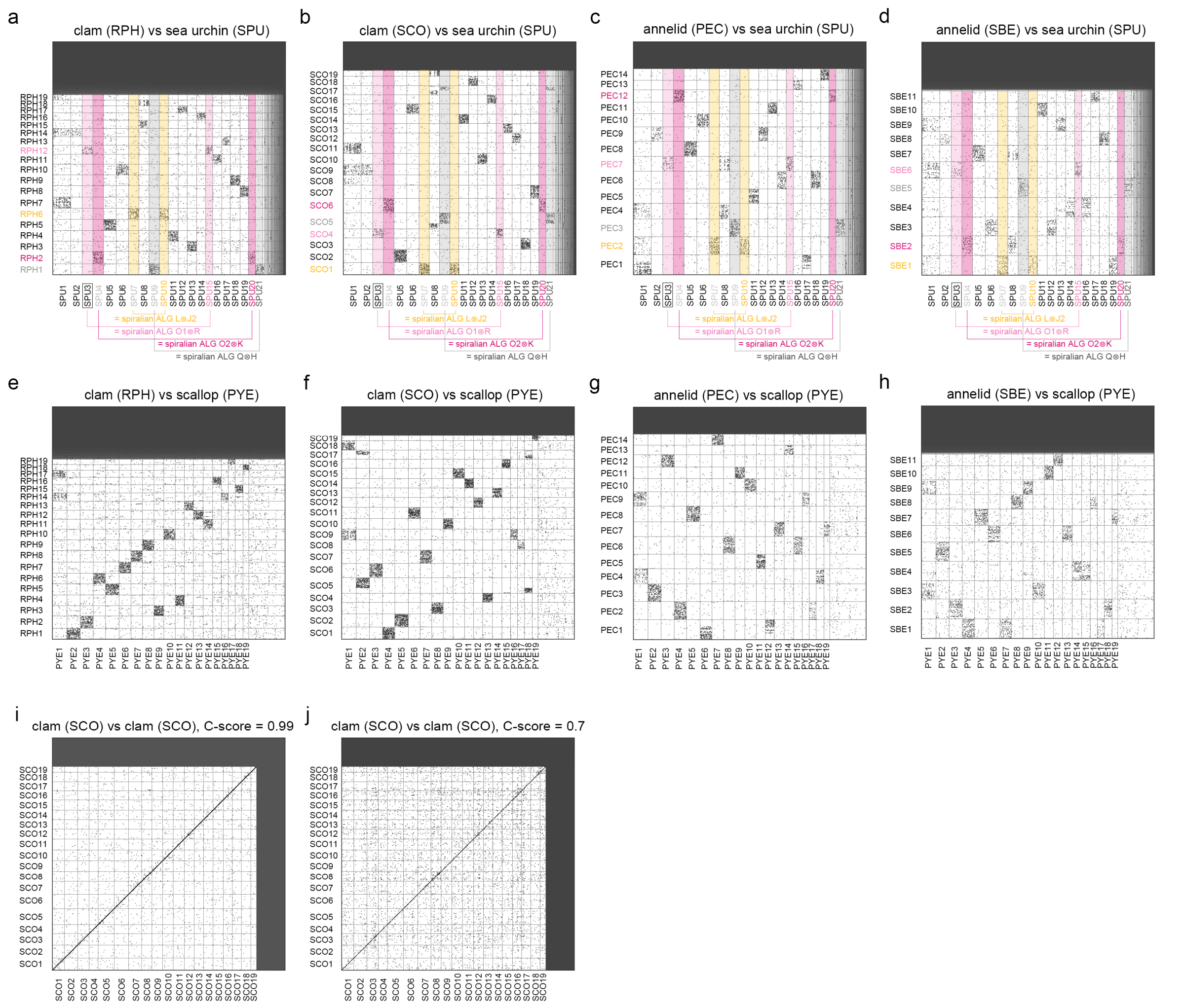


**Supplementary Fig. 13 | Pairwise syntenic dot plots of spiralian chromosomes.** Syntenic analysis showing four common fusion events in the spiralian genomes. For example, PYE3 (see Supplementary Fig. 5), RPH2 (a), SCO6 (b), PEC12 (c) and SBE2 (d) correspond to two sea urchin chromosomes SPU4 and SPU20. These two sea urchin chromosomes were initially derived from two bilaterian ALGs (BALGs K and O2, respectively) and also correspond to two different jellyfish chromosomes RPE15 and RPE18 (see Supplementary Fig. 14d), supporting the conclusion that PYE3, RPH2, SCO6, PEC12 and SBE2 were all derived from a fused ancestral chromosome in their LCA. These spiralian species also underwent the following lineage-specific chromosomal rearrangement event(s) (e-h and Supplementary Fig. 5). (1) Both RPH14 and SCO9 resulted from A2⊗B2 (e-f). (2) SCO5 and SCO17 are either products of a fused (J1●(Q⊗H)) and subsequently split chromosomes, or they are duplicates of the fused chromosome. The latter scenario is less likely because we did not detect significant conservation between SCO5 and SCO17 (i-j). (3) PYE1 was from M⊗B2. (4) SBE1 resulted from F⊗(C2⊗(L⊗J2)) (h). (4) PEC1 (B1⊗E), PEC4 (J1⊗B2), PEC6 (P⊗D), PEC7 (B3⊗(O1⊗R)) and PEC9 (M⊗A2) were each fused from two annelid ancestral chromosomes (g). (5) SBE2 (J1⊗(O2⊗K)), SBE3 (G⊗M), SBE4 (P⊗N), SBE6 (E⊗(O1⊗R)),

SBE7 (A1⊗B3), SBE8 (D⊗A2) and SBE9 (C1⊗B2) were also each fused from two annelid ancestral chromosomes (h).

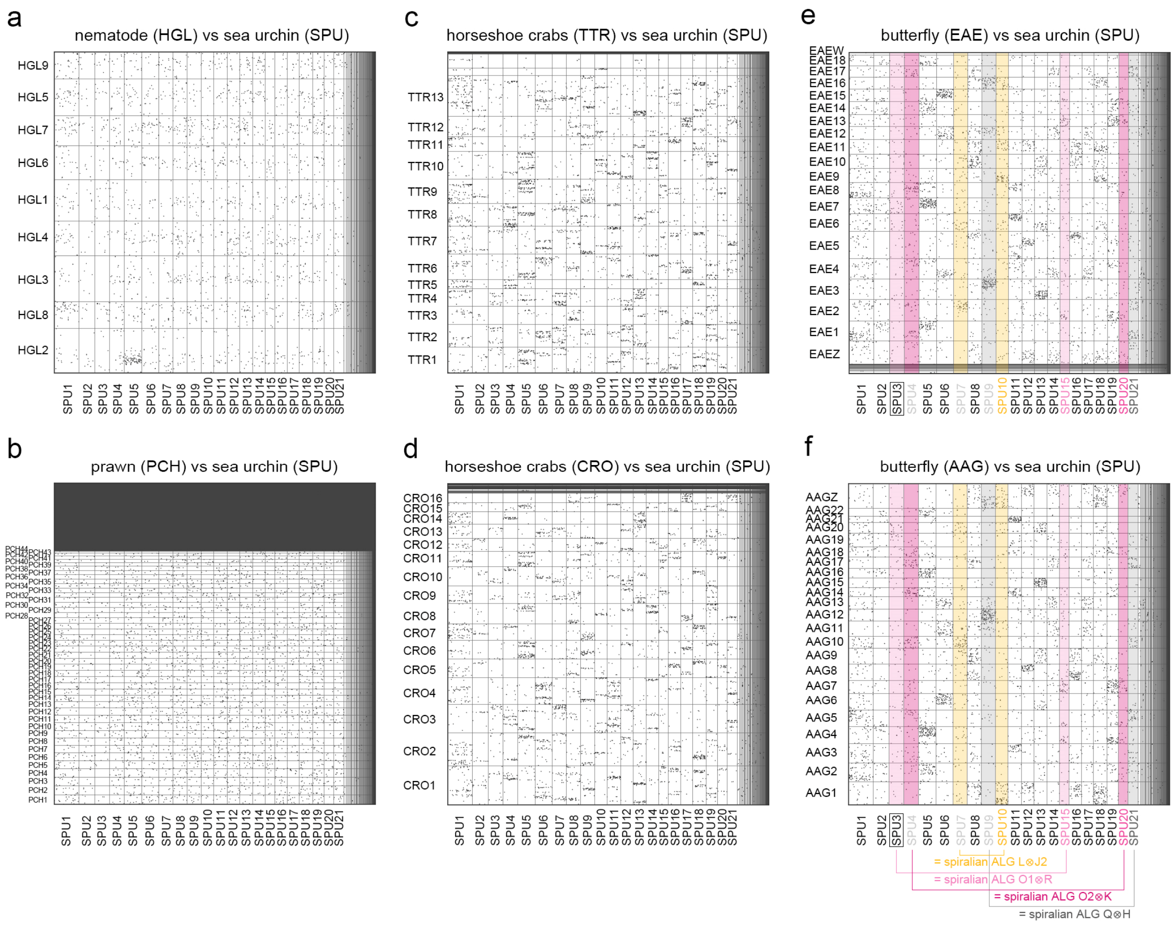


**Supplementary Fig. 14 | Pairwise syntenic dot plots between chromosomes of ecdysozoan species and sea urchin (SPU).**

Pairwise genome comparisons between ecdysozoans and sea urchin SPU showing complex chromosomal rearrangement events in ecdysozoan species, including nematode (a), prawn (b) and horseshoe crabs (c and d). The butterfly genome seems more conserved than the other examined ecdysozoans (e and f). The four spiralian fusion events were not found in butterflies, as sea urchin chromosomes corresponding to fused spiralian chromosomes match to different butterfly chromosomes (indicated by columns of the same color).

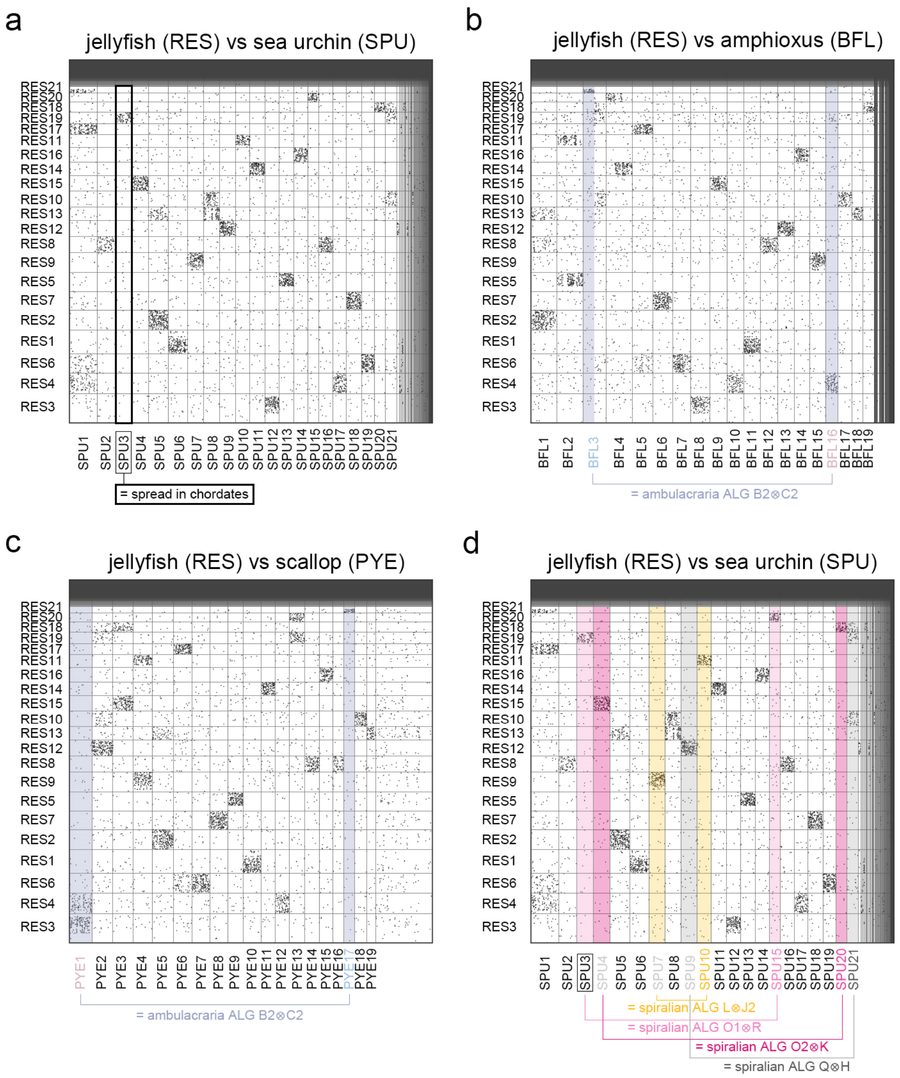


**Supplementary Fig. 15 | Pairwise syntenic dot plots between chromosomes of jellyfish (RES) and bilaterian species.**

The identified chromosomal rearrangement events in bilaterians are not found in the jellyfish genome. SPU3 was derived from DALG R, which dispersed into other chromosomes in chordates. Since SPU3 corresponds to RES19, the chordate dispersal event did not occur in the jellyfish (a). BFL3 and BFL16 correspond to different RES chromosomes, while their ALGs (DALGs B2 and C2) fused into ambulacraria AALG B2⊗C2. Thus, the ambulacrarian fusion event was not found in the jellyfish (b). Similarly, PYE1 and PYE17 both correspond to ambulacraria AALG B2⊗C2 and match to different RES chromosomes (c). The four shared fusion events in spiralians were not found in the jellyfish genome, as sea urchin chromosomes corresponding to fused spiralian chromosomes match to different jellyfish chromosomes (indicated by columns of the same color) (d).

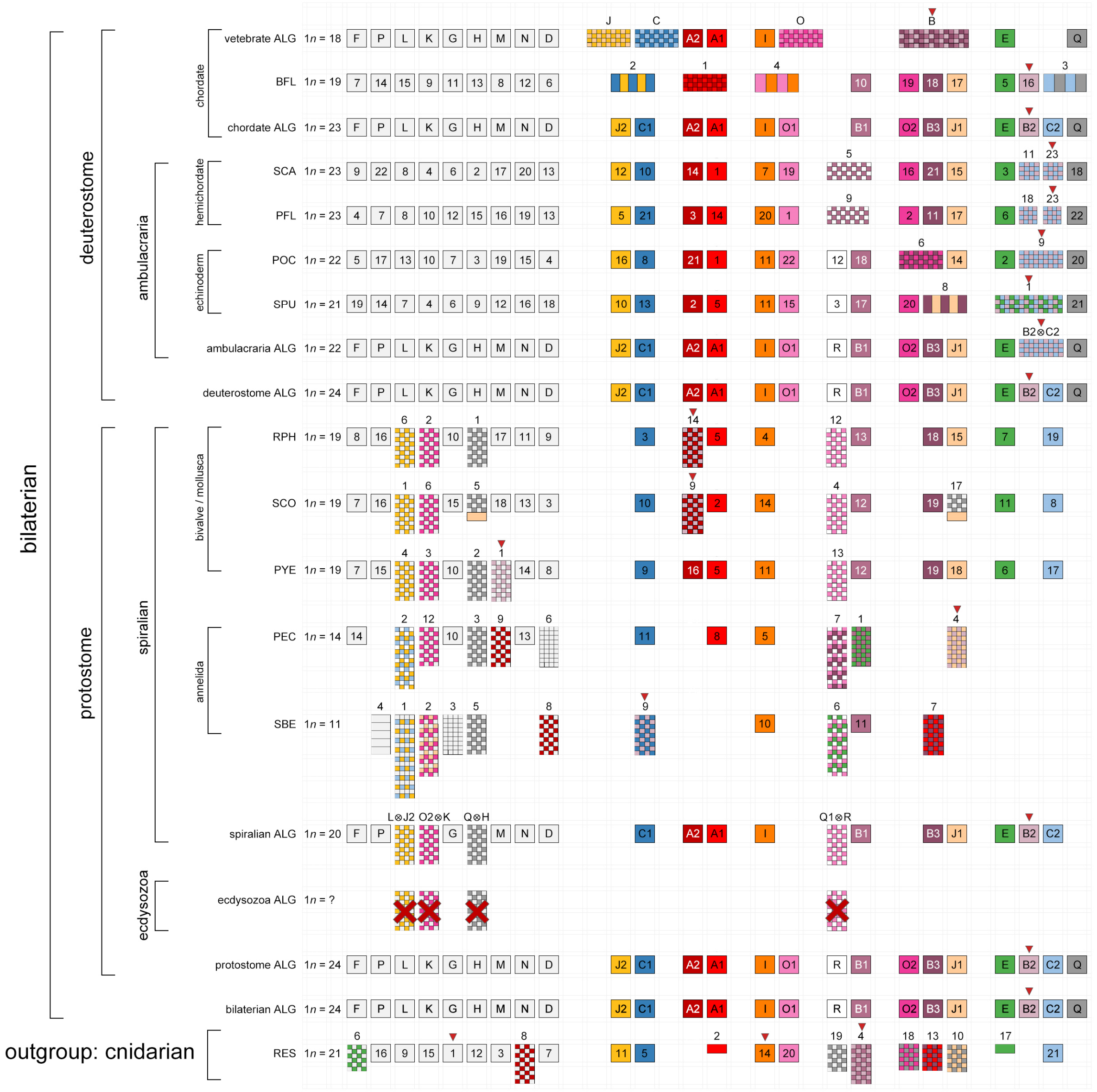


**Supplementary Fig. 16 | Summary of the identified chromosomal rearrangement events.** The genomic architectures of bilaterians and the outgroup jellyfish RES are illustrated. The chromosomal rearrangement events of the jellyfish RES are depicted based on the color codes of the 24 bilaterian ALGs. Red arrowheads indicate Hox cluster-containing chromosomes. Box sizes do not reflect the actual sizes of chromosomes.

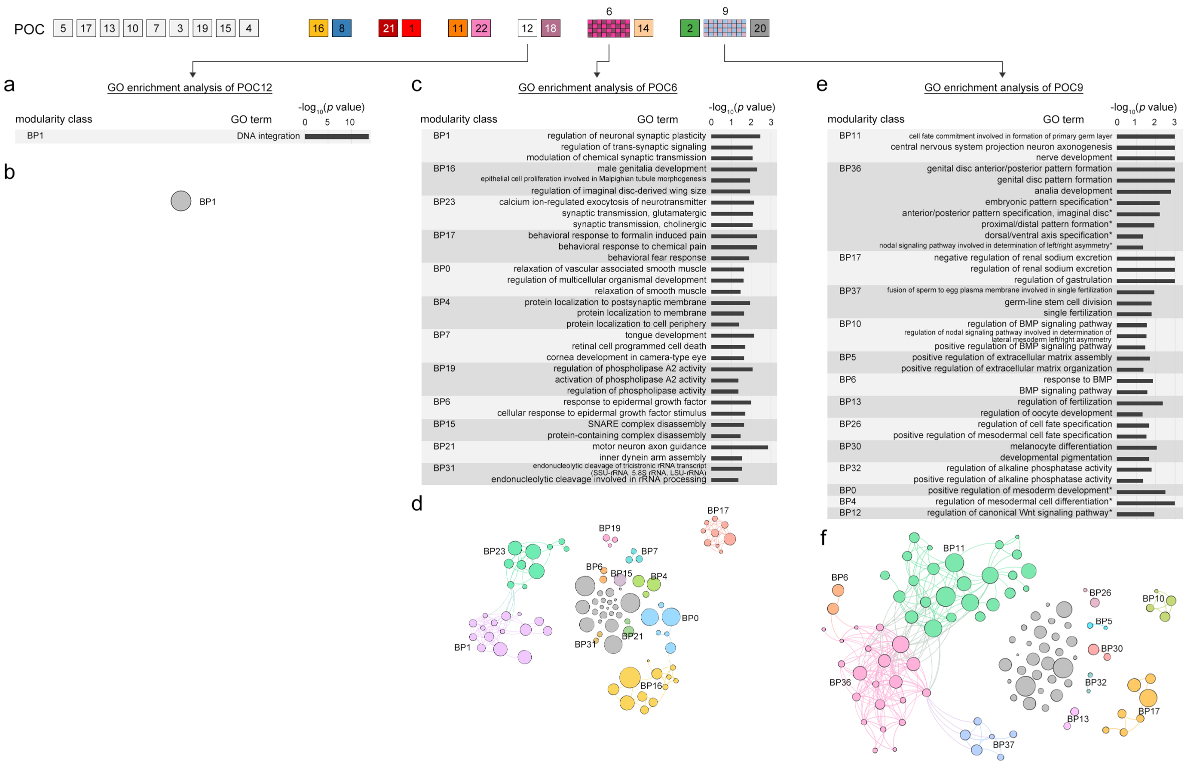


**Supplementary Fig. 17 | Gene ontology (GO) enrichment analyses of the sea star POC chromosomes 12, 6 and 9.**

GO enrichment analyses of genes located on the specific chromosomes of the sea star POC. The enriched GO terms (adjusted *p*-value < 0.1) are clustered and divided into different modules. Descriptions of the most enriched GO terms of biological process (BP) within each module for genes located on POC12 (**a**), POC6 (**c**) and POC9 (**e**). The bars indicate -log10 adjusted *p*-values for the corresponding GO terms. The full list of enriched GO terms, including BP (biological process), CC (cellular component) and MF (molecular function), is provided in Supplementary Data 2. Results of the GO enrichment network analysis of genes located on POC12 (**b**), POC6 (**d**) and POC9 (**f**). Each individual node of the network denotes a specific enriched GO term. Different colors represent different modules of GO terms. Unclassified GO terms are labeled in gray color. Sizes of the circles indicate numbers of genes in each GO term. Manually selected GO terms are indicated with asterisks (*).

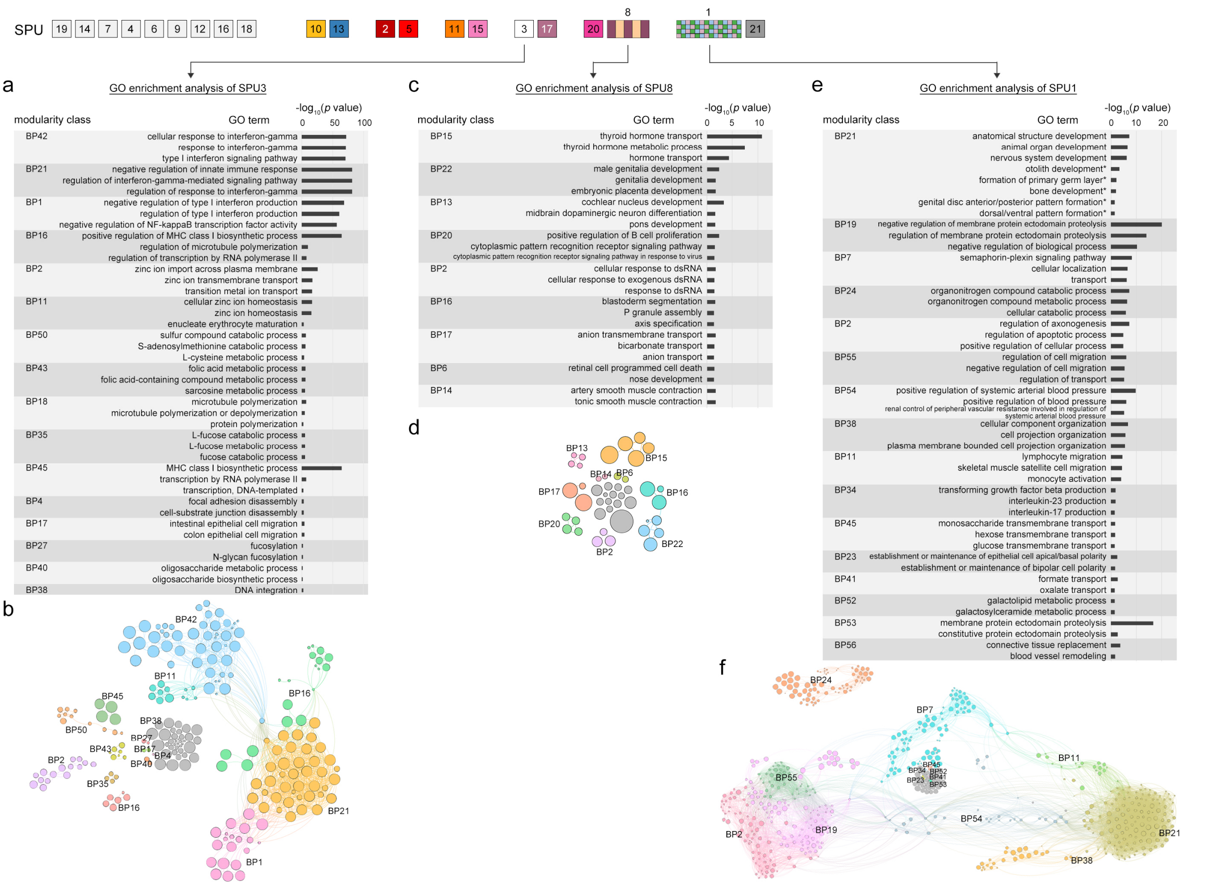

**Supplementary Fig. 18 | GO enrichment analyses of the sea urchin SPU chromosomes 3, 8 and 1.**

GO enrichment analyses of genes located on the specific chromosomes of the sea urchin SPU. The enriched GO terms (adjusted *p*-value < 0.1) are clustered and divided into different modules. Descriptions of the most enriched GO terms of biological process (BP) within each module for genes located on SPU3 (**a**), SPU8 (**c**) and SPU1 (**e**). The full list of enriched GO terms is provided in Supplementary Data 3. Results of the GO enrichment network analysis of genes located on SPU3 (**b**), SPU8 (**d**) and SPU1 (**f**). All labels are consistent with Supplementary Fig. 17.

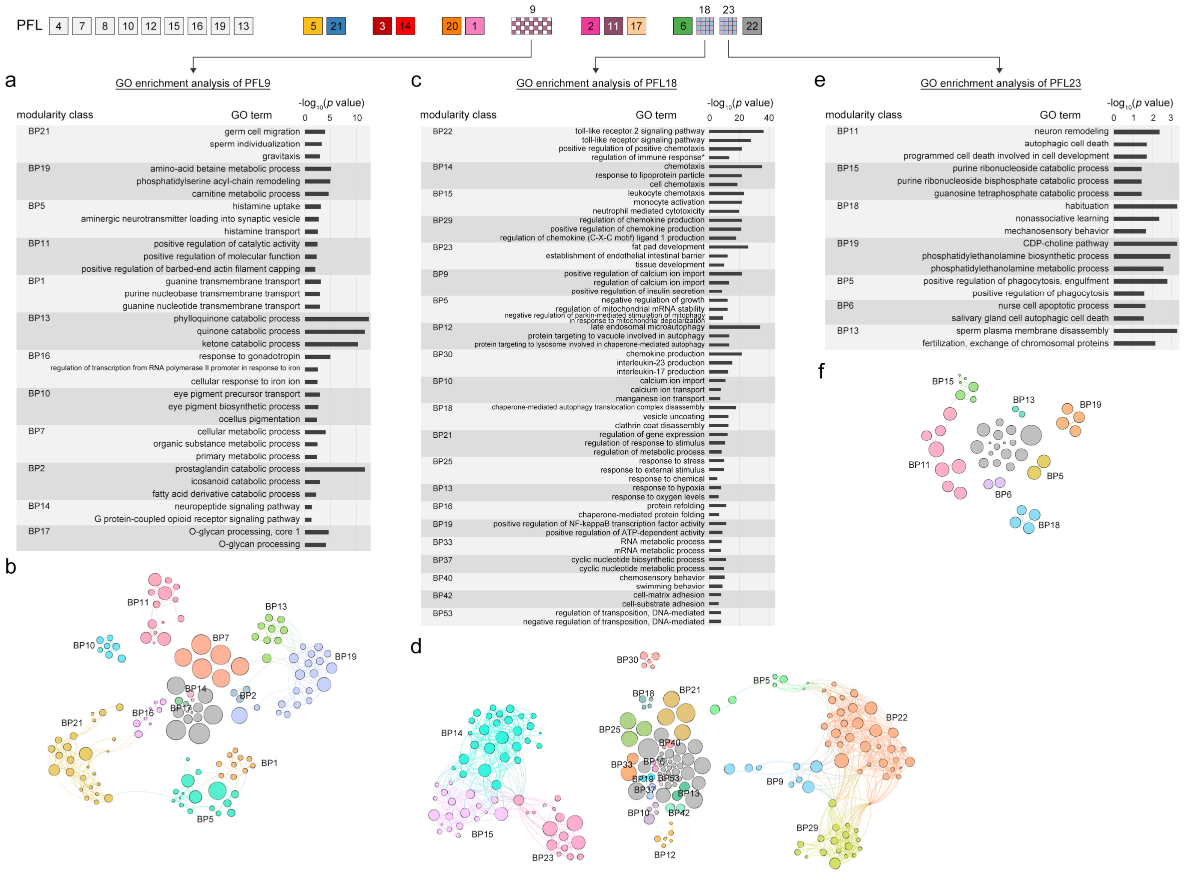


**Supplementary Fig. 19 | GO enrichment analyses of the hemichordate PFL chromosomes 9, 18 and 23.**

GO enrichment analyses of genes located on the specific chromosomes of the hemichordate PFL. The enriched GO terms (adjusted *p*-value < 0.1) are clustered and divided into different modules. Descriptions of the most enriched GO terms of biological process (BP) within each module for genes located on PFL 9 (**a**), PFL18 (**c**) and PFL23 (**e**). The full list of enriched GO terms is provided in Supplementary Data 4. Results of the GO enrichment network analysis of genes located on PFL9 (**b**), PFL18 (**d**) and PFL23 (**f**). All labels are consistent with Supplementary Fig. 17.

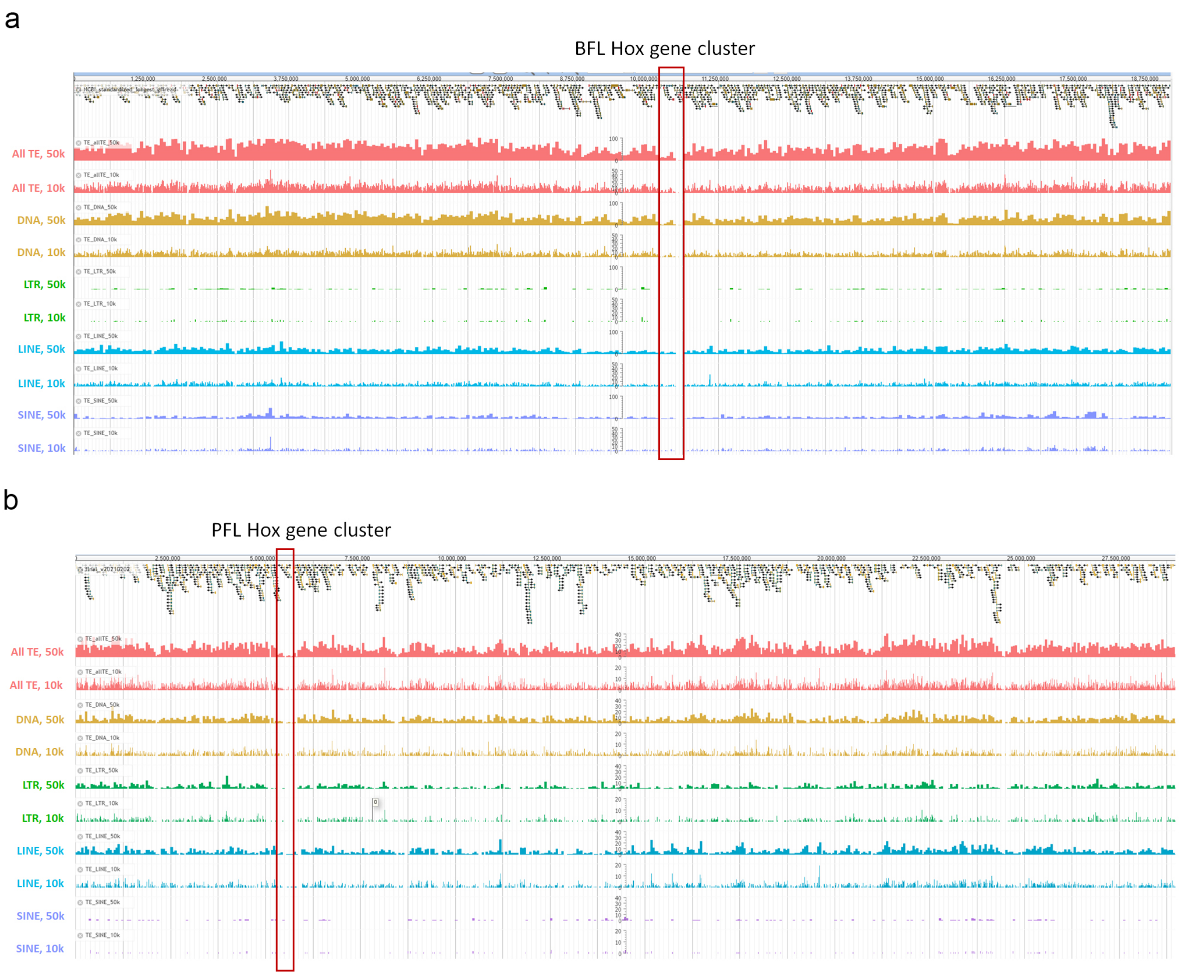


**Supplementary Fig. 20 | Distributions of TEs in amphioxus (BFL) and hemichordate (PFL) Hox-bearing chromosomes.**

The genome browser screenshots of the Hox-located chromosomes of BFL (a) and PFL (b). Histograms of all TEs (red), DNA transposons (DNA, yellow), long terminal repeats (LTR, green), long interspersed nuclear elements (LINE, blue) and short interspersed nuclear elements (SIINE, purple) are shown. The bin size for each histogram of TEs is 50,000 bp or 10,000 bp (indicated on the left). Red boxes denote the genomic regions of the Hox clusters.

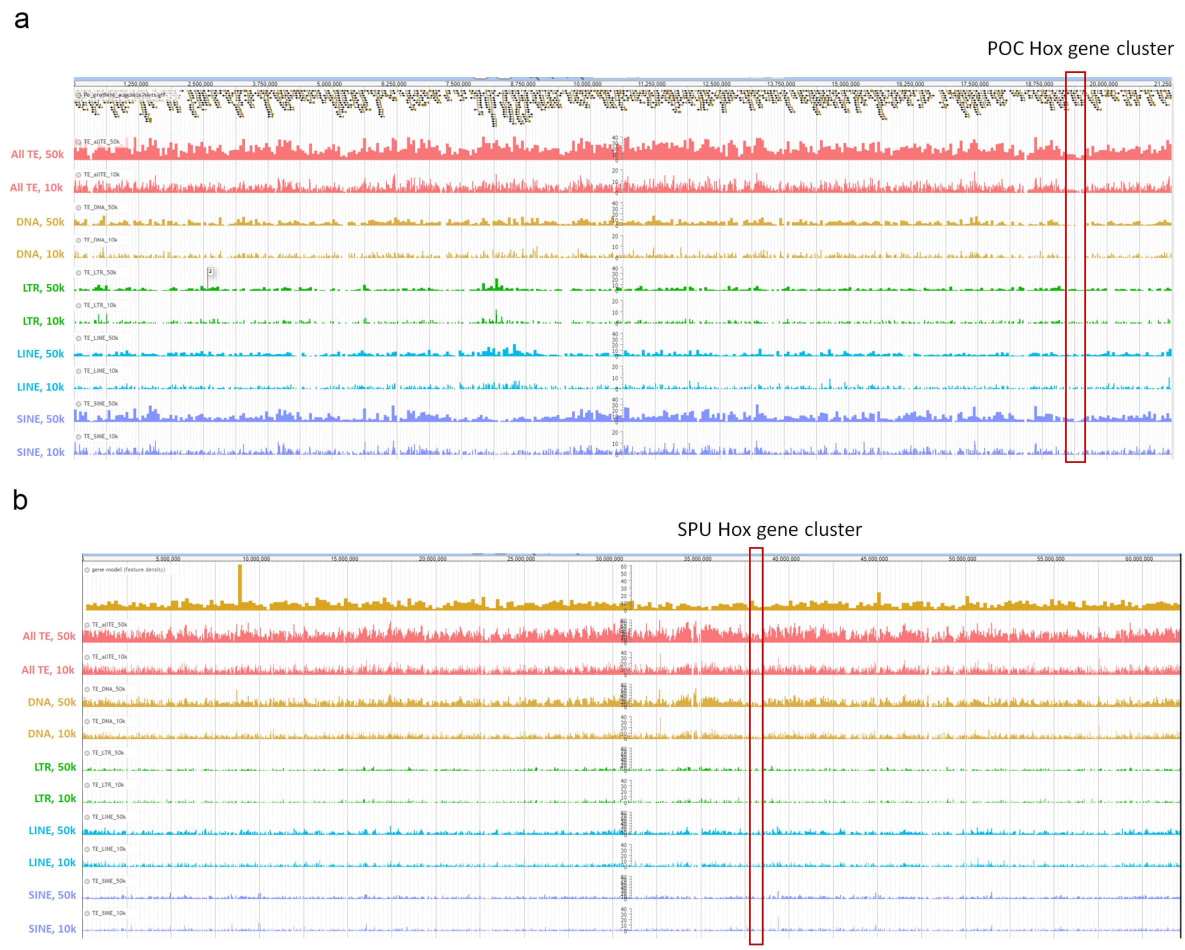


**Supplementary Fig. 21 | Distributions of TEs in sea star (POC) and sea urchin (SPU) Hox-bearing chromosomes.**

Positions of various types of TEs in the Hox-bearing chromosomes of POC (a) and SPU (b). All labels are consistent with Supplementary Fig. 21.

a

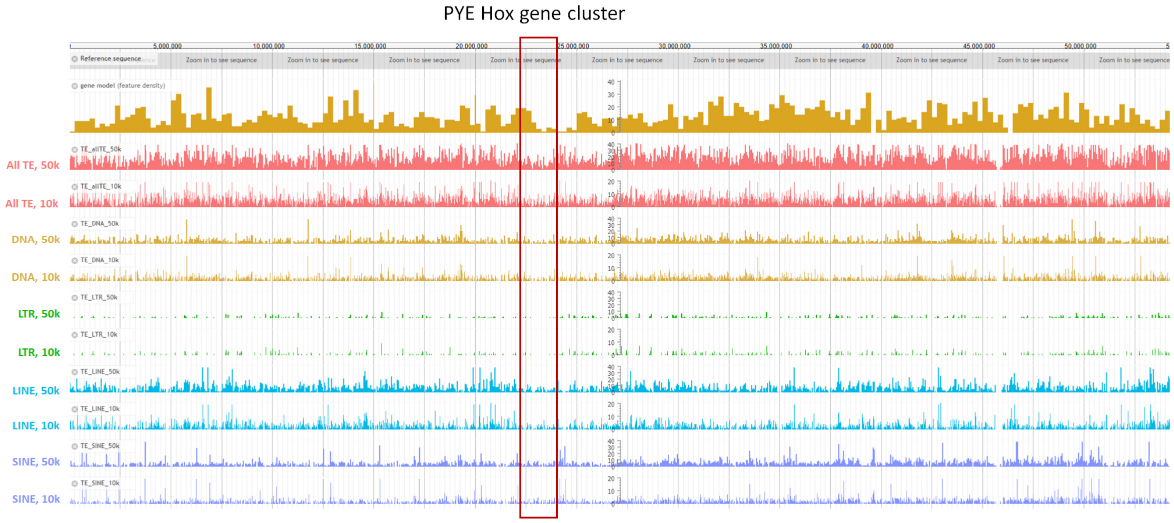


b

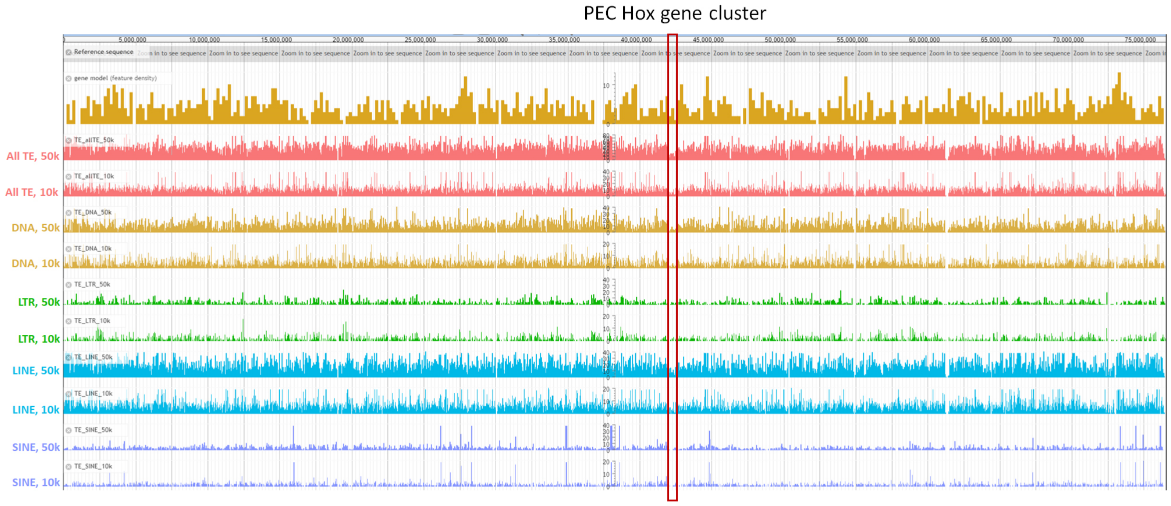


**Supplementary Fig. 22 | Distributions of TEs in scallop (PYE) and annelid (PEC) Hox-bearing chromosomes.**

Positions of TEs in the Hox-bearing chromosomes of PYE (a) and PEC (b). All labels are consistent with Supplementary Fig. 21.

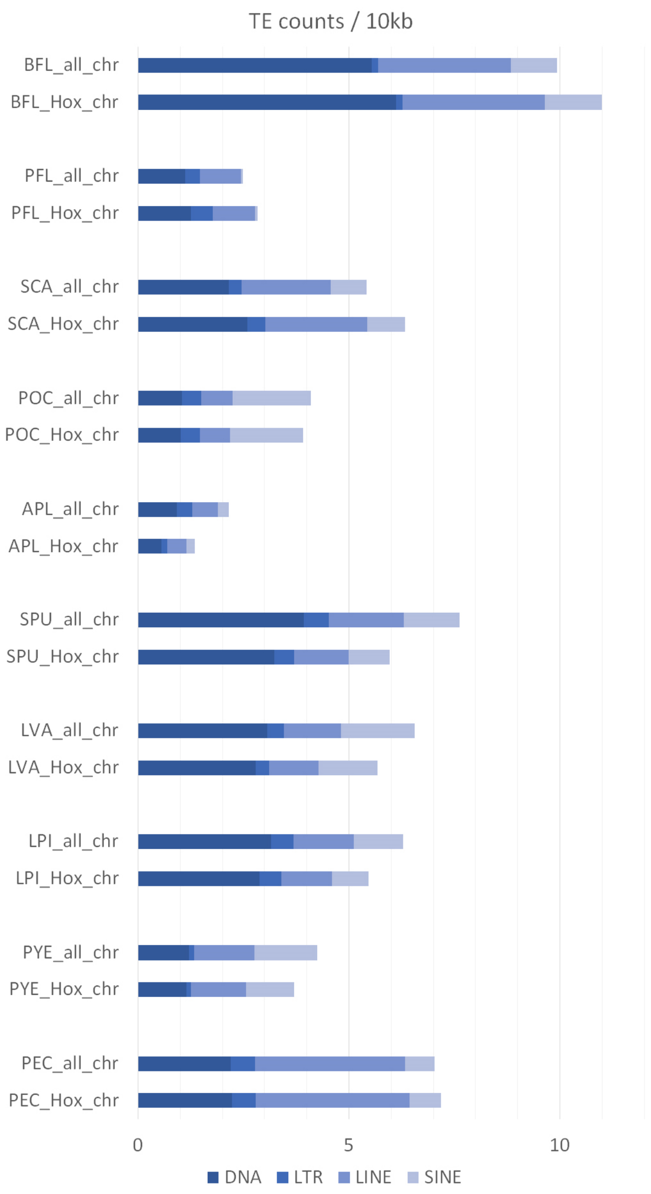


**Supplementary Fig. 23 | TE counts in the ten bilaterian species.**
Numbers of all TEs (DNA + LTR + LINE + SINE) in the whole genome assembly and the Hox-bearing chromosome/scaffold of each species. The TE counts were normalized to a fixed genomic distance (10,000 bp).

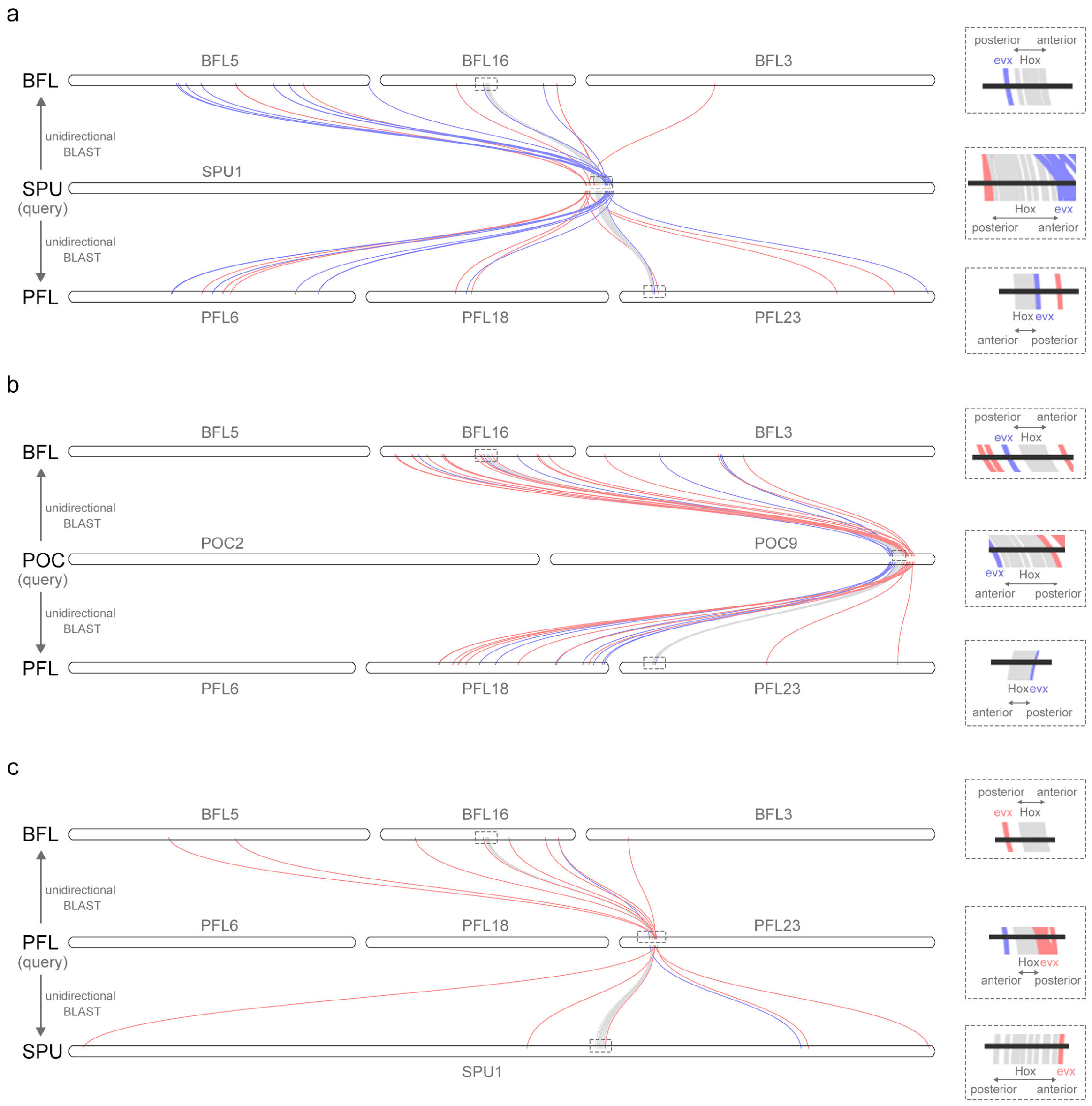


**Supplementary Fig. 24 | Genes neighboring Hox clusters are highly rearranged.** Positional analysis around Hox clusters based on unidirectional BLAST. The query species is shown in the middle of each panel. The curved lines connect gene pairs of the BLAST best hits. Hox genes are labeled in gray. Up to 20 neighboring genes of anterior and posterior Hox genes are shown and labeled in blue and red, respectively. Orthologous genes that are not located in chromosomes descended from DALGs E, B2 and C2 are omitted. The full list of BLAST comparisons is provided in Supplementary Data 5.

a

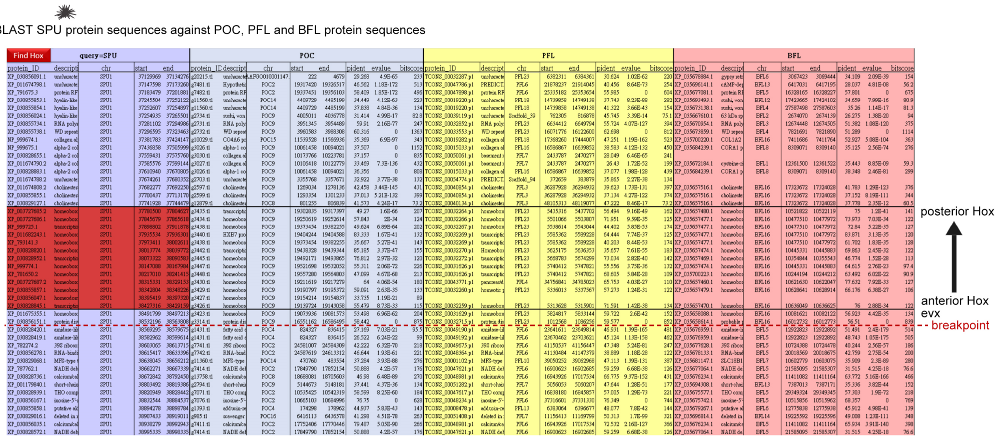
BLAST SPU protein sequences against POC, PFL and BFL protein sequences
Find Hox
query=SPU
POC
PFL
BFL
posterior Hox
anterior Hox
evx
breakpoint

b

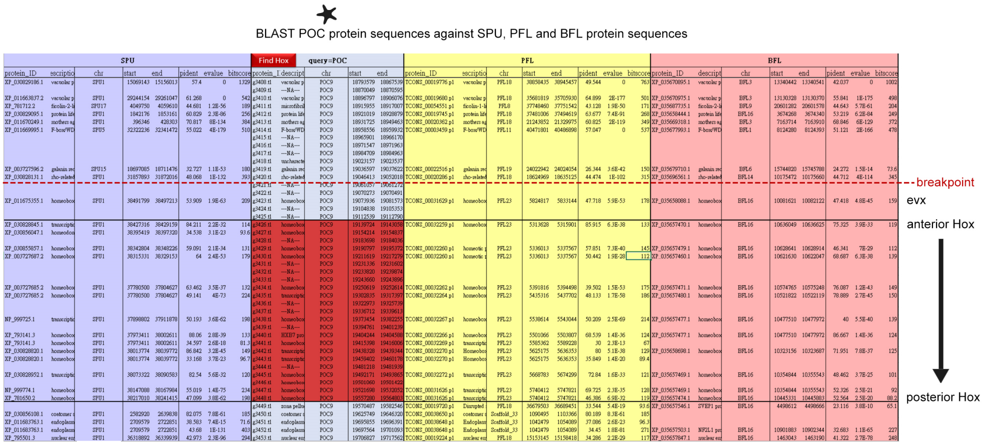
BLAST POC protein sequences against SPU, PFL and BFL protein sequences
SPU
Find Hox
query=POC
PFL
BFL
breakpoint
evx
anterior Hox
posterior Hox

c

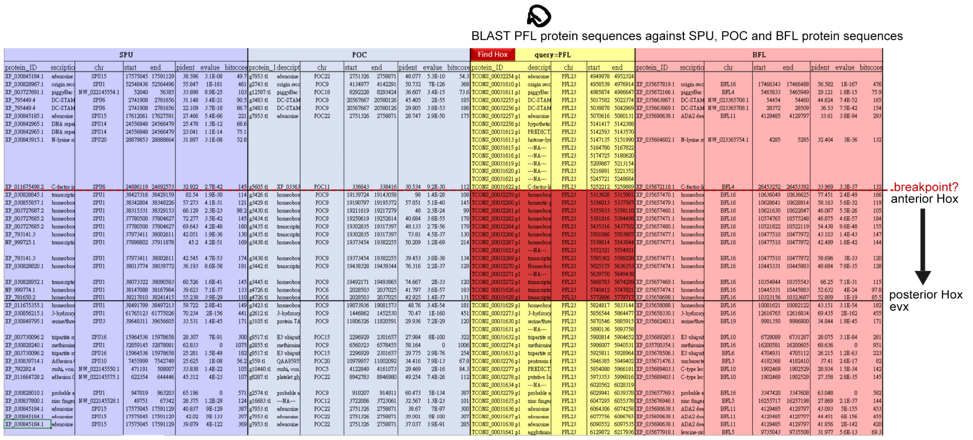
BLAST PFL protein sequences against SPU, POC and BFL protein sequences
SPU
POC
Find Hox
query=PFL
BFL
breakpoint?
anterior Hox
posterior Hox
evx

d

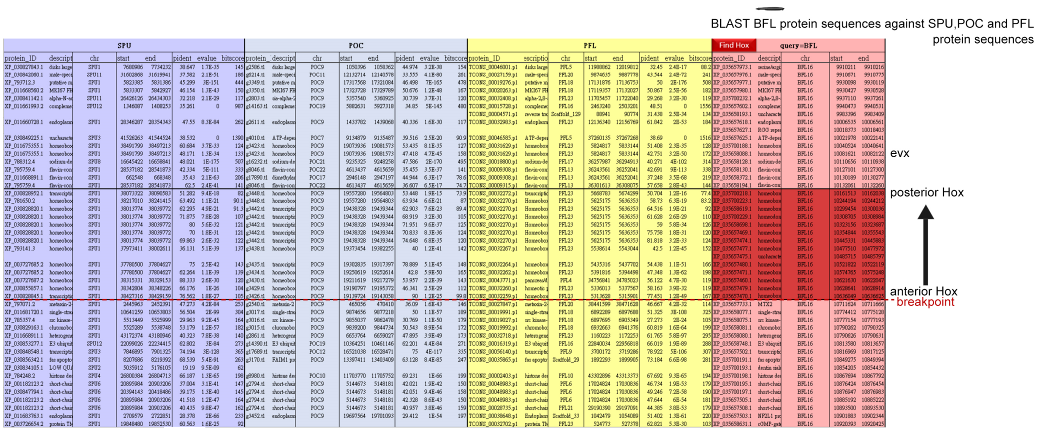
BLAST BFL protein sequences against SPU,POC and PFL protein sequences
SPU
POC
PFL
Find Hox
query=BFL
evx
posterior Hox
anterior Hox
breakpoint

**Supplementary Fig. 25 | Positions of Hox-neighboring genes in deuterostomes.**
Comparing SPU (**a**), POC (**b**), PFL (**c**) and BFL (**d**) protein query to protein databases of other deuterostomes using blastp around HOX gene clusters. Each panel is a screenshot from Supplementary Data 5. The results are sorted by chromosome number followed by the position of the query IDs.

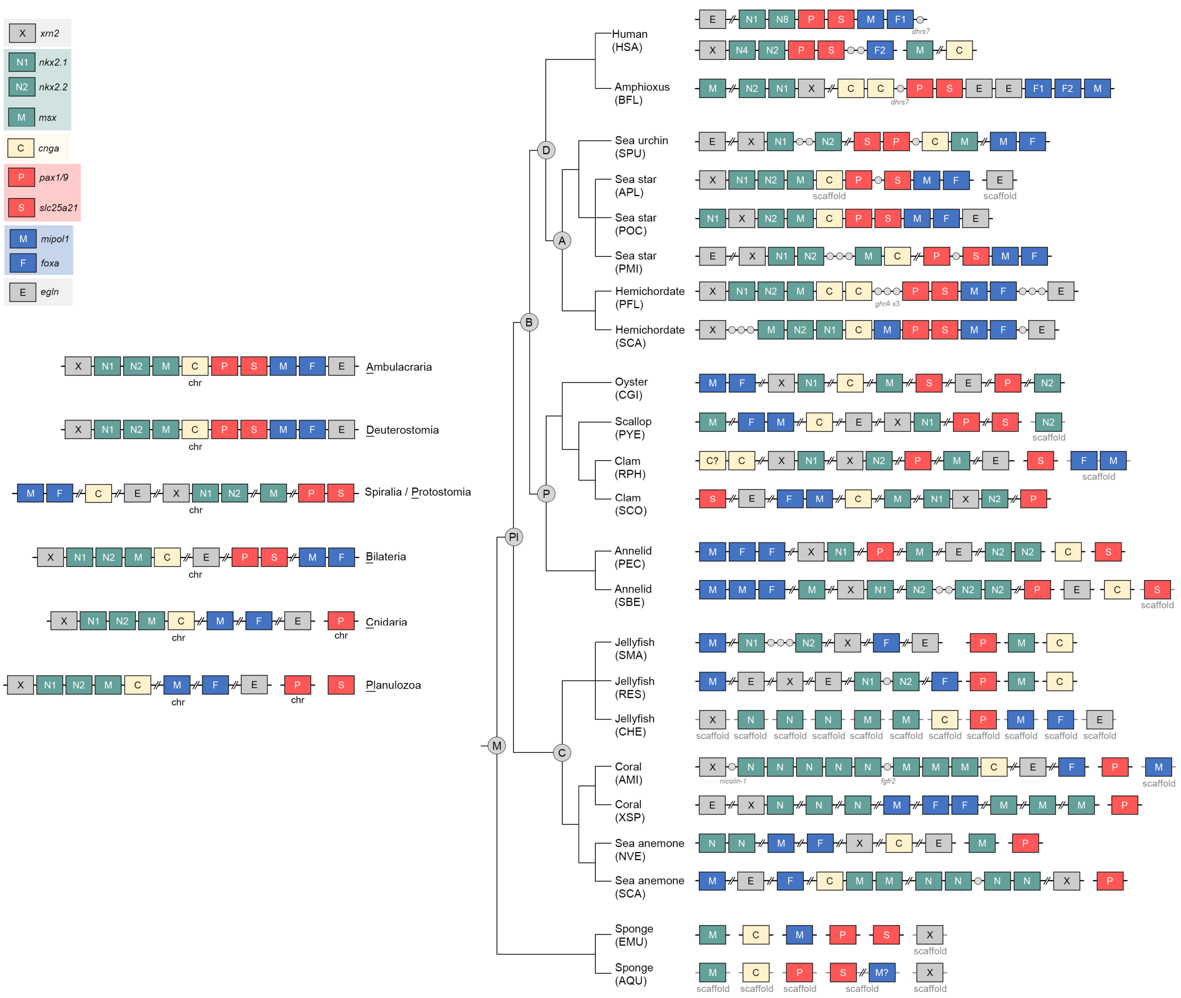


**Supplementary Fig. 26 | Evolutionary history of the pharyngeal gene cluster with the full dataset.** All symbols are consistent with Fig. 6.

### Reference


1 Servant, N. *et al.* HiC-Pro: an optimized and flexible pipeline for Hi-C data processing. *Genome Biology* **16** (2015). https://doi.org:10.1186/s13059-015-0831-x

2 Wang, S. *et al.* Scallop genome provides insights into evolution of bilaterian karyotype and development. *Nat Ecol Evol* **1**, 120 (2017). https://doi.org:10.1038/s41559-017-0120

3 Simakov, O. *et al.* Deeply conserved synteny and the evolution of metazoan chromosomes. *Sci Adv* **8**, eabi5884 (2022). https://doi.org:10.1126/sciadv.abi5884

4 Simakov, O. *et al.* Deeply conserved synteny resolves early events in vertebrate evolution. *Nat Ecol Evol* **4**, 820-830 (2020). https://doi.org:10.1038/s41559-020-1156-z
